# Identification of Novel Endogenous Retroviruses in Anura

**DOI:** 10.1101/2024.07.05.601823

**Authors:** Hayley Beth Free, Richard Nichols, Ravinder Kanda

## Abstract

Endogenous retroviruses (ERVs) represent remnants of past retroviral infections and have had a pivotal role in vertebrate evolution, contributing toward the generation of novel genes and regulatory elements. The majority of ERV research to date has had a mammalian focus; Anurans (the order of amphibians consisting of frogs and toads) are comparable in genome size and number of extant species and yet very few ERVs have been described in this order. Under the current classification, alpharetroviruses appear to be avian-specific; here we screened 47 publicly available Anuran genomes for alpharetroviruses using a bioinformatic pipeline. Both truncated and full-length ERV elements were identified through this screening process which were represented by multiple loci. Ten families of novel Anuran ERVs (AnERV1-10) with intact viral genes were identified in 11 anuran species, some of which are shared between multiple species. Furthermore, based on conserved domain and phylogenetic analysis, all ten novel ERV families appeared in recombined states. This included two families with alpharetrovirus classified Env proteins, although the majority had epsilonretrovirus classified Pol and gammaretrovirus classified Env proteins. These findings extend the pattern of recombination presented in two other Anuran ERVs and adds to the complex evolutionary history of retroviruses. These results suggest that analysis of other vertebrate groups beyond the mammals would provide new information on the presence and state of ERVs, providing further evidence of the modes of ERV evolution within host genomes and their role in the diversification of the vertebrate tree of life.

## 1. Introduction

Endogenous retroviruses (ERVs) are found in the genomes of all vertebrates and represent remnants of past retroviral infections, with many ERVs appearing as ancient insertions without closely related exogenous forms (Boeke & Stoye, 1997; Chu et al., 2023; Herniou et al., 1998; Wang & Han, 2021). Endogenization of a retrovirus can occur from germline integration of the double stranded copy of viral RNA, an obligatory process for retroviral replication which usually occurs in somatic cells (Katzourakis & Gifford, 2010). Although ERV insertions are typically lost through genetic drift, a substantial number have reached fixation. In humans and mice for example, ERVs constitute approximately 10% of the genome (Lander et al., 2001; Waterston et al., 2002). Within host genomes ERVs have contributed toward novel genes and regulatory elements through gene co-option and consequently it has been proposed that they play a pivotal role in vertebrate evolution (Dupressoir et al., 2012; Katzourakis et al., 2005; Pastuzyn et al., 2018).

Full-length retroviruses (approximately 10,000 nucleotides) are comprised of three core genes: *gag* (internal proteins forming matrix and capsid), *pol* (protease, reverse transcriptase and integrase) and *env* (envelope protein), flanked by two long terminal repeats (LTRs) which contain regulatory sequences (Coffin, 1992). The phylogenetic relationship of ERVs is usually determined by their reverse transcriptase (RT) motif in Pol due to the high level of sequence conservation as a key enzyme in the retrovirus replication cycle, which allows the sequences to be reliably aligned and compared. This approach has grouped ERVs with related exogenous virus counterparts (R. Gifford & Tristem, 2003; Jern et al., 2005). These analyses identify three ERV clades: class I consisting of retroviral genera gammaretrovirus and epsilonretrovirus; class II made up of alpharetrovirus, betaretrovirus, delatretrovirus and lentivirus genera; and class III of spumaretrovirus and ancient *env*-less insertion ERV-L (**Figure 1**; Gifford et al., 2018; Gifford & Tristem, 2003). There appears to be a degree of host specificity demonstrated, for example alpharetrovirus have been primarily identified in avian genomes and betaretrovirus in mammalian genomes (Gifford et al., 2005; Henzy & Coffin, 2013; Herniou et al., 1998; Zheng et al., 2022).

**Figure 1:**
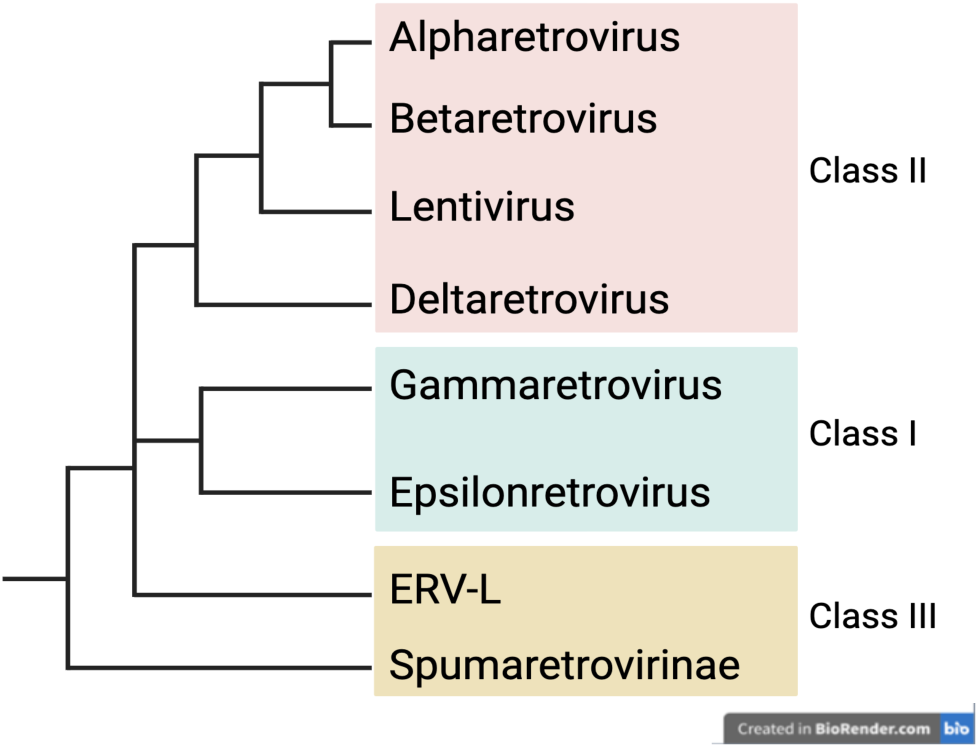
Endogenous and exogenous retrovirus taxonomy and naming system. The family Retroviridae contains seven genera; these are grouped into clades categorising ERVs (class I, II and III), which is based on the phylogeny of reverse transcriptase genes. Adapted from Gifford et al., 2018.

The majority of ERV research to date has focused on mammals; Anurans represent a similar number of extant species with comparable genome sizes, yet very few ERVs have been described in this order (frog and toad species). To date only 55 unique ERVs have been identified in 18 anuran species across three retroviral genera (gammaretrovirus, epsilonretrovirus and spumaretrovirus; Aiewsakun & Katzourakis, 2017; Y. Chen et al., 2021, 2022; Herniou et al., 1998; Kambol et al., 2003; Martin et al., 1999; Russo et al., 2018; Sinzelle et al., 2011; Tristem et al., 1996; Yedavalli et al., 2021). Here we describe the screening of 47 publicly available Anuran genomes (15 toad species and 32 frog species) with representative alpharetroviruses; this approach identified truncated and full-length elements, represented by multiple loci within and between genomes. Ten families of novel ERVs (multiple loci within a genome) were identified in 11 anuran species.

## 2. Methods

An outline of the methods can be seen in **Figure 2**.

**Figure 2:**
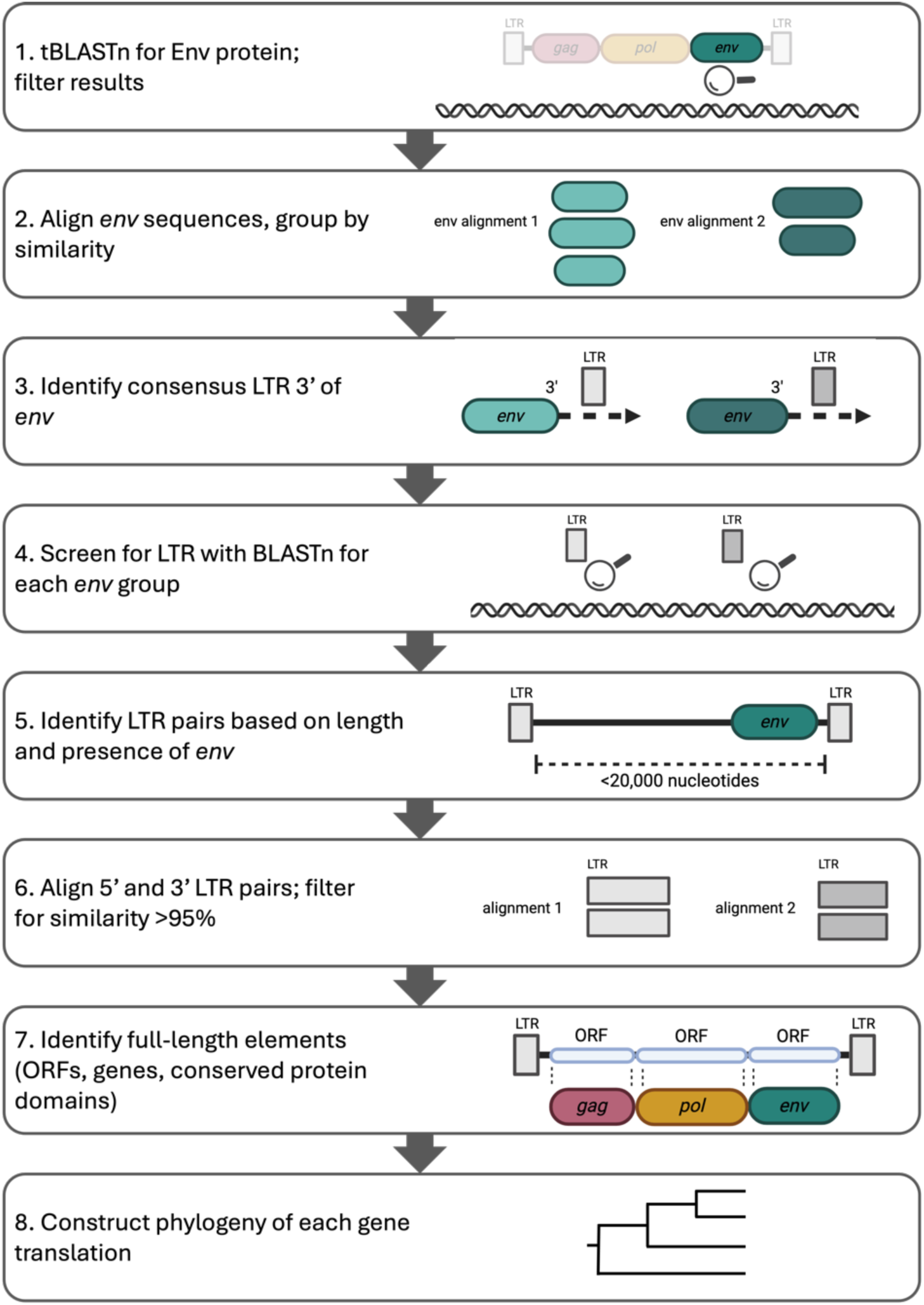
Overview of bioinformatic approach to identify novel ERVs.

### 2.1. Initial tBLASTn Screening of Anuran Genomes Using Env

To identify alpha- ERVs the 47 publicly available genomes (15 toad species and 32 frog species; **Table S1**) were initially screened using tBLASTn (default command line parameters; Altschul et al., 1990), against amino acid sequences of Env proteins from four representative alpharetroviruses: Avian leukemia virus (NC_015116.1); Avian leukosis virus (KU375453.1); Rous sarcoma virus (J02342.1); Avian endogenous retrovirus EAV-HP (AJ238121.1). The results were subsequently filtered based on length (>30% of representative total Env) and an E value of <1e^-5^; multiple tBLASTn hits within 2,000 nucleotides in the same chromosome, comprising different parts of the Env were considered to be a single locus and concatenated. The representative retroviruses used as queries in the tBLASTn have some sequence similarity in Env sequence, resulting in overlapping hits which were removed. The loci identified by these criteria were extracted from the genome and the sequences were grouped by similarity using a pairwise BLAST (percentage identity and query coverage >60%).

### 2.2. LTR Identification

In each *env* group found through pairwise BLAST, an additional 1,000 nucleotides were added to the 3’ of each sequence to identify the LTR sequence. These were aligned using MUSCLE and manually inspected in Geneious. The *env* open reading frame (ORF) was identified using NCBI ORFfinder from the consensus sequence of the alignment, and the 3’ flanking region from the stop codon of *env* was estimated as to where an LTR would be located if there was alignment throughout the sequences (**Figure 2**, step 3). The consensus LTR was identified for all *env* alignments, and then using MUSCLE pairwise alignment in Geneious, the percentage identity was used to classify similar LTRs between different env groups (LTRs showing at least 50% identity).

### 2.3. Full-Length Element Identification

Once the consensus LTRs had been identified they were used to screen the genomes using BLASTn (default parameters, with exception of word size as 10). The results were filtered (>30% of representative LTR length, E value <1e-5); multiple blast hits in the vicinity of 2,000 nucleotides in the same chromosome were considered to be the same locus and concatenated. To identify full-length ERV elements, the LTR BLAST results were used alongside the *env* results, to identify LTR sequences which are both within 20,000 nucleotides of another LTR, and *env* is present between them (**Figure 2**, step 5). These LTR sequences were then extracted, and the LTR pairs aligned with MUSCLE and/or manually in Geneious. The distance matrix of the LTR alignment from Geneious was used to identify LTR pairs which have >95% identity; pairs of LTRs which have a higher similarity are more likely to be recent full-length insertions as the LTRs are identical upon integration (Belshaw et al., 2005). These LTR pairs (>95% similarity) were extracted and aligned as single full-length elements: from the start of 5’ LTR to end of 3’ LTR, with sites containing 25% gaps masked. For each species genome, the consensus from each potential full-length ERV alignment were compared between the *env* groups and were merged if there was >80% identity across the full-length ERV. The consensus sequence of the alignments was used in ORFfinder and BLASTx to Retroviridae and Viruses databases (taxid ID 11632 and 10239 respectively) to identify internal genes. The full-length alignments were then filtered for having ORFs coinciding with the three internal genes (**Figure 2**, step 7). The translated *env* gene of these elements were then compared between species genomes, using BLASTp, and considered the same insertion if there was >50% identity and >80% query coverage. The full-length alignments which appeared as the same insertion between species were then also merged. The full-length alignments which had three ORFs coinciding with *gag, pol* and *env* >1000 nucleotides were analysed further.

### 2.4. Viral Genome Annotation

Conserved protein domains were assessed by translating the individual genes and using conserved domain database to identify key domains: zinc fingers in Gag, protease, reverse transcriptase, RNaseH, integrase and dUTPase in Pol, and heptad repeats and immunosuppressive domain (ISD) in Env TM (Wang *et al*., 2023). Myristylation signals on Gag were identified with SUPLpredictor (mendel.imp.ac.at/myristate/SUPLpredictor) and manually inspected. The remaining conserved domains were identified through manual inspection (see results for list of conserved domains assessed).

### 2.5. Phylogenetic Analysis

To classify the insertions in addition to analysing the conserved domains of the proteins, the evolutionary relationship of the full-length ERV insertions were investigated through phylogenetic analysis of each protein. The amino acid sequences of the ERV groups for the RT of the Pol and the TM subunit of the Env were aligned with representative retroviruses from each genera using MUSCLE and manually adjusted where necessary in Geneious (**Table S2**). The model for Pol and Env was LG with four gamma categories and estimated proportion of invariable sites, a random local clock and 100,000,000 chain length and 10,00,000 burn-in. The trees were visualised and annotated with iTOL (Letunic and Bork, 2024). Epsilonretrovirus, spumaretrovirus and lentivirus were not used in the Env TM phylogeny due to poor quality alignment. Phylogenetic analysis of the Gag protein was also undertaken, however due to high amounts of variability between the full-length ERV insertions identified in this study with representative retroviruses, these were undertaken for each retroviral genus with the same parameters as the Env and Pol trees.

## 3. Results

### 3.1. Screening for Env Protein

All 47 genomes were screened with Env protein sequences from four representative alpharetroviruses; 7/47 genomes did not have any tBLASTn results and a further four did not have any results after filtering for length and E value. Of the 36 genomes with *env* blast results, the *env* sequences were grouped based on their similarity, resulting in multiple *env* families in each genome screened. This ranged from one to five *env* families.

### 3.2. Identifying LTRs

Extracting an extra 1000 nucleotides from the 3’ end of the env enabled us to identify the LTR. The *env* blast results were used to identify the flanking LTR sequences. Assumed LTR sequences flanking the env were identified in 35 genomes. The consensus LTR sequences were extracted for each *env* group and used to screen back against the genomes with BLASTn and filtered for quality. All genomes had BLASTn results from screening with consensus LTR sequences.

### 3.3. Identifying Full-Length Elements

Upon identifying the LTR sequence, then extracting an extra 1000 nucleotides from the 3’ end of the *env*, and screening for the consensus LTRs against the genomes, we could identify full-length ERV elements. The databases of *env* and LTR sequences was used to identify potential full-length elements, based on the distance between LTRs and presence of *env*. Potential full-length elements were identified in 28 genomes, and the 5’ and 3’ flanking LTRs of these elements were extracted and aligned. The LTR pairs which had >95% identity were identified in 25 of these genomes, which were extracted as full-length elements (start of 5’ LTR to end of 3’ LTR) to identify internal genes. The potential full-length elements were aligned and sites containing 25% gaps were masked. There were between 1-13 full-length ERV groups from 24 genomes (**Table S3**). The ORFs were assessed for their length and position coinciding with the *gag, pol* and *env* genes. There were 29 full-length ERV families in 13 species which did not have ORFs corresponding to one or more internal genes. In four species, six of these families appear to represent segmental duplication when 1000 nucleotides were added on both the 5’ and 3’ showed further alignment. Of the insertions which had BLASTx hits to three internal genes, 44 ERV families from 15 species had ORFs which were either <1000 nucleotides or were fragmented. Insertions with ORFs coinciding with the three genes, and an *env* ORF >1000 nucleotides were compared between genomes. This consisted of 31 families of full-length ERV insertions from 19 Anurans (seven frog and 12 toad species). The *env* sequence was translated from the consensus of the alignment and compared between genomes, merging ERV groups with >80% query coverage and >50% identity of the Env. This identified 26 families of ERV insertions present across the 19 Anuran species. To annotate these full-length insertions further, those which contained whole ORFs for *gag* and *pol* (>1000 nucleotides) were used. Of these, there were 10 groups of full-length ERV insertions in 11 Anuran species (five frogs and six toads; **Table 1**, **Figure S1**). These insertions were named AnERV1 (**An**uran **e**ndogenous **r**etro**v**irus and the numerical value of the group). AnERV2, AnERV4 and AnERV5 were shared between multiple species, where AnERV2 and AnERV4 were shared between species of the same genus (*Spea* and *Bufo* respectively), and AnERV5 was shared between species which have almost 40 million years of divergence (Kumar et al., 2017).There are also Anuran species which contain multiple novel ERV elements: *Spea multiplicata* had AnERV1 and AnERV2, *Bombina bombina* had AnERV7 and AnERV8, and *Gastrophryne carolinensis* had AnERV6 and AnERV9.

**Table 1:** Novel full-length ERV insertions identified in 10 species. Years of Anuran divergence calculated in TimeTree.org.

| ERV ID | Species | Genome size (Gb) | Number of solo LTRs | Number full-length ERV loci (LTRs >95% identity) | Years of Anuran divergence (those with shared insertions) |
| --- | --- | --- | --- | --- | --- |
| AnERV1 | *Spea multiplicata* | 1.8 | 142 | 1 |  |
| AnERV2 | *Spea multiplicata* | 1.8 | 49 | 1 | 8-19 million |
|  | *Spea bombifrons* | 1 | 120 | 5 |  |
|  | *Spea hammondi* | 1.2 | 195 | 7 |  |
| AnERV3 | *Pelobates cultripes* | 3.1 | 1428 | 17 |  |
| AnERV4 | *Bufo bufo* | 5 | 348 | 2 | 14 million |
|  | *Bufo gargarizans* | 4.5 | 315 | 26 |  |
| AnERV5 | *Lithobates sylvaticus* | 5.2 | 115 | 1 | 39 million |
|  | *Glandirana rugosa* | 7.6 | 754 | 1 |  |
| AnERV6 | *Gastrophryne carolinensis* | 4.3 | 312 | 23 |  |
| AnERV7 | *Bombina bombina* | 10 | 2722 | 23 |  |
| AnERV8 | *Bombina bombina* | 10 | 449 | 12 |  |
| AnERV9 | *Gastrophryne carolinensis* | 4.3 | 284 | 18 |  |
| AnERV10 | *Xenopus tropicalis* | 1.5 | 169 | 2 |  |

### 3.4. Annotating ERV Insertions

The 10 families of full-length ERV insertions identified across 11 species were annotated by identifying conserved protein domains in the translated genes. This can be useful in identifying the completeness of an insertion and for classification, as the presence and/or location of certain domains can be specific to retroviral genera. A summary of these domains of each insertion and comparison to similar ERVs can be seen in **Figure 3**.

**Figure 3:**
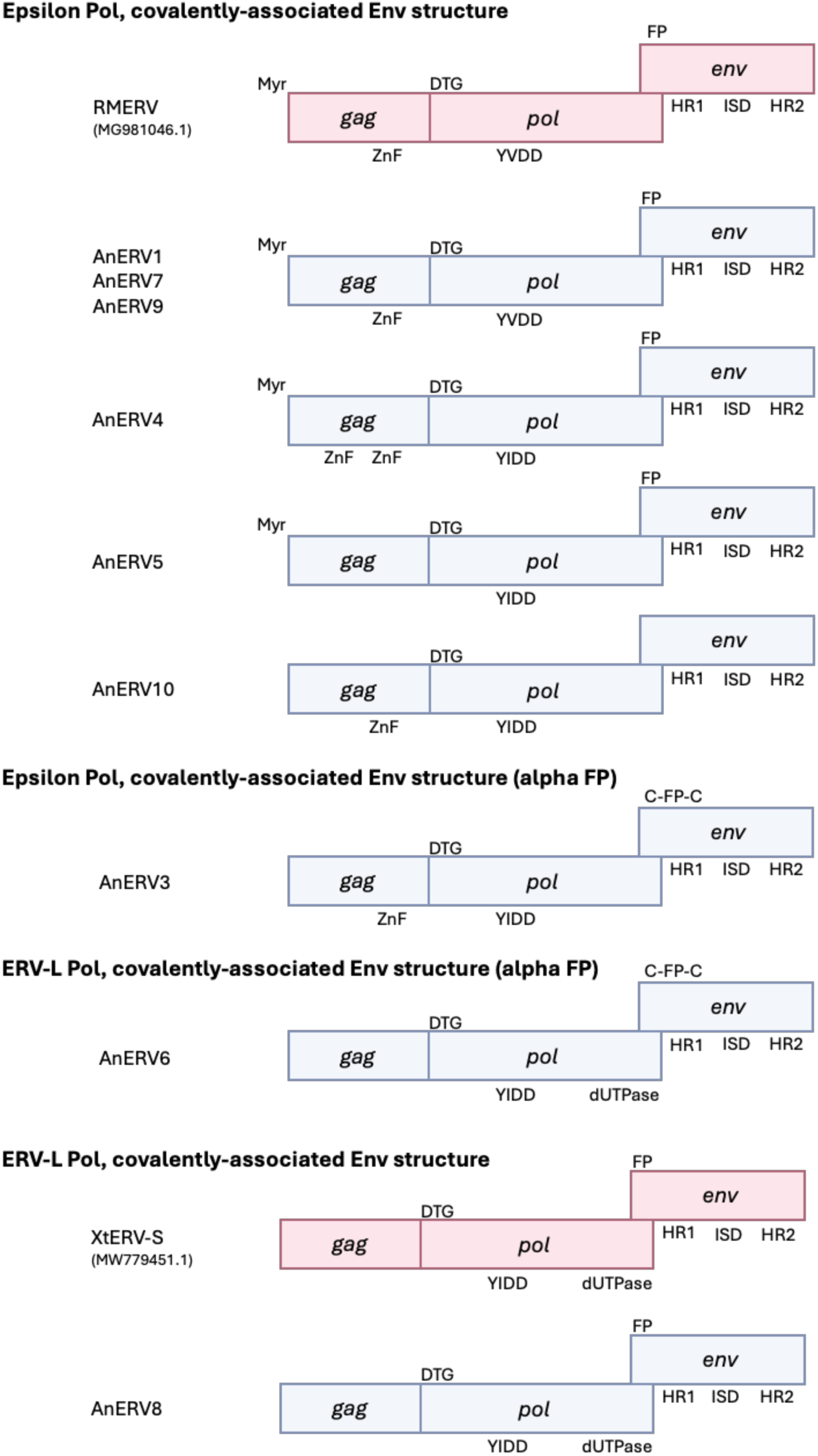
A selection of the conserved domains in the identified full-length ERVs in 11 Anuran species (blue) and compared to previously described ERV insertions (pink) gag, pol and env genes (Russo et al., 2018; Yedavalli et al., 2021). Headings indicate the classification based on the conserved domain and phylogenetic analysis of Pol reverse transcriptase (RT) and Env transmembrane subunit amino acid sequences. AnERV2 was not included due to lack of conserved domains in Gag and Pol, and exclusion from Pol phylogeny due to lack of RT. Myr = Myristoylation signal; ZnF = zinc finger, DTG = catalytic motif; YXDD = catalytic centre of RT; FP = fusion peptide; C-FP-C = cysteine flanked FP; HR = heptad repeat; ISD = immunosuppressive domain.

#### 3.4.1. Gag Conserved Domains

The myristylation signal on the N-terminus of Gag is present in betaretrovirus, gammaretrovirus and lentivirus, which functions to target Gag to the plasma membrane (Gottlinger et al., 1989). Here it is identified in all full-length ERVs excluding AnERV6, AnERV8, AnERV9 and AnERV10 (**Table 2**). The capsid major homology region (MHR) is found in all retroviruses except for spumaretroviruses (Demirov & Freed, 2004; Hütter et al., 2013) and is present here in AnERV1, AnERV4, AnERV6, AnERV7 and AnERV8. Lastly, the zinc finger domain, also called Cys-His motif as it is made up of CX2CX4HX4C, is present in all retroviral genera bar spumaretrovirus and ERV-L, with two copies in alpharetrovirus, betaretrovirus, epsilonretrovirus and some gammaretroviruses, as well as lentivirus (Jern et al., 2005; Llorens et al., 2009; Strack et al., 2003; Von Schwedler et al., 2003). In AnERV1, AnERV3, AnERV5, AnERV7, AnERV9 and AnERV10 the Cys-His zinc finger domain was identified as a single copy, and in AnERV4 with two copies. Although AnERV9 only possessed one copy of the Cys-His zinc finger domain, two other zinc fingers were identified in Gag and classified as RanBP2-type. These are typically found in eukaryotic proteins, specifically as components of nuclear pore complexes (Kamei et al., 1998; Nguyen et al., 2011; Wu et al., 1995; Yokoyama et al., 1995).

**Table 2:** Conserved domains identified in Gag of the 10 AnERVs (labelled 1-10) identified in 11 species and their inclusion in different retroviral genera.

| Feature | Retroviral genera with domain | 1 | 2 | 3 | 4 | 5 | 6 | 7 | 8 | 9 | 10 |
| --- | --- | --- | --- | --- | --- | --- | --- | --- | --- | --- | --- |
| Myristylation signal | Beta, gamma, lenti | ✓ | ✓ | ✓ | ✓ | ✓ | x | ✓ | x | x | x |
| Capsid major homology region | All but spuma | ✓ | x | ✓ | x | x | ✓ | ✓ | ✓ | ✓ | ✓ |
| Zinc finger / Cys-His motif | One-two copies in all but spuma | ✓ | x | ✓ | ✓ (x2) | ✓ | x | ✓ | x | ✓ | ✓ |

#### 3.4.2. Pro/Pol Conserved Domains

In all full-length AnERVs which contained Pro, Gag was in the same reading frame as Pro and Pol, and the catalytic motif was present as DTG, which is typically identified in betaretroviruses, gammaretroviruses and ERV-L (**Table 3**; Chameettachal et al., 2023; Konvalinka et al., 2015; Wlodawer & Gustchina, 2000). In AnERV2, Pro was not identified and there was also an absence of the catalytic centre in the RT of Pol, which implies that there may be low sequence quality of Pro/Pol in AnERV2. Nonetheless, the zinc finger motif present in the integrase of Pol was identified in AnERV2, as well as the remaining nine groups. The catalytic centre of RT of the remaining groups is present but shows variability between YVDD and YIDD motifs. The YIDD motif is found in MuERV-L, human ERV-K, a novel recombined ERV in *Xenopus tropicalis*, and a variant of HIV-1 (Bénit et al., 1997; Contreras-Galindo et al., 2017; Sharma et al., 2005; Yedavalli et al., 2021). The YVDD motif is found in gammaretroviruses and spumaretroviruses (Donahue et al., 1988; Harrison et al., 2021; Katzourakis et Poch et al., 1989). AnERV6 and AnERV8 at the 3’-terminus of Pol was a dUTPase motif. This domain is typically found in betaretroviruses, non-primate lentiviruses and ERV-L, but the genomic location of the dUTPase identified here is only similar to that of the ERV-L structure (Barabás et al., 2003; Bénit et al., 1997, 1999; Bergman et al., 1994; Coffin et al., 1997; Hizi et al., 1987, 1989). This location of dUTPase in Pol has also been identified in a novel ERV found in the *X. tropicalis* genome, which is a recombination of class III ERVs with a Env structure similar to that of gammaretrovirus (Yedavalli et al., 2021).

**Table 3:**
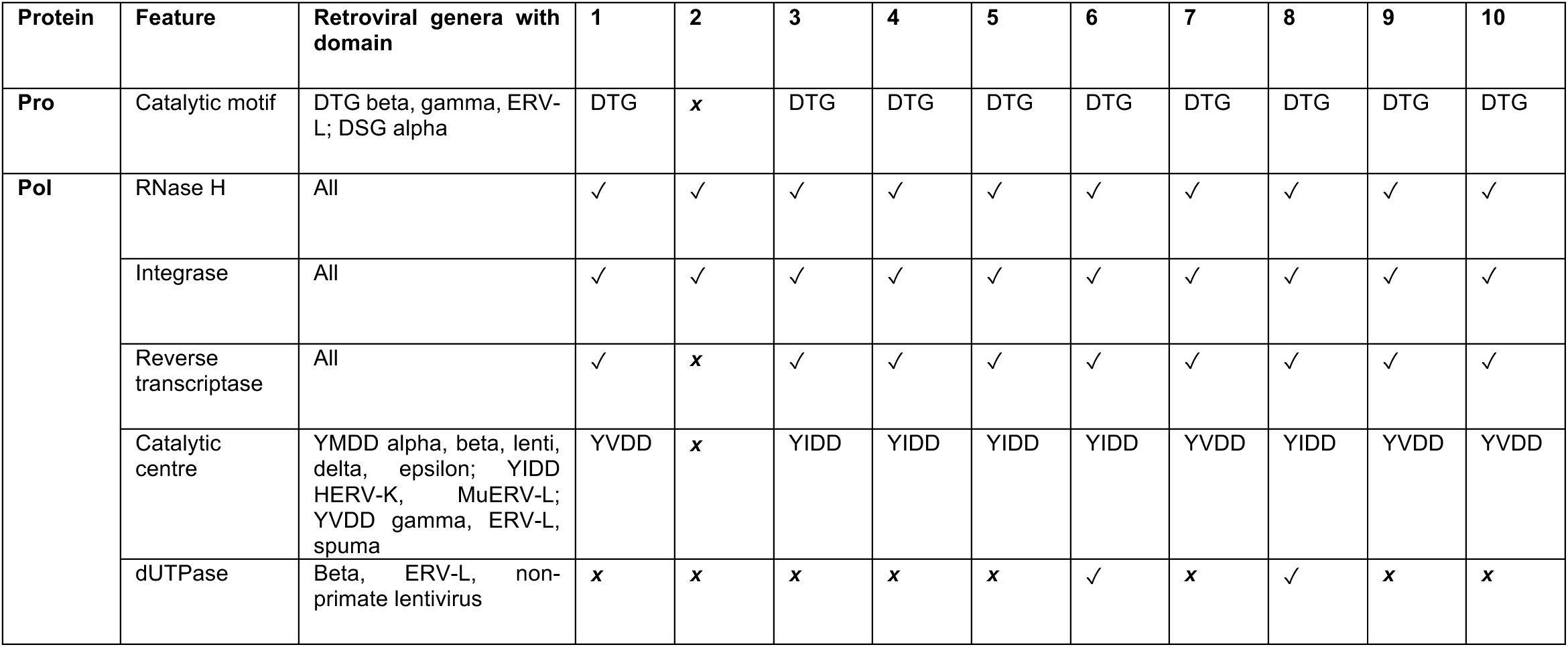
Conserved domains identified in Pol of the 10 AnERVs (labelled 1-10) identified in 11 species and their inclusion in different retroviral genera.

| Protein | Feature | Retroviral genera with domain | 1 | 2 | 3 | 4 | 5 | 6 | 7 | 8 | 9 | 10 |
| --- | --- | --- | --- | --- | --- | --- | --- | --- | --- | --- | --- | --- |
| **Pro** | Catalytic motif | DTG beta, gamma, ERV-L; DSG alpha | DTG | x | DTG | DTG | DTG | DTG | DTG | DTG | DTG | DTG |
| **Pol** | RNase H | All | ✓ | ✓ | ✓ | ✓ | ✓ | ✓ | ✓ | ✓ | ✓ | ✓ |
|  | Integrase | All | ✓ | ✓ | ✓ | ✓ | ✓ | ✓ | ✓ | ✓ | ✓ | ✓ |
|  | Reverse transcriptase | All | ✓ | x | ✓ | ✓ | ✓ | ✓ | ✓ | ✓ | ✓ | ✓ |
|  | Catalytic centre | YMDD alpha, beta, lenti, delta, epsilon; YIDD HERV-K, MuERV-L; YVDD gamma, ERV-L, spuma | YVDD | x | YIDD | YIDD | YIDD | YIDD | YVDD | YIDD | YVDD | YVDD |
|  | dUTPase | Beta, ERV-L, non-primate lentivirus | x | x | x | x | x | ✓ | x | ✓ | x | x |

#### 3.4.3. Env Conserved Domains

The Env protein can be split into the two subunits: surface (SU) and transmembrane (TM). Through identification of the conserved domains in Env, the structure (covalently associated found in alpharetrovirus, type-D betaretrovirus, gammaretrovirus, and deltaretrovirus genera, or non-covalently associated found in betaretrovirus and lentivirus) can be used to determine the classification the novel AnERVs. The SU domain of the covalently associated Env structures contain an isomerase domain, which in type-D betaretrovirus, deltaretrovirus and gammaretrovirus is represented by CXXC but is not present in alpharetrovirus (Henzy & Johnson, 2013; Hogan & Johnson, 2023; Pinter et al., 1997). This motif was identified in all AnERVs except AnERV2 (**Table 4**) which had a distinct lack of conserved domains in Gag, Pol and Pro. The isomerase domain was identical in AnERV1, AnERV2, AnERV5, AnERV7, AnERV8 and AnERV9 (CWVC). In AnERV9 this domain was identified as CWIC. The isomerase domain was not present in AnERV3. The furin cleavage site where the SU and TM are cleaved is present in all groups (Hosaka et al., 1991). In AnERV1, AnERV2 and AnERV5 there are 2-3 more amino acids interrupting the sequence; in AnERV2 these insertions are present across >50% of the alignment (AnERV1 and AnERV5 currently have too few full length loci to determine the prevalence of these insertions). It is evident that all insertions have covalently associated Env structure by the presence of characteristic heptad repeats which flank an immunosuppressive domain. The fusion peptide (FP) can be distinctive for the genera of Env, where alpharetrovirus FP is present approximately nine amino acids from the furin cleavage site (as opposed to immediately after the cleavage site) and is flanked by cysteines; this structure was identified in AnERV3 and AnERV6. Further support for the classification of env in AnERV3 is provided the lack of isomerase domain in the SU of Env (Henzy & Coffin, 2013b; Henzy & Johnson, 2013).

**Table 4:** Conserved domains identified in Env surface (SU) and transmembrane (TM) subunit of the 10 AnERVs (labelled 1-10) identified in 11 species and their inclusion in different retroviral genera.

| Env region | Feature | Retroviral genera with domain | 1 | 2 | 3 | 4 | 5 | 6 | 7 | 8 | 9 | 10 |
| --- | --- | --- | --- | --- | --- | --- | --- | --- | --- | --- | --- | --- |
| **SU subunit** | Isomerase domain | All but alpha | CWVC | CWVC | x | CWIC | CWVC | CFNC | CWVC | CWVC | CWVC | CWIC |
|  | Furin cleavage site | All | RIRNYR | RVRNYR | RDRR | RNKR | RKDTV<br>R | KFRR | RTRR | RTRR | RFKR | KQKR |
| **TM subunit** | Fusion peptide | Type-D beta, delta, gamma; flanked by cystines (C) in alpha | ✓ | ✓ | ✓<br>C flanked | ✓ | ✓ | ✓<br>C flanked | ✓ | ✓ | ✓ | ✓ |
|  | Glycosylation motif | Alpha, type-D beta, delta, gamma | ✓ | ✓ | ✓ | ✓ | ✓ | ✓ | ✓ | ✓ | ✓ | ✓ |
|  | Heptad repeats | Alpha, type-D beta, delta, gamma | ✓ | ✓ | ✓ | ✓ | ✓ | ✓ | ✓ | ✓ | ✓ | ✓ |
|  | Immuno-suppressive domain | Alpha, type-D beta, delta, gamma | ✓ | ✓ | ✓ | ✓ | ✓ | ✓ | ✓ | ✓ | ✓ | ✓ |

### 3.5. Phylogenetic Analysis

To further classify the novel AnERVs, phylogenetic analysis was undertaken on each of the three translated genes (Gag, the RT of Pol and the TM of Env) against known retroviruses (**Table S2**).

#### 3.5.1. Gag

The Gag protein of the AnERVs had poor identity with known retroviruses in alignment, and so the phylogenetic analysis was undertaken for each genus of retroviruses separately. All ten AnERVs appeared to branch separately from alpha-, delta- and gamma- retrovirus (**Figure S2**). AnERV6 and 8 appear to branch separately from the remaining groups, showing clustering with betaretrovirus and ERV-L sequences, both with strong posterior values (**Figure 4A, 4B**). AnERV1 and 7 cluster strongly with two epsilon-ERVs identified in *Bufo gargarizans* (Y. Chen et al., 2022), and AnERV2 clusters with fish epsilon- ERVs but has a weak posterior value on basal branches (**Fgure 4C**).

**Figure 4:**
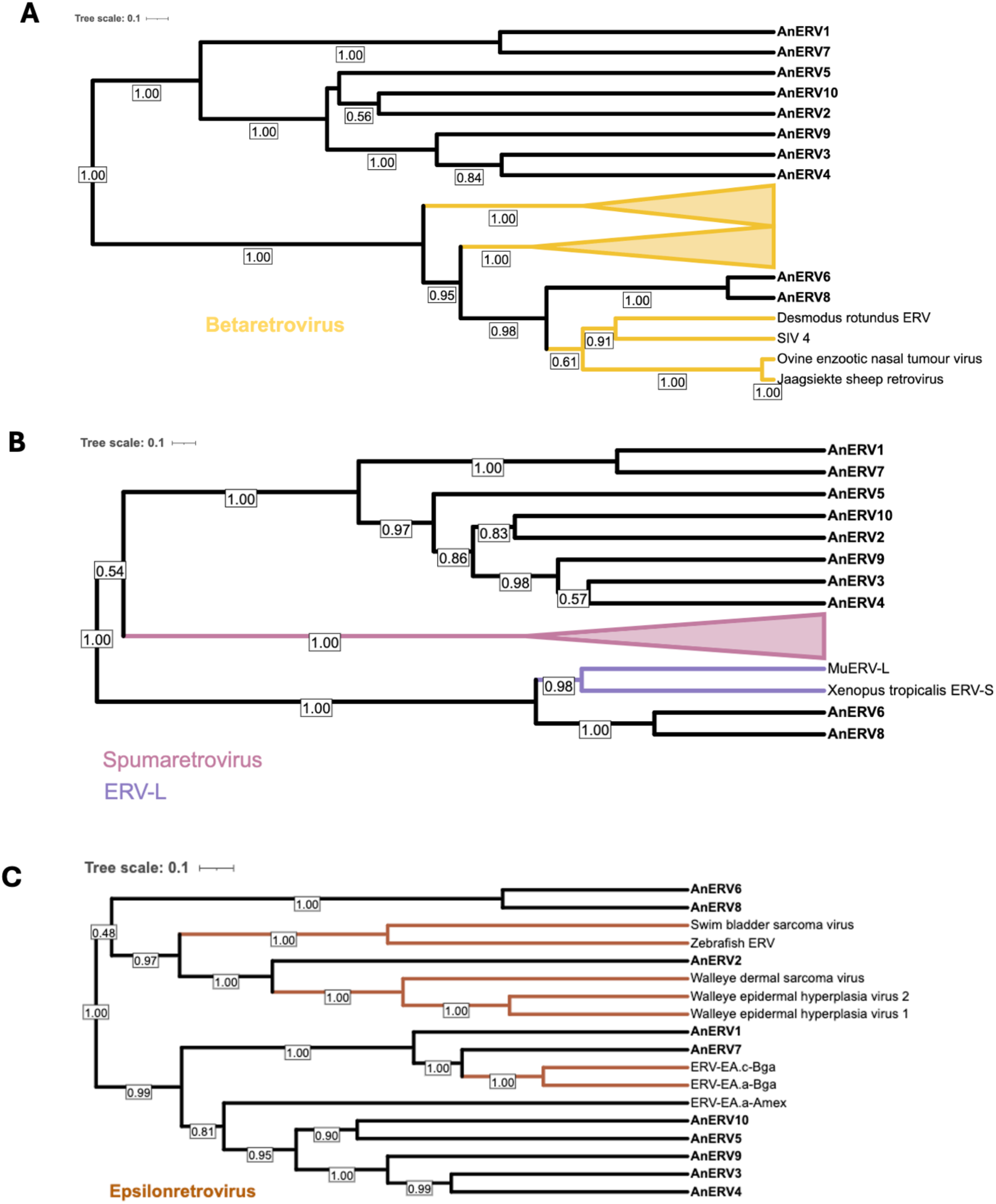
Phylogenetic relationship of Gag protein from the 10 AnERVs identified in 11 species with **A** betaretrovirus; **B** spumaretroviruses and ERV-L and **C** epsilonretrovirus. AnERV6 and 8 appear to cluster with ERV-L and AnERV2 with epsilonretrovirus. The trees were inferred using Bayesian phylogeny with BEAST and visualised with the interactive tree of life webpage. Posterior values are displayed on nodes, only those >0.50 are shown.

#### 3.5.2. Pol

Phylogenetic analysis of the RT of the Pol protein showed all AnERVs clustering with epsilonretroviruses (**Figure 5**), with the exception of AnERV6 and AnERV8 which cluster with ERV-L. AnERV2 was not included in Pol RT phylogenetic analysis as it lacked a clear RT domain.

**Figure 5:**
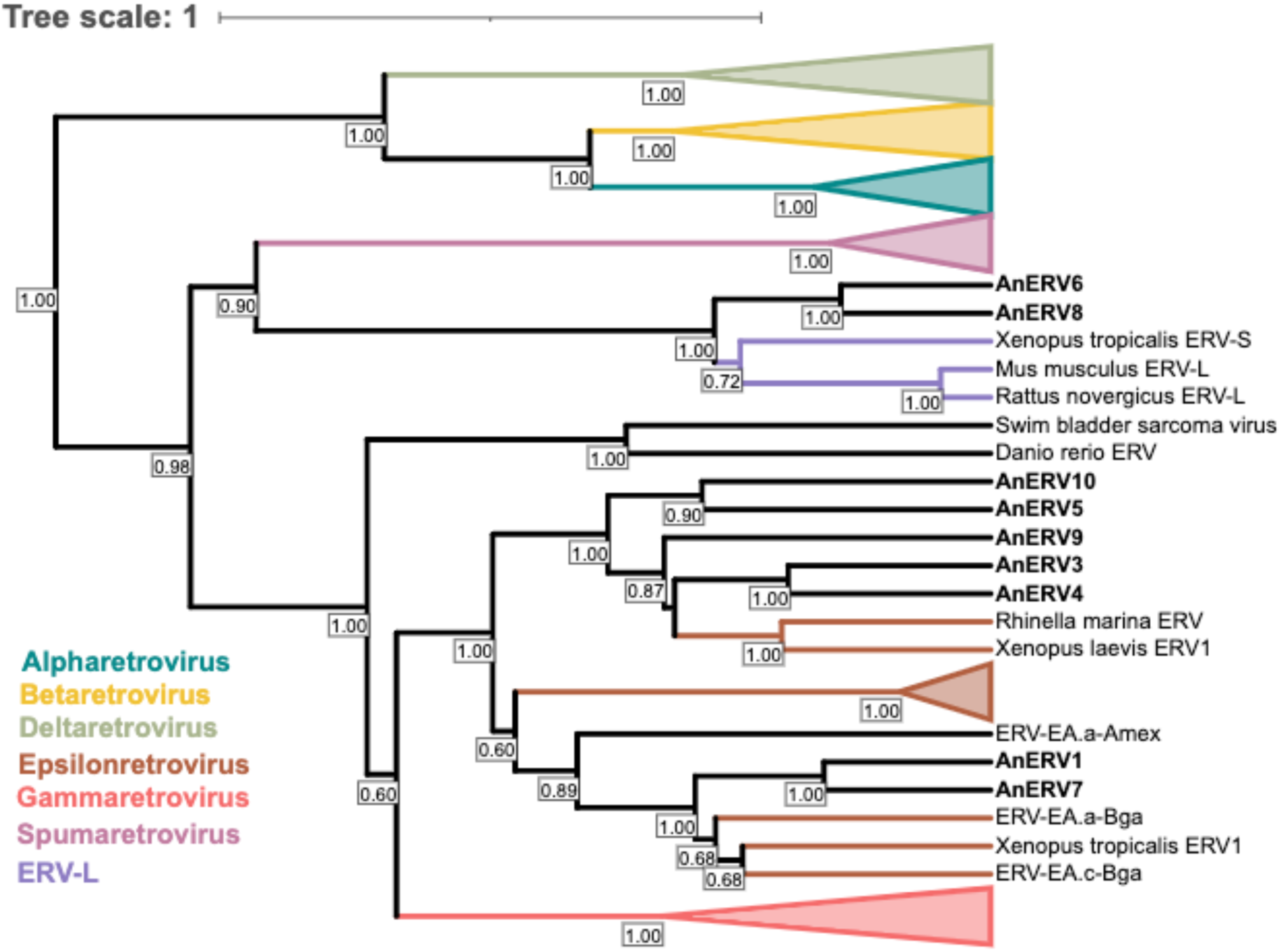
Phylogenetic relationship of Pol protein reverse transcriptase from seven AnERVs identified in 11 species in this study. AnERV2 was not included as reverse transcriptase was not present. AnERV6 and AnERV8 appear most closely related to ERV-L and spumaretrovirus; the remaining groups appear to cluster with epsilonretrovirus. The trees were inferred using Bayesian phylogeny with BEAST and visualised with the interactive tree of life webpage. Posterior values are displayed on nodes, only those >0.50 are shown.

#### 3.5.3. Env

For the phylogenetic analysis of the Env TM region epsilonretrovirus sequences were excluded due to their complex Env structure resulting in a poor sequence alignment. All AnERVs excluding AnERV3 and AnERV6 cluster with gammaretrovirus with high posterior values (**Figure 6**). AnERV3 and AnERV6 cluster with alpharetrovirus Env with high posterior values, which again is supported by the presence of a cysteine flanked fusion peptide domain, typically found in alpharetrovirus.

**Figure 6:**
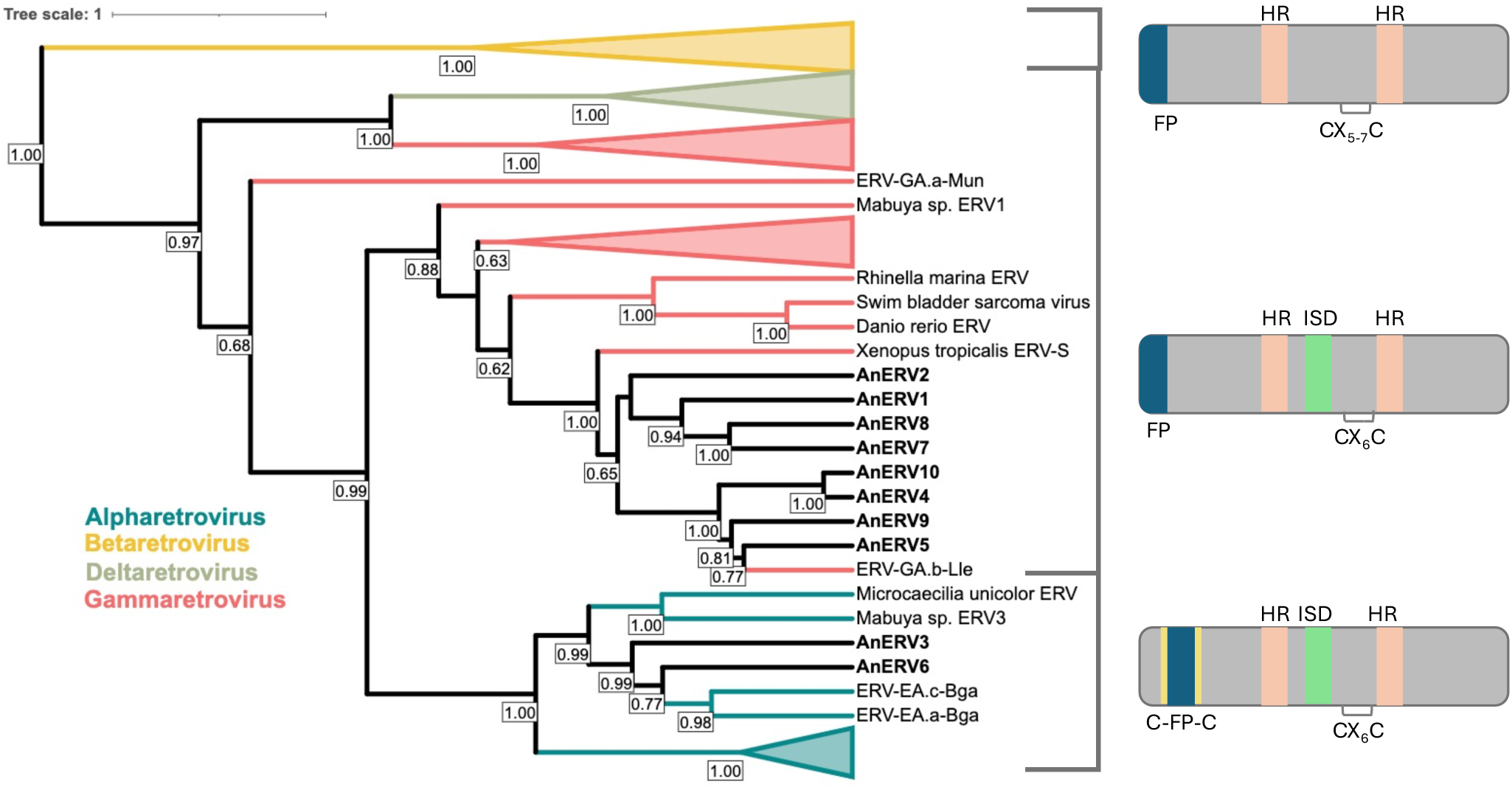
Phylogenetic relationship of Env protein transmembrane (TM) subunit of 10 AnERVs identified in 11 species with representative retroviruses. AnERV3 and AnERV6 appear most closely related to alpharetrovirus; the remaining groups appear to cluster with gammaretrovirus. Epsilonretrovirus were not included in the phylogeny due to the difference in Env TM structure resulting in a poor sequence alignment. The trees were inferred using Bayesian phylogeny with BEAST and visualised with the interactive tree of life webpage. On the right-hand side are the structures of the Env TM subunit: betaretrovirus has non-covalently associated Env; gammaretrovirus has covalently associated Env; alpharetrovirus has covalently associated Env with cysteine-flanked fusion peptide (C-FP-C). FP = fusion peptide; HR = heptad repeats; ISD = immunosuppressive domain. Posterior values are displayed on nodes, only those >0.50 are shown.

#### 3.5.4. Phylogenetic Analysis Summary

The phylogenetic analyses on each gene highlighted the recombinant nature of the novel AnERVs we have identified. A summary of the phylogenetic analysis is provided in **Table 5**.

**Table 5:** Summary of the phylogenetic analysis of each protein/protein region of the 10 AnERVs.

| Group | Species | gag | pol RT | env TM |
| --- | --- | --- | --- | --- |
| AnERV1 | *Spea multiplicata* | epsilon | epsilon | gamma |
| AnERV2 | *Spea multiplicata* | epsilon | not present | gamma |
|  | *Spea bombifrons* |  |  |  |
|  | *Spea hammondi* |  |  |  |
| AnERV3 | *Pelobates cultripes* | undetermined | epsilon | alpha |
| AnERV4 | *Bufo bufo* | undetermined | epsilon | gamma |
|  | *Bufo gargarizans* |  |  |  |
| AnERV5 | *Lithobates sylvaticus* | undetermined | epsilon | gamma |
|  | *Glandirana rugosa* |  |  |  |
| AnERV6 | *Gastrophryne carolinensis* | ERV-L | ERV-L | alpha |
| AnERV7 | *Bombina bombina* | epsilon | epsilon | gamma |
| AnERV8 | *Bombina bombina* | ERV-L | ERV-L | gamma |
| AnERV9 | *Gastrophryne carolinensis* | undetermined | epsilon | gamma |
| AnERV10 | *Xenopus tropicalis* | undetermined | epsilon | gamma |

## 4. Discussion

In this study we screened Anuran genomes to identify and classify alpha- ERVs. This process has resulted in the identification of ten novel ERV families in 11 Anuran species (five frogs and six toads) termed AnERV1-10, through pinpointing the *env* gene and walking outward to classify the flanking LTRs, identifying solo LTRs and full-length ERV elements encoding the remaining internal genes. Screening with this method did identify ERVs beyond the alpharetrovirus genera. Identifying these sequences was possible using the structure of the Env transmembrane (TM) subunit, which is also present in gammaretrovirus, deltaretrovirus and type-D betaretroviruses. These retroviral genera have an Env TM subunit which is categorised based on its ability to covalently associate to the surface (SU) Env subunit. The Env structure of alpharetrovirus is distinctive in the location and composition of the fusion peptide domain (Bénit et al., 2001; Henzy & Coffin, 2013b; Henzy & Johnson, 2013). The majority of previous studies of anuran sequences have identified single unique ERVs (Aiewsakun & Katzourakis, 2017; Y. Chen et al., 2021, 2022; Herniou et al., 1998; Kambol et al., 2003; Martin et al., 1999; Russo et al., 2018; Sinzelle et al., 2011; Tristem et al., 1996; Yedavalli et al., 2021), with one study describing several: four gamma- and 29 epsilon- ERVs from nine species (Y. Chen et al., 2022).

Based on Pol RT classification, seven of the ten novel ERVs described here (AnERV1, AnERV3, AnERV4, AnERV5, AnERV7, AnERV9 and AnERV10) classify as epsilonretrovirus, although AnERV1 and AnERV7 appear in a distinct branch to AnERV3, AnERV4, AnERV5 and AnERV10. Through analysis of the Pol protein, these AnERVs are phylogenetically most closely related to Anuran ERVs previously described, and distinct from those identified in fish genomes (Kambol et al., 2003; Russo et al., 2018; Sinzelle et al., 2011). However, the phylogenetic relationship of the Env TM subunit of the identified elements, as well as assessment of their conserved domains, showed that all elements have a covalently associated Env which is not typical of epsilonretrovirus (Henzy & Coffin, 2013b; Hogan & Johnson, 2023; Paul et al., 2006). The Env of AnERV1, AnERV2, AnERV4, AnERV5, AnERV7, AnERV8, AnERV9 and AnERV10 cluster with gammaretroviruses, most closely related to two identified previously in Anura, of which XtERV-S also exhibits variable classification depending on whether the Pol or Env proteins are used (Chen et al., 2022; Yedavalli et al., 2021).

Incongruences in conserved domain and phylogenetic analysis of Pol and Env in AnERV3 and AnERV6 suggest recombination has played a role in their evolutionary history. This proposal is supported by evidence of recombination amongst epsilonretrovirus related ERVs reported in amphibian and fish species (Paul et al., 2006; Yedavalli et al., 2021; Chen et al., 2022), including those which have covalently associated Env structures. AnERV3 clusters with epsilonretrovirus based on the Pol protein and has a similar structure to that of an ERV identified in *Bufo gargarizans* from a large screening study of amphibian and fish species, which classified as an epsilonretrovirus by Pol, but the Env was identified as an alpharetrovirus Env structure (Chen et al., 2022). However, AnERV3 does not cluster with these sequences based on the Pol protein (ERV-EA.a-Bga and ERV-EA.c-Bag in **Figure 5**). In amphibians, alpharetrovirus Env structure has also been described in ERVs in the sister order to Anura, in two caecilian species, whereby the insertions were closely related to betaretrovirus based on the Pol (Chen et al., 2021). Other than avian species, ERVs with this Env structure have only been previously described in a series of syncytin-like genes (those which have integral roles in placental formation co-opted from ERVs) in a Mabuya lizard species (Cornelis et al., 2017; Dupressoir et al., 2005; Mi et al., 2000). Here we identify alpharetrovirus Env structure in two AnERV families (AnERV3 and AnERV6) from two different Anuran species, *Pelobates cultripes* and *Gastrophyrne carolinensis*. However, these two families did not share similarity in the Pol phylogeny or conserved domains; AnERV3 clustered with AnERV1, AnERV4, AnERV5, AnERV7, AnERV9 and AnERV10 with epsilonretrovirus, whereas AnERV6 clustered with AnERV8 alongside ERV-L.

The classification of AnERV6, with the Pol most closely related to an ancient group of retroviruses identified in mammals called ERV-L, was similarly observed in AnERV8. These two AnERV families contain a dUTPase motif at the 3’-terminus of Pol, and although dUTPase can be present in multiple retroviral genera, this location is most alike to that of ERV-L. While ERV-L are termed retroviruses, in mammals an *env* gene has not to date been found associated with them (Barabás et al., 2003; Bénit et al., 1997, 1999; Hizi et al., 1989). However, an ERV-L with a covalently associated Env structure has been previously identified in the Anuran species *Xenopus tropicalis* (XtERV-S), and here AnERV8 shows a similar classification and a close phylogenetic relationship: Pol classifying as ERV-L and Env with a covalently associated structure (**Figure 3, 5, 6**; Yedavalli et al., 2021). The group of ERV-L have predominantly been described in mammals and are phylogenetically most closely related to Spumaretroviridae, which are one of the most basal groups of Retroviridae (Bénit et al., 1997, 1999; Wang & Han, 2021). The ERV-L-like AnERVs identified here, AnERV6 and AnERV8, appear to have independently acquired *env* from an alpharetrovirus and gammaretrovirus respectively; this acquisition could facilitate access to a new host species (XtERV appears to have similarly acquired a gamma *env*).

It is evident that recombination is present amongst Anuran ERVs and resulting incongruences between Pol and Env highlight the importance of investigating more than the *pol* gene alone. The recombination identified here, particularly involving the *env* gene, is not an uncommon observation in ERVs (Chabaukswar et al., 2023). A well characterised example of this is the baboon endogenous retrovirus (BaEV) in the emergence of domestic cat ERV RD-114, which has *gag-pol* of domestic cat ERV-DC and the *env* of BaEV (Anai et al., 2012; Kuyl et al., 1999). This can occur through multiple means, such as interactions between exogenous retroviruses with ERVs in the host genome, leading to the emergence of recombinant variants, as seen within murine and feline leukaemia virus (Chiu & Vandewoude, 2021; Evans et al., 2009; Levy, 2008). The exchange of *env* can also occur between ERV classes, potentially enabling access to new host species; cross-species transmission has been hypothesised for ERVs found in pythons, whereby a betaretrovirus has gained a gammaretrovirus *env,* which supposedly allowed a retroviral genus which is typically restricted to birds and mammals (betaretrovirus) into these reptiles (Henzy & Johnson, 2013; Huder et al., 2002).

The discovery of 10 novel AnERV families (with multiple loci) in 11 Anuran species adds to the complex evolutionary history of retroviruses; by shedding light upon the recombination events that occur over time we can further understand ERV evolution within vertebrate genomes, and the presence and role of ERVs in vertebrate evolution.

## Notes

### Competing Interest Statement

The authors have declared no competing interest.

## References

Aiewsakun, P., & Katzourakis, A. (2017). Marine origin of retroviruses in the early Palaeozoic Era. Nature Communications 2017 8:1, 8(1), 1–12. 10.1038/ncomms13954

Altschul, S. F., Gish, W., Miller, W., Myers, E. W., & Lipman, D. J. (1990). Basic local alignment search tool. Journal of Molecular Biology, 215(3), 403–410. 10.1016/S0022-2836(05)80360-2

Anai, Y., Ochi, H., Watanabe, S., Nakagawa, S., Kawamura, M., Gojobori, T., & Nishigaki, K. (2012). Infectious Endogenous Retroviruses in Cats and Emergence of Recombinant Viruses. Journal of Virology, 86(16), 8634–8644. 10.1128/JVI.00280-12/SUPPL_FILE/ZJV999096331SO1.PDF

Barabás, O., Rumlová, M., Erdei, A., Pongrácz, V., Pichová, I., & Vértessy, B. G. (2003). dUTPase and Nucleocapsid Polypeptides of the Mason-Pfizer Monkey Virus Form a Fusion Protein in the Virion with Homotrimeric Organization and Low Catalytic Efficiency. Journal of Biological Chemistry, 278(40), 38803–38812. 10.1074/JBC.M306967200

Bénit, L., De Parseval, N., Casella, J.-F. O., Callebaut, I., Agnè, A., Cordonnier, A., And, †, & Heidmann, T. (1997). Cloning of a new murine endogenous retrovirus, MuERV-L, with strong similarity to the human HERV-L element and with a gag coding sequence closely related to the Fv1 restriction gene. Journal of Virology, 71(7), 5652–5657. 10.1128/JVI.71.7.5652-5657.1997

Bénit, L., Dessen, P., & Heidmann, T. (2001). Identification, Phylogeny, and Evolution of Retroviral Elements Based on Their Envelope Genes. Journal of Virology, 75(23), 11709–11719. 10.1128/JVI.75.23.11709-11719.2001/ASSET/E5A9C2C9-E7BE-4C4E-A1B4-EACDDFFCC87F/ASSETS/GRAPHIC/JV2311162005.JPEG

Bénit, L., Lallemand, J.-B., Casella, J.-F., Philippe, H., & Heidmann, T. (1999). ERV-L Elements: a Family of Endogenous Retrovirus-Like Elements Active throughout the Evolution of Mammals. Journal of Virology, 73(4), 3301–3308. 10.1128/JVI.73.4.3301-3308.1999/ASSET/E697912B-5389-4A9A-B1E5-1A68342AE882/ASSETS/GRAPHIC/JV0491617005.JPEG

Bergman, A. C., Björnberg, O., Nord, J., Nyman, P. O., & Rosengren, A. M. (1994). The Protein p30, Encoded at the gag-pro Junction of Mouse Mammary Tumor Virus, is a dUTPase Fused with a Nucleocapsid Protein. Virology, 204(1), 420–421. 10.1006/VIRO.1994.1547

Boeke, J., & Stoye, J. (1997). Retrotransposons, Endogenous Retroviruses, and the Evolution of Retroelements. Retroviruses. https://www.ncbi.nlm.nih.gov/books/NBK19468/

Bolisetty, M., Blomberg, J., Benachenhou, F., Sperber, G., & Beemon, K. (2012). Unexpected diversity and expression of avian endogenous retroviruses. MBio, 3(5). 10.1128/mBio.00344-12

Chabukswar, S., Grandi, N., Lin, L.T. and Tramontano, E., 2023. Envelope Recombination: A Major Driver in Shaping Retroviral Diversification and Evolution within the Host Genome. Viruses, 15(9), p.1856.

Chameettachal, A., Mustafa, F., & Rizvi, T. A. (2023). Understanding Retroviral Life Cycle and its Genomic RNA Packaging. Journal of Molecular Biology, 435(3), 167924. 10.1016/J.JMB.2022.167924

Chen, M., Guo, X., & Zhang, L. (2021). Unexpected Discovery and Expression of Amphibian Class II Endogenous Retroviruses. Journal of Virology, 95(3). 10.1128/JVI.01806-20/SUPPL_FILE/JVI.01806-20-S0001.PDF

Chen, Y., Wang, X., Liao, M.-E., Song, Y., Zhang, Y.-Y., & Cui, J. (2022). Evolution and Genetic Diversity of the Retroviral Envelope in Anamniotes. Journal of Virology, 96(8). 10.1128/JVI.02072-21

Chen, Y., Zhang, Y.-Y., Wei, X., & Cui, J. (2021). Multiple Infiltration and Cross-Species Transmission of Foamy Viruses across the Paleozoic to the Cenozoic Era. Journal of Virology, 95(14). 10.1128/JVI.00484-21/SUPPL_FILE/JVI.00484-21-S0001.PDF

Chiu, E. S., & Vandewoude, S. (2021). Endogenous Retroviruses Drive Resistance and Promotion of Exogenous Retroviral Homologs. Annual Review of Animal Biosciences, 9(Volume 9, 2021), 225–248. 10.1146/ANNUREV-ANIMAL-050620-101416/CITE/REFWORKS

Chu, L., Su, F., Han, G. Z., & Wang, J. (2023). Jawless vertebrates do not escape retrovirus infection. Virology, 583, 52–55. 10.1016/j.virol.2023.04.010

Coffin, J. M. (1992). Structure and Classification of Retroviruses. In The Retroviridae. 10.1007/978-1-4615-3372-6_2

Coffin, J. M., Hughes, S. H., & Varmus, Harold. (1997). Retroviruses. Cold Spring Harbor Laboratory Press.

Contreras-Galindo, R., Dube, D., Fujinaga, K., Kaplan, M. H., & Markovitz, D. M. (2017). Susceptibility of Human Endogenous Retrovirus Type K to Reverse Transcriptase Inhibitors. Journal of Virology, 91(23). 10.1128/JVI.01309-17/SUPPL_FILE/ZJV999183119S1.PDF

Cornelis, G., Funk, M., Vernochet, C., Leal, F., Tarazona, O. A., Meurice, G., Heidmann, O., Dupressoir, A., Miralles, A., Ramirez-Pinilla, M. P., Heidmann, T., & Roberts, R. M. (2017). An endogenous retroviral envelope syncytin and its cognate receptor identified in the viviparous placental Mabuya lizard. Proceedings of the National Academy of Sciences of the United States of America, 114(51), E10991–E11000. 10.1073/PNAS.1714590114/SUPPL_FILE/PNAS.1714590114.SD04.XLSX

Demirov, D. G., & Freed, E. O. (2004). Retrovirus budding. Virus Research, 106(2), 87–102. 10.1016/J.VIRUSRES.2004.08.007

Donahue, P. R., Hoover, E. A., Beltz, G. A., Riedel,’, N., Hirsch,’, V. M., Overbaugh, J., & Mullins’, J. I. (1988). Strong sequence conservation among horizontally transmissible, minimally pathogenic feline leukemia viruses. Journal of Virology, 62(3), 722–731. 10.1128/JVI.62.3.722-731.1988

Dupressoir, A., Lavialle, C., & Heidmann, T. (2012). From ancestral infectious retroviruses to bona fide cellular genes: Role of the captured syncytins in placentation. Placenta, 33(9), 663–671. 10.1016/J.PLACENTA.2012.05.005

Dupressoir, A., Marceau, G., Vernochet, C., Bénit, L., Kanellopoulos, C., Sapin, V., & Heidmann, T. (2005). Syncytin-A and syncytin-B, two fusogenic placenta-specific murine envelope genes of retroviral origin conserved in Muridae. Proceedings of the National Academy of Sciences of the United States of America, 102(3), 725–730. 10.1073/PNAS.0406509102

Evans, L. H., Alamgir, A. S. M., Owens, N., Weber, N., Virtaneva, K., Barbian, K., Babar, A., Malik, F., & Rosenke, K. (2009). Mobilization of Endogenous Retroviruses in Mice after Infection with an Exogenous Retrovirus. Journal of Virology, 83(6), 2429–2435. 10.1128/JVI.01926-08/ASSET/C2321EFE-1B3E-44F5-B713-1392A8AFD705/ASSETS/GRAPHIC/ZJV0060916480005.JPEG

Gifford, R., Blomberg, J., Coffin, J. M., Fan, H., Heidmann, T., Mayer, J., Stoye, J., Tristem, M., & Johnson, W. E. (2018). Nomenclature for endogenous retrovirus (ERV) loci. Retrovirology, 15(1), 1–11. 10.1186/S12977-018-0442-1/FIGURES/4

Gifford, R., Kabat, P., Martin, J., Lynch, C., & Tristem, M. (2005). Evolution and distribution of class II-related endogenous retroviruses. Journal of Virology, 79(10), 6478–6486. 10.1128/JVI.79.10.6478-6486.2005

Gifford, R., & Tristem, M. (2003). The Evolution, Distribution and Diversity of Endogenous Retroviruses. Virus Genes 2003 26:3, 26(3), 291–315. 10.1023/A:1024455415443

Gottlinger, H. G., Sodroski, J. G., & Haseltine, W. A. (1989). Role of capsid precursor processing and myristoylation in morphogenesis and infectivity of human immunodeficiency virus type 1. Proceedings of the National Academy of Sciences of the United States of America, 86(15), 5781. 10.1073/PNAS.86.15.5781

Harrison, J. J. E. K., Tuske, S., Das, K., Ruiz, F. X., Bauman, J. D., Boyer, P. L., Destefano, J. J., Hughes, S. H., & Arnold, E. (2021). Crystal structure of a retroviral polyprotein: Prototype foamy virus protease-reverse transcriptase (pr-rt). Viruses, 13(8), 1495. 10.3390/V13081495/S1

Henzy, J. E., & Coffin, J. M. (2013a). Betaretroviral Envelope Subunits Are Noncovalently Associated and Restricted to the Mammalian Class. Journal of Virology, 87(4), 1937–1946. 10.1128/JVI.01442-12/ASSET/2251F669-94BF-4A5A-890C-F3E1BE77BC05/ASSETS/GRAPHIC/ZJV9990972370005.JPEG

Henzy, J. E., & Coffin, J. M. (2013b). Betaretroviral Envelope Subunits Are Noncovalently Associated and Restricted to the Mammalian Class. Journal of Virology, 87(4), 1937. 10.1128/JVI.01442-12

Henzy, J. E., Gifford, R. J., Johnson, W. E., & Coffin, J. M. (2014). A Novel Recombinant Retrovirus in the Genomes of Modern Birds Combines Features of Avian and Mammalian Retroviruses. Journal of Virology, 88(5). 10.1128/jvi.02863-13

Henzy, J. E., & Johnson, W. E. (2013). Pushing the endogenous envelope. Philosophical Transactions of the Royal Society B: Biological Sciences, 368(1626), 20120506. 10.1098/RSTB.2012.0506

Herniou, E., Martin, J., Miller, K., Cook, J., Wilkinson, M., & Tristem, M. (1998). Retroviral Diversity and Distribution in Vertebrates. Journal of Virology, 72(7), 5955–5966. 10.1128/JVI.72.7.5955-5966.1998/ASSET/14DC1DC1-736F-44E5-BBE1-2D86989F1F70/ASSETS/GRAPHIC/JV0781750004.JPEG

Hizi, A., Henderson, L. E., Copeland, T. D., Sowder, R. C., Hixson, C. V., & Oroszlan, S. (1987). Characterization of mouse mammary tumor virus gag-pro gene products and the ribosomal frameshift site by protein sequencing. Proceedings of the National Academy of Sciences, 84(20), 7041–7045. 10.1073/PNAS.84.20.7041

Hizi, A., Henderson, L. E., Copeland, T. D., Sowder, R. C., Krutzsch, H. C., & Oroszlan’, S. (1989). Analysis of gag proteins from mouse mammary tumor virus. Journal of Virology, 63(6), 2543–2549. 10.1128/JVI.63.6.2543-2549.1989

Hogan, V., & Johnson, W. E. (2023). Unique Structure and Distinctive Properties of the Ancient and Ubiquitous Gamma-Type Envelope Glycoprotein. Viruses 2023, Vol. 15, Page 274, 15(2), 274. 10.3390/V15020274

Hosaka, M., Nagahama, M., Kim, W. S., Watanabe, T., Hatsuzawa, K., Ikemizu, J., Murakami, K., & Nakayama, K. (1991). Arg-X-Lys/Arg-Arg motif as a signal for precursor cleavage catalyzed by furin within the constitutive secretory pathway. Journal of Biological Chemistry, 266(19), 12127–12130. 10.1016/S0021-9258(18)98867-8

Huder, J. B., Böni, J., Hatt, J.-M., Soldati, G., Lutz, H., & Schüpbach, J. (2002). Identification and Characterization of Two Closely Related Unclassifiable Endogenous Retroviruses in Pythons (Python molurus and Python curtus). Journal of Virology, 76(15), 7607–7615. 10.1128/jvi.76.15.7607-7615.2002

Hütter, S., Zurnic, I., & Lindemann, D. (2013). Foamy Virus Budding and Release. Viruses, 5(4), 1075. 10.3390/V5041075

Jern, P., Sperber, G. O., & Blomberg, J. (2005). Use of Endogenous Retroviral Sequences (ERVs) and structural markers for retroviral phylogenetic inference and taxonomy. Retrovirology, 2. 10.1186/1742-4690-2-50

Kambol, R., Kabat, P., & Tristem, M. (2003). Complete nucleotide sequence of an endogenous retrovirus from the amphibian, Xenopus laevis. Virology, 311(1), 1–6. 10.1016/S0042-6822(03)00263-0

Kamei, M., Webb, G. C., Young, I. G., & Campbell, H. D. (1998). SOLH, a Human Homologue of theDrosophila melanogaster small optic lobesGene Is a Member of the Calpain and Zinc-Finger Gene Families and Maps to Human Chromosome 16p13.3 nearCATM(Cataract with Microphthalmia). Genomics, 51(2), 197–206. 10.1006/GENO.1998.5395

Katzourakis, A., Aiewsakun, P., Jia, H., Wolfe, N. D., LeBreton, M., Yoder, A. D., & Switzer, W. M. (2014). Discovery of prosimian and afrotherian foamy viruses and potential cross species transmissions amidst stable and ancient mammalian co-evolution. Retrovirology, 11(1), 1–17. 10.1186/1742-4690-11-61/FIGURES/4

Katzourakis, A., & Gifford, R. J. (2010). Endogenous Viral Elements in Animal Genomes. PLOS Genetics, 6(11), e1001191. 10.1371/JOURNAL.PGEN.1001191

Katzourakis, A., Rambaut, A., & Pybus, O. G. (2005). The evolutionary dynamics of endogenous retroviruses. Trends in Microbiology, 13(10), 463–468. 10.1016/J.TIM.2005.08.004

Konvalinka, J., Kräusslich, H. G., & Müller, B. (2015). Retroviral proteases and their roles in virion maturation. Virology, 479–480, 403–417. 10.1016/J.VIROL.2015.03.021

Kumar, S., Stecher, G., Suleski, M., & Hedges, S. B. (2017). TimeTree: A Resource for Timelines, Timetrees, and Divergence Times. Molecular Biology and Evolution, 34(7), 1812–1819. 10.1093/molbev/msx116

Kuyl, A. C. van der, Dekker, J. T., & Goudsmit, J. (1999). Discovery of a New Endogenous Type C Retrovirus (FcEV) in Cats: Evidence for RD-114 Being an FcEVGag-Pol/Baboon Endogenous Virus BaEVEnv Recombinant. Journal of Virology, 73(10), 7994. 10.1128/JVI.73.10.7994-8002.1999

Lander, S., Linton, L. M., Birren, B., Nusbaum, C., Zody, M. C., Baldwin, J., Devon, K., Dewar, K., Doyle, M., FitzHugh, W., Funke, R., Gage, D., Harris, K., Heaford, A., Howland, J., Kann, L., Lehoczky, J., LeVine, R., McEwan, P., … Yeh, R.-F. (2001). Initial sequencing and analysis of the human genome International Human Genome Sequencing Consortium* The Sanger Centre: Beijing Genomics Institute/Human Genome Center. NATURE, 409. www.nature.com

Levy, L. S. (2008). Advances in Understanding Molecular Determinants in FeLV Pathology. Veterinary Immunology and Immunopathology, 123(1–2), 14. 10.1016/J.VETIMM.2008.01.008

Llorens, C., Muñoz-Pomer, A., Bernad, L., Botella, H., & Moya, A. (2009). Network dynamics of eukaryotic LTR retroelements beyond phylogenetic trees. Biology Direct, 4, 41. 10.1186/1745-6150-4-41

Martin, J., Herniou, E., Cook, J., O’Neill, R. W., & Tristem, M. (1999). Interclass Transmission and Phyletic Host Tracking in Murine Leukemia Virus-Related Retroviruses. Journal of Virology, 73(3), 2442–2449. 10.1128/JVI.73.3.2442-2449.1999/ASSET/E6A5BB6C-D3F1-4909-BC70-50E9974A6CA2/ASSETS/GRAPHIC/JV0390979005.JPEG

Mi, S., Xinhua, L., Xiang-ping, L., Veldman, GM., Finnerty, H., Racie, L., LaVallie, E., Tang, X. Y., Edouard, P., Howes, S., Keith, J. C., & McCoy, J. M. (2000). Syncytin is a captive retroviral envelope protein involved in human placental morphogenesis. Nature 2000 403:6771, 403(6771), 785–789. 10.1038/35001608

Nguyen, C. D., Mansfield, R. E., Leung, W., Vaz, P. M., Loughlin, F. E., Grant, R. P., & MacKay, J. P. (2011). Characterization of a Family of RanBP2-Type Zinc Fingers that Can Recognize Single-Stranded RNA. Journal of Molecular Biology, 407(2), 273–283. 10.1016/J.JMB.2010.12.041

Pastuzyn, E. D., Day, C. E., Kearns, R. B., Kyrke-Smith, M., Taibi, A. V., McCormick, J., Yoder, N., Belnap, D. M., Erlendsson, S., Morado, D. R., Briggs, J. A. G., Feschotte, C., & Shepherd, J. D. (2018). The Neuronal Gene Arc Encodes a Repurposed Retrotransposon Gag Protein that Mediates Intercellular RNA Transfer. Cell, 172(1–2), 275–288.e18. 10.1016/j.cell.2017.12.024

Paul, T. A., Quackenbush, S. L., Sutton, C., Casey, R. N., Bowser, P. R., & Casey, J. W. (2006). Identification and Characterization of an Exogenous Retrovirus from Atlantic Salmon Swim Bladder Sarcomas. Journal of Virology, 80(6), 2941–2948. 10.1128/JVI.80.6.2941-2948.2006/ASSET/76BB40EE-71DB-426F-A322-63CF80F69CD1/ASSETS/GRAPHIC/ZJV0060675360005.JPEG

Pinter, A., Kopelman, R., Li, Z., Kayman, S. C., & Sanders, D. A. (1997). Localization of the labile disulfide bond between SU and TM of the murine leukemia virus envelope protein complex to a highly conserved CWLC motif in SU that resembles the active-site sequence of thiol-disulfide exchange enzymes. Journal of Virology, 71(10), 8073. 10.1128/JVI.71.10.8073-8077.1997

Poch, O., Sauvaget, I., Delarue, M., & Tordo, N. (1989). Identification of four conserved motifs among the RNA-dependent polymerase encoding elements. The EMBO Journal, 8(12), 3867. 10.1002/J.1460-2075.1989.TB08565.X

Russo, A. G., Eden, J.-S., Enosi Tuipulotu, D., Shi, M., Selechnik, D., Shine, R., Rollins, L. A., Holmes, E. C., & White, P. A. (2018). Viral Discovery in the Invasive Australian Cane Toad (Rhinella marina) Using Metatranscriptomic and Genomic Approaches. Journal of Virology, 92(17), 768–786. 10.1128/JVI.00768-18/SUPPL_FILE/ZJV017183801SD4.XLSX

Sharma, P. L., Nurpeisov, V., & Schinazi, R. F. (2005). Retrovirus Reverse Transcriptases Containing a Modified YXDD Motif. http://dx.doi.org/10.1177/095632020501600303, 16(3), 169–182. 10.1177/095632020501600303

Sinzelle, L., Carradec, Q., Paillard, E., Bronchain, O. J., & Pollet, N. (2011). Characterization of a Xenopus tropicalis Endogenous Retrovirus with Developmental and Stress-Dependent Expression. Journal of Virology, 85(5), 2167–2179. 10.1128/JVI.01979-10/SUPPL_FILE/SUPPLLEGENDS1979.ZIP

Strack, B., Calistri, A., Craig, S., Popova, E., & Göttlinger, H. G. (2003). AIP1/ALIX is a binding partner for HIV-1 p6 and EIAV p9 functioning in virus budding. Cell, 114(6), 689–699. 10.1016/S0092-8674(03)00653-6

Tristem, M., Herniou, E., Summers, K., & Cook, J. (1996). Three retroviral sequences in amphibians are distinct from those in mammals and birds. Journal of Virology, 70(7), 4864–4870. 10.1128/JVI.70.7.4864-4870.1996

Von Schwedler, U. K., Stuchell, M., Müller, B., Ward, D. M., Chung, H. Y., Morita, E., Wang, H. E., Davis, T., He, G. P., Cimbora, D. M., Scott, A., Kräusslich, H. G., Kaplan, J., Morham, S. G., & Sundquist, W. I. (2003). The protein network of HIV budding. Cell, 114(6), 701–713. 10.1016/S0092-8674(03)00714-1

Wang, J., & Han, G. Z. (2021). A Sister Lineage of Sampled Retroviruses Corroborates the Complex Evolution of Retroviruses. Molecular Biology and Evolution, 38(3), 1031–1039. 10.1093/MOLBEV/MSAA272

Waterston, R. H., Lindblad-Toh, K., Birney, E., Rogers, J., Abril, J. F., Agarwal, P., Agarwala, R., Ainscough, R., Alexandersson, M., An, P., Antonarakis, S. E., Attwood, J., Baertsch, R., Bailey, J., Barlow, K., Beck, S., Berry, E., Birren, B., Bloom, T., … Lander, E. S. (2002). Initial sequencing and comparative analysis of the mouse genome. Nature 2003 420:6915, 420(6915), 520–562. 10.1038/nature01262

Wlodawer, A., & Gustchina, A. (2000). Structural and biochemical studies of retroviral proteases. Biochimica et Biophysica Acta (BBA) - Protein Structure and Molecular Enzymology, 1477(1–2), 16–34. 10.1016/S0167-4838(99)00267-8

Wu, J., Matunis, M. J., Kraemer, D., Blobel, G., & Coutavas, E. (1995). Nup358, a Cytoplasmically Exposed Nucleoporin with Peptide Repeats, Ran-GTP Binding Sites, Zinc Fingers, a Cyclophilin A Homologous Domain, and a Leucine-rich Region. Journal of Biological Chemistry, 270(23), 14209–14213. 10.1074/JBC.270.23.14209

Yedavalli, V. R. K., Patil, A., Parrish, J., & Kozak, C. A. (2021). A novel class III endogenous retrovirus with a class I envelope gene in African frogs with an intact genome and developmentally regulated transcripts in Xenopus tropicalis. Retrovirology, 18(1), 1–16. 10.1186/S12977-021-00564-2/FIGURES/7

Yokoyama, N., Hayashi, N., Seki, T., Panté, N., Ohba, T., Nishii, K., Kuma, K., Hayashida, T., Miyata, T., Aebi, U., Fukui, M., & Nishimoto, T. (1995). A giant nucleopore protein that binds Ran/TC4. Nature 1995 376:6536, 376(6536), 184–188. 10.1038/376184a0

Zheng, J., Wei, Y., & Han, G. Z. (2022). The diversity and evolution of retroviruses: Perspectives from viral “fossils.” Virologica Sinica, 37(1), 11–18. 10.1016/J.VIRS.2022.01.019

