## Supplementary_tables_and_figures for "Identification of Novel Endogenous Retroviruses in Anura"

### Supplementary Material

### Figure S1

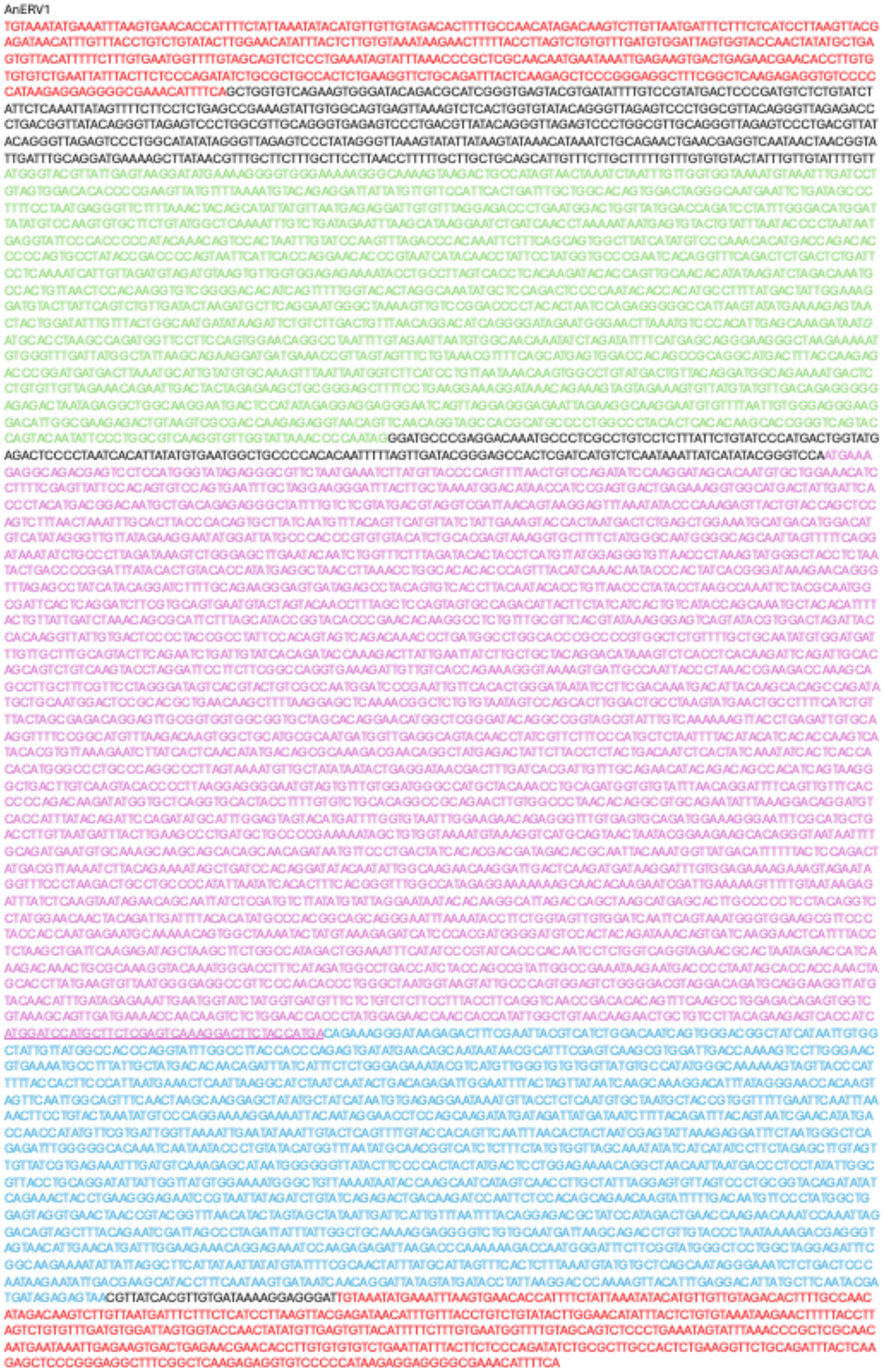

AnERV2

TGTGGAAGTTGAAGAATTTTGCTGATTCTGGATGTCAGCACAATGATCTTTACCTCATTGCTTATTCTGCTGATAGCAGATAAGATAACACTACAATCACCTGTCTACCTCA
TTTCTTAGTTTACTTATGTCTTGTTTGTGTCTCGGAAGAATGTGTACATATGGTATGTTTTCGTATGTTTTCTTTGCTCTAAGTGCCTTGTTGCAAAAGAATCAGAGTCCGCCA
CGACAAAGGACAAGTTGTCCCTTTCTTGACCTTCTGTAAGTAAAGATTGCACATGTCTGGGGGGGGGGTTATTTTTCTTCACCTCTTTAAACCGTGTGTGTCAGCAAATC
AAAGTAAATATACCAACAGATGGCTAGACAATATGCTAATAACTGTTTGCGCTTCTTTGCCTATATATTGTTAGAAAATTAAGGAGGGGGAGAGAAGTCTGCGAACTGGA
CTCTCCCCTGTGCGCACTTGAATAAAGAAGCTCTTGCTTACTCAACTCATTGAACTGGTGTCGTCAGATTTCTTGCCGACATTCACACTTGGGGGCTCGTCCGGGATC
CCAACAGTCTCGGAAGCGGTAACCGAAGGAGCGGACTAAACCTTGATCCACCTAATCCTTGGCCATCAAAAAGAGGTAAGGCACCTTTTGAAATGCCTATAGTTTAA
TTATTGGGTGGATGAGGGTAAGGTCGTCCTCGGTATTCCATCCGGGTGACGGGATCCAGAGTTGAGTAAGTAAGACCTATAAGATCTTGAATTGTTTAATTGTAACTATTG
TTTACTACATGGTCAGGGACCATTGTAAAGTACCTAGTTGTGAGAGACTAGTGTAACTATTGTTTACTACATGGTCAGGGACCATTGTAAAGTACCTAGTTGTGAGAGACT
AGTGTAAYTATTGTTTACTACATGGTCAGGGACCATTGTAAAGTACCTAGTTGTGAGAGACTAGTGTAAYTATTGTTTACTACATGGTCAGGGACCATTGTAAAGTACCTAGT
TGTGAGAGACTAGTGTAGCTATTTGAGTACATAATCACGGGCTATTGTAAAGTACCTAGTGGTGAGAACTAGTGTAAATATTTTGAAGTACATAGTCACGGATTATTGTAGC
GGAGGAGATGGGGAGTTTGTTGTCAAAAAATGTGCCTAAGGGGACGCCCTTAGAATATATGATAAGGAAACATGGAAAGGAGGCAGGAAAATATATGTCTCATTGGAA
GCRAATATCTGAAGAGGATAATGAAAGCCGTTATGTTTGGCCAGCTGGCGGTAGTTTGAATGTAGAAATACTAGATAATTTGAAAGAATGTTTGAATATGTGTGATGTGAAA
AGGAATGAGAGAATGATTTTCTTGAAATGGTATCAAGAAGGGTGGGACAACGTTAAAATACAGAAACAGCAGAAGTTACTAGACCCCACCTGTTCTGCCAATTCCCAA
TATCCTTGCCGTCCGGCAGGTAACTTTAAGAATCAAAAGGGTGCTCAAACACCTCAACCTGTTGAGACTTCTAAGAAAGCACAGGAGGTGTCTGTAACCCCAGGAC
CACCTAAGGTCCATAATTCGCTATATCCATCCTTAGATCCATCTGATCTATGGACTCAAGAATTATTTGACCCACCCCCATATAATGGACCTCAACCAGCAGACCCTAAT
ATGCCTGGTTCTAATCAACACCCTGCAGGTCCAGAAGCAGAGTTGCCACTTTCTTTGGAGCTTGTCCCATTTGTACCACCAGCTCCTATTTCACCCCTAGACCAAGAA
AGGCAACCTGATCCTCCTAAAATATCAAGGTCCAATATTCAGCCCACAGGACCATCATGTAGAACTAGATCCAGAAAGCCCCATTTAGATCCTCATTCAGATTCTGATG
CTGAGGAAAAAACATTTGTGCATGCTATGCCCATGAGAGAGACTGGGGGGGCTAGAGATGAGAGAGGTGTACCTCTATCTGTCCATCATCCATGGACACCAGGGGA
ATTATTGCTGTTGAAAGATCGATTCCCCAGACCTGAAATTGTTGGTCAGTTTTGTAAAGAATTACGTAGTGTGTCTATTCCTTATGATCCAGTGAAAAAGGATTTTATGCAAT
TGTTTGATTTGGTTATGACTGACGCAGAGAGAAAACAGTTAATTGAAGCTTATAACAGAATTCCTGATGTAACACCTTGAGCAAATGAACTGTTTACTAAGAGATTGAGTAC
TGAGAGACAGCAGACAATAGATTGTATAATACAAGCTTGTGGGACTGCATGGCTACAGCGACCTGACTGGGGAAAGATTAACTCTACAATGCAAGGAAAGAATGAATC
AGTAAGTGAATTCGCTGACAGAGTAGCGAAAGTTGTAATAGGACACAGTGGTGTGACAGATCCAGAGGTGAATGCAAGACAGTTGATTGTATCTTGTTTTGTAAAAGGG
TTGTTACCAGACTTCCATAAGGGTGTCACTAATAATGTTTTAGGATGGGAAGAGAAGCCGCTTTCTGAAATGATAGCAGCTGCTAAACATTTTGAAAAACAAAAGGAGTG
AATAAGGAACGTCAAGAGAAAGTTAGAGAGAAAGAACAGAGAATGTTGATGGAAAGTCAGAGACAGTTCTATTCACAGAACTCTCCGTCTGATGTAGTGGGAGATAAC
GGACAATCTTCCTCTAGAAAGGAAATTAGAAAGAATGTTATTTGTTTTAATTGTAACAAGCCAGGGCATTTGGCTAGAAATTGTAGATCTGCTAGTAACAATTTGCCGATG
CTGGCTCTATTGAACCTAATACTGACGATGCTGAATTGATTTACCCTTTCAGTTAAAAGAGGCTTTGTGTAGTGCCCCTGCTCTTGGGCTCCCTAACTATGATATGCCATT
TACTTTGTTTTGTCATGAAAYACAGGGTAATGCCTCTGGARTACTAACCCAGAGCCACGGAGAGGGACAGAGACCCCTTGCCTACTATAGCACTCCACTTGATTCTGT
GGCAGCTGGATTAACCCCTTGTTTGCGGGCTGTGGCGGCGGCTGCTGCCTTASTGGAATCAGCACAGGCCATAGTTCAGGGACAGGAACTGAAAGTGATGGTTCC
TCATGCAGTGAGTGCATTGTTAACCCGTCGATCGACACAATTTACTAATAGTAGGCTGACTAAATACGAGATTGTTTTGCAGGGTACACCCAATGCCACCGTCCACCG
GTGCACAGTGTTAAACCCTGCCACTCTTTTGCCAGANCCAGAGGAGGGACTCCTCATGATTGTGTGATTTTGACAGAACAATTACAAACTACTTGCGACCGATTGTAT
GAGCGACCTTTGACTGACCCTGATATCGAATACTTTGTAGATGGTTCCTCATTTATGGATATTGACGGCAGGCGTTGTACTGGGTATGCTGTAGTTACATCAGAAGGAGA
AGTGGAAGCTAATGCTCCCCACCTCACTGTTCAGCNCAGGTGGAAAGACTCCCTGTCCTCGAGGACCCTTCCAAAATTTGCAAATTGATTTCATTGAAATGCCTAGAT
CTCTACATTATCAATATGTACTAGTGGTGGTGTGTACATATTCCCGCTGGATAGAAGCCTATCCTTGCCAACATGCGGACGCTCTAACAGTTGCTAAAAAGTTGCTACAA
GAGTTCATTCCTAGGTATGGAATTCCTCAGCATATATACTCTGATCATTTTACTGCTGAGATAGTGCAGGAGATAGCACATAGGCTAGGAATTGTGTACCATTTACATTGTC
CCTACCATCCACAAAGCGCAGGAAAAGTAGAACGCGCTAACGGCACCTTAAAGAATAAGATTAGCAAGATATGTGCCGAGACAGGTTTAAAATGGCCAGATGCATTA
CCTATAGCCTTAATGTCAATAAGAAATGCCCGACACTCAGCTATTAAATGTTCTCCTCATGAACTCATAATGGGTCGACCCATGATAGCTGCTAATACTCTAGAGTTACTA
AGACATGATCATGCTCTAACTGATGACACAATAGAAATGTATTGTAAAAAACTAATAAAATCTCTTGAGCAATTTAAATCCCAGGTTGCTGCTGTGCGCACGAACACCGA
GGACGAAAAACCAATTCATCACATCCAACCAGGTGATTACGTGCTGGTAAAGCATTTCCAACGGAAAAAGACTCTTCAGCCCAGGTGGAAAGGTCCATACCAAGTA
CTACTGATCACTAAAACAGCTGTAAAGGTAGCTGAAATTACACCGTGGATTCATTGCACTCATCTTAAATTGGTGCCAAAACCCAATCCACGCTCCTAAATTAACCATTC
CATCGGGCACAATAATCCCAGCTGGTTAACCAACATAAGACGAGATTGGCAAGAGCCCCTCGTGAAGATACGAAGAATGAGGTCATCAGGAGGTGCAAAAATGTTT
GAATATATTATGGTTACCTGTTTTATACTAAGTGTATTTGTTATTTGTTTCTTTTTTCTTTGCAGTCCTTTTCAGGTGTTCAGATCAGAACCTACTAAAAATATGGCAGAGAATAA
GATACATTCACGAATTAAACGTGATCTGCACGCTTCTCCTTGGGAAAAAGATAATACCTTTGTAGAAATGTTGAGAAAGGTGTATAATGCTTCTAATGTTTCGGAGCCTTG
TTGGGTTTGTGGATATATTCCGCATACTCAGCAGAGCCCTTGGATAGGAATACCTTTTACAGGAGATGATTTTGGAATAAATTTTAATAGGAGCTACACTATCCGCACAAG
TAGAACTAGTGCCCCTATTAGTTCTGAGAGAATACCATTAATGGATGTAGGACCAGCGGGTGGGACCCTATGGATGAGGCGGAATGAGTCTAGAGGTGCTCCCCGTA
AAGAAGTAGGTTGCCTATCCAGATTGTTAAATATGACAGGATATAACATCTGGGAATCAGTAGAGTACGGAAATACCAGTGATTGGGTCGCCGCACAATTTCTTGCAGTT
TATGCCTATACTGCTAATCTGATGGTATCTAATGAGAAAACTCACATCTCATTACAAACATTTGGAAACAATCTGCAAGTAGGGATTTATTGGTTGTGTGGAAAATGGGCCT
ATAGAGTATTGCCATTAGGTTGGGTAGGATGTTGTTTATTAGGAAGAATAGTACCCGCAGTTACAATGCATAATACTTTACCGGCAGGTAGAGTTAGAAATTATCGGGATG
TTTCCTCTGTATCCAAGACGCACATTAAGTCAGGACTTTGGGAAGAACAATTTTGGGGATCCTTATTCCCTATGGTGGGTGTGGGTCTATTATATGATAAAGTTAATGCTTT
AGCTGATGTACTTGATGATATGCTTAATCATACTACTGCTGCGATAAGGAAACTAAATGATGAACAAAGACAAATAAGAATGCTTGCCTTGCAGAATAGAATGGCATTAGA
TTATATATTAGCTTCAAAGGGAGGGGTTTGTGCGCTTCTAAAAGATAAATGTTGTGTTTACATTGCAGACAATAGTGTATCCATAGAACATGATATGGAAAAGGTGAGCGA
ACTTCAGGAAAGACTTAGATCTAGGGAGACTACTTGGGATCCTTTTGGTCTGGGCGGACTATTTGGTTCACTTGGAGCAAAGCTGATGTCTGGTATGTTTTTGTTGCTTG
TGGCTTTTGTATGTATCTATGTAATTGTTGTTTGTTGCAAAACACTTATTCAAAAATGTCTCGGACCAACAAGAGCAATGCCACAGATACAGGAGGAGAAATATGATGATG
ACTTCCCACTTGATGAGTTTCCCCTAGAGACTTTGTACAGTGACATGTTTGAAGATCAATATGAGAAGGAGCAAGGCTCATGGAATTGGCCAACTTATAACTCCACTGG
ACTCTGAAGAAATTGATTCTTACAGTGAGTCCGTAAGAATAAAAAGGGGGGACTGTGGAAGTTGAAGAATTTTGCTGATTCTGGATGTCAGCACAATGATCTTTACCTCA
TTGCTTATTCTGCTGATAGCAGATAAGATAACACTACAATCACCTGTCTACCTCATTTCTTAGTTTACTTATGTCTTGTTTGTGTCTCGGAAGAATGTGTACATATGGTATGTTT
TCGTATGTTTTCTTTGCTCTAAGTGCCTTGTTGCAAAAGAATCAGAGTCCGCCACGACAAAGGACAAGTTGTCCCTTTCTTGACCTTCTGTAAGTAAAGATTGCACATGT
CTGGGGGGGGGGTTATTTTTCTTCACCTCTTTAAACCGTGTGTGTCAGCAAATCAAAGTAAATATACCAACAGATGGCTAGACAATATGCTAATAACTGTTTGCGCTTCTT
TGCCTATATATTGTTAGAAAATTAAGGAGGGGGAGAGAAGTCTGCGAACTGGACTCTCCCCTGTGCGCACTTGAATAAAGAAGCTCTTGCTTACTCAACTCATTGAACT
GGTGTCGTCAGATTTCTTGCCGACATTCACA

AnERV3

TGTGATAGAATTTATCTTGATTCCAAGGCTCAAGCTTATAGGTGTCATATAATGATATCCTATATTAAAATTGGCTAACTTGGACTCAGGAAATGCTTGATGCTTAAACAACA
CTTAAGACCATATATGTCTAAGTTCCGGGAGCAGAGAGCCCCCACCTAAGATGTCTCTGCTCTAATCATTTTGTTCTCTCATACCTTGAAACTTATTTGTTATGGGTACAT
GACGCTCATCCCTATGTCTTATCTGTATTCTCTTCCTATTACTGTTCTAAGAGTATTTGCACGTAAGCACTTACTTTTTAATGTATAAAACCCAGCTGATGAATAAAACGTCA
GAGGCTTATTTTGAATACCAACAGGTGTCTGGTGTCTAATTCTCGCTTCTCCAAGTGCACTTGTAACAACAATTTGGGCTTCCTCTGAATATAACAAAGATCAACATTTCT
GATCCACAACAATTTGGTGCCGTGACCCGGGATCTGACGGTCGAGAGACTCGGACACTAGCGTACAGGTGGACGTCCCTCTGACAGTGAAATCAGCTGTACAGGA
AGGTAAGAAAAACTTTTTTCTACCTTTATCAATTTCGCTCTCAGATCCACGTCCTACTAGTCTGTCTGGGTAAAGAACGACTTAAAAACGTCTGATCTAAAGAACTTGTTTT
CTGTGGTATTATGGTTATATTCAGTTGGAGGTTGTGTTATGTCTTGTGAAAGGAATCTTGAGATAGTGGAATATAACTGTGGGTTGTAAGTTATTTTTATTGTTTGCCTTGTCAT
TTGTTTTATAATTTGAGTTAGAATCCTCCAAGGGCAGATCTCCTGATCTGTGTTTAGAAGACAAAACTGTGAAAATTCCACTGCAGCGGTCCTAAGTGTTTAAATACACTGT
TTTTGTACAGTTTGTCTTGTGTAAGGAAGTAATTAACCCTTACAAGTTTTAAGATGGGTAGCTCTGAAAGTACCCAAGGGGGATTTACGGCTTTGACTATTGTCAAGCATC
ACAAAGGACAAGGAGCAATTAAGAATGTTAAGAGGTTACTTAAGAAAGCTGGCATGCCCTCAGAAGGCACTCTAGACCCACATATATGGAGTGAATTTATGGCAAATTA
CAGAGGTTGGTTAGAAGATAATGACTTAGTAGAAGTCGCACAAAAATGGCATGATACTGCTGTTGAATTAACTAAATGTGGTTGTAGAGAAACCACCAGGAAAGGTGTA
CCATATTATCAATGTTTTGGACCTCTAGGGTCCCAGAGTAGGCAAGATCCTGGGAAAAATCTTGCAAACCCACCCTTTTACCAGGATACTGCAGGGACTCCTTTGGTC
CCAAAAGACAAGGCGATTTCCCTTTTACAGAGATCAAAACCTCTCTTTCCAGTAATATTACGCAGTGGGACTTCATATAGTGTCCCAGTAAGACCTATGGGTTCATTAGA
GCAGGAAGGTAGTCCAGACTTAAAGGAACCCCTAGTTGCCCCCACAGATACCCCAGTGGGGCCATTAGCTCCACAATTTTCTACTCCCATGATACCCAAAGTTGAG
AGTGTATACCATGAGCACAGACAGCCGAGCTCAGGTATCCAATACTGGGAAGAATATAAGCCCTGGACACCCAGTGAACAACTGGCAATAATACAGCAGCTGCCAG
ATGTTGAAAAGGCCCCCTTAGCTCTAGTCCAGTCTCTATCCACAATTCAAGCTACTTACCAGTGTACTTGGATGGATTTAAAACAGATTGTGGCACTAAAAGCTGGTCC
CATTTTGGCTAGAATAATTGCTAAATATACAGATGCCCATGAAGATAACCTCAGAAAAAGCAACATGCAAACAGATGAATGGTTTAGAACCAAGTCTGCATCTGGAGCAT
GCTATGTCACTGCAATGAGAGAATGGGCTAAGGTAAAGACAGAGGAACTAGCCCATAAATTAGGAGACATACTACAAGGGAAAGATGAATCTGTAGAAACATTCTTCA
CCAGACTACAAATGGCTTGGGAGGCATTGGGTTACACTGATGTTGTAACCACTTCGACCAGACTGCTTATTAACTGCTTCATGGATGGTATCCACATACCCATAAGGAC
TGGTCTCCAAAATGCTAGACCTGAGTGGAGAAGTTTAACCTATGGTGATCTGGTGAAGGTGGCTAGGGGAGTGGAAGCTAATCTAAATACTAGAAAGAGGAGCACAC
CTCTAATGGCTGTAACTGTCCCTAGAGCTGATGGGGACAGAAGGCAGGTAGACTGCTGGAATTGTGGCAGGAAAGGACACATTAAACGTAACTGCAGGTGTCCTCC
TAAAAATAGAAAACCCCAAGATTCTAATGATAACTCCCATGACAGCCAACCACCACCACCACCACCTCCCCCCCCCCAATAGGAAATTGGCAAACCAGTGTCCCTA
CATATCATCAATATCCCTGACTCTGCAGGACCCTCACCCATTATCAATGTTACACTGCCTGGGGACCGAATAGTCCCTTTCCTGTTGGACACTGGCGCTGCTAAATCA
ATGTTACAGGAAAAATATGTATTCCCGTATGAGAAATCTAATGATGTATTATCATGTGTAGGGGTAGATGGTATTGAAAATGACATTCCCCTGACCCACAAGTTGAAATTGA
AGCTACATGACTCTACTGCCCTTGTAACAAGGCTACTTGTCAGCCCTCATGCCCCTGCTAATCTACTAGGCAGTGATATACTGTCTGCTCTTAGTGCAAATATCCAGTAT
AGCCCCACTGGAAAAGTAACATTGTCTGTGCCTTTAGAGGACAAACAAATGATATCTCTTTCAAATGTTCTGGCAACTCTCACTATGCCCAGCCATGTGCCAGAACCTA
CCATTGACTTGTCCTCAGTCCCTGATTCCTTATGGGCTACAGGGAAGACAGATACAGGCAGACTCTCTGTGCCCCCAGTCAAAGTGAATCTCCTGCCCGGAGCCAA
ACCTCCCTATCAAAAACAATACCCCCTAAGCCCAGCACAGGAAGATGCTATTTCGCAATCTGTGCAAGAGTTTCTGCAGGCAGGGGTAGTTGTCCAGACCACCTCC
CCTTGTAACACACCATTGTACCCAATTAAGAAAAAGCAAGAGAAAGGGAAACCTGTGGTCTACCGTATGGTACATGATTTGAGGGCAATTAATGCTGTAACTGAACTCA
TTGCCCCTAATGTTCCCAACCCACACACCTTGTTGGCACAAATACCACCTTCCTCTAAGGTATTTACTGTTATTGACTTGGCAAATGCATTTTTCAGTGTCCCCCTGGCT
CCCGACAGTCAGTTTTTATTTGCATTTACACATAAACACAAGCAGTACACTTGGACAGTATTGCCACAGGGTGCACAAAACTCACCCACTATGTACAGCATGGCTATG
GCCCAGATTCTTGAAGAATGGCAAAGGCCAAATAACGTAATTTTATTGCAGTATATCGATGACCTACTCCTATGTTGCGATAGTATGGAATCTTGTATTGTAAACACTCTTT
CCTTACTAAAATTCCTAGAAAAAGCAGGCTGTAAAGCAACAAAAACGAAACTGCAAGCATGTGTAGAGAAAGTTATTTTCCTAGGACATTGCATTTCCCAGGGCATAAA
ACATCTCACGGAAAGTAGAAAAACTGCAGTTCAAAATATTAGACCCCCCCGCAGTTTAGCCCAGGTTCGTACTTTCCTCGGTTTGATTTCTTACTGCCGCCCTTGGCT
GCGTTCCGTAGCTATCTACATGCAACCACTGTATGATTGTCTTGCCAACGAAAACAAACCATTTAAATTAACCCATGCGGCAGAAAATGCCTTCTATACACTTAAGATAC
TGGTAACGTCAGCTCCGGCACTGGGTCTCCCAGATTACAGAAAGAATTTTAACCTGTTCTGTACTGAGCAATCAGGTTATGCTACATCTGTCCTTACACAGGATCATGG
TGGTAAACAAAGACCAATAGCATACTACTCCACCAGATTGGACCCTGTTGCCAGGGGTGCCCCCTCATGTGTTAGGGCAGTGGCGGCTGTTACTGCCCTCCTGGA
CAAAGCCACTGATATTGTTTTAGATAATCCTGTGACCATATACACACCGCATGATATCAACAGCATACTCTCACAAGTTCAACCAAAACACATACATACTGCAAGGCATC
TCAGAATGCAATGTGCCTTGCTTATGCCACCTAATGTTACTCTTAAAAGGTGTATGGTGTTAAATCCATCTACATTGCTTCCCCTGGATTTAAAAGAGGGGAGTGGGGAT
AGTGTGAATGGTTGTACTTCTGATCTTTTTGTAAATTCACTTTCACCAGATGAGTTAGCTGATGCCCAGGTACCCCATGATTGTGCAGCACTAATGGCACAAGAGACAGC
AGGTTTTTTGCATGTATTTGATTCTCCATTATCTAATGCAGATTTTGAGTTTTTTGTAGATGGTTCCAGATATGCTGATGAGAAGGGTAATTTCCACACTGGATATGCTGTAGT
CTCTGCACATGAGGTAATAAAAGCTGAACCACTCGCTCCTCACTGCTCTGCACAGGAGGCAGAGCTGAAGGCACTCGCTGAAGCGTGTAAAATGGCTTCTGGTAAG
ACTGCTAATATTTATACAGACTCTCGCTATGCTTTTGGTATAGCCCATGATTTTGGCCCTATATGGCGGGCTCGTGATTTTCTCACCTCAGCAGGAACTCCAATCAAGAA
TTCTGAAGCCGTGAGTGCACTTATGGACGCTATCCTGCTACCCACTCAAGTAGGCATCATCAAGGTGGCAGCTCACGTGAAGATCACAGATGAAATCTCAAGAGGAA
ACCACAAAGCAGACGAAGCTGCCAAGCAAGCAGCAATCAAAGTTCATCCCTCTACAGGTAATACCTTTCTACAAGTTGTTCTTACAGATGACGCCGTTGACTACAATA
TACTGTTGAGCTTACAAAGACAGGCAAGCAAGGAAGAACATGACCTCTGGAGGAAACAAGATGCAGAACCTGATGATGACAGAATGTGGAGGAAAGGTAAAAGACT
GTGTTTACCACGAGTGTTGTACCCCATGATGGCCCAGATTGCCCATGGACCTACACATCTATCCAAGATGCACATGAACAGCATAGTAAACACCCATTGGGTAGCAC
CCGGATTTACTGTTGTAGCAGCTAAACACACCCAAGCTTGCATGATTTGTGCTAGAAACAATCCAGGACAAGCTGTGAAAACACCAAGTAAGCACACACCAAGGCC
ACTTTACCCGTTTCAAAGACTGCAGATAGACTATATTCAACTACCTAAGGTAGGAGTGTATGAATATGTGTTGGTGTGCATTGATCTCTTTTCAGGTTGGCCTGAGGCCTA
CCCAGTGTCGAAAGCCACAACGAAGGTCACAGCAAGTAAGCTTCTGAAGGAACTGGTATGTAGATATGGAGTACCTGAAGTGATTGAAAGTGACAGAGGTAGCCATT
TCACTGGGGAAATAATGCAAGAAATTATGAAAGCCCTAGGGATCAGTCAGGCCTTCCACACCCCCTATCACCCCCAGAGTAGTGGGAAAGTAGAAAAGGGCAAATG
GCACCTTAAAGAACAAGATTGCAAAAGCCATGCAAGACACAGGAAAGCCCTGGACTGAATGCCTTCCCCTAGCACTCTTTTCTATGAGATTCACCCCCAATTCCAGA
CATGGCTTATCCCCTTTTGAGATCCTGTTTGGATGTGCCCCAAAGACAGGCCTCTATTTTCCCCAAACACTACAGATGCAGCATGGTAGTATGACTGCCTATGTTCAGG
CCCTACACCAAAGATTAGGCAGGGTGCATAAACAAGTCTTTGATTCTATTCCAGATCCAGACTCTGATTCTGCAACTCACAAACTGCAACCTGGTGATTGGGTGGTGAT
AAAGAGGCATGTGAGGAAAGGACTTGAACCACGCTTTGACGGTCCATTCCAAGTGCTCTTGACTACGGCTACCTCAGTGAAACTAGAAGGAAAAGCCACGTGGATA
CACGCCAGTCACTGTAAACAAGTCACTCCGACTGACCCACCAACATGAAGCTGTTCTTATCCTTTACCATTTTATACTGTGTTCAGATATGCTACTCTGACTTGAGACCA
GAAAATAATAATGCATGGCAGGAAAATATATGGTTGCAGATAGCAAAGGCTTTTAATTTAACTTCCTTTTGCCTCAACCCTCTCCTCAGTACTGAAAACGCCATAGAATCT
TGTCTTATAGCTGTCCCTACCCCCATCTCACTGTTAAGAAAGTACCTACATACCCAAAGACCCCATAAATCTCTACAACTATATTCTTATGCCTTCAACCCACTCGTAGG
CAAGCCTCCTCTAGCCCTGAAGACAATTAATGTGACAGACGCTGAACTATGTTGGTCTTTTTCATGTAATTGTTCAAGAGACCCTAATACTGAATCCCTGTTTTGCCCCA
CATGGTGGAAAAATACCACGAATTGTAGAGCCCTAAGCACCACTATGAATTGTACCAAAACTCACCAAGTTATGAGTTACCACTACAATGTCTATCTTCCTAAAGGATGG
ACTTTCTCCTGTGGAAATATTACTTATGCTTATGTCCCAGGTAACATTACAGGAGGTCCCTGTGCAATCTCTCGCCTAACCATGACCATGTTTTCCAAATCTGACCTGATC
ACTAAAACCACCCAGCTACACAAACTGCTGAGAGACAGGAGAGATGTTAAAAACACCCTTAATGCAGATTGTAACCCAGAAATCACTCTCTTATCTAAAGCCGAATAC
TCTGCCCTAGCGTACACCTTGGTAGGAGTCCCAGGTTTAGCATTGTATAATACCCGAGTAGTAAATGCTGTAGCCTGTTCCCTTGCAAAAGGTTTAAATGCCACTTCAC
AGGCCCTAGAAGAGCTACTTATGGACAGCCAGGCAAATAAGAAGGCAATCCTTGAGAATAGAGCAGCCATTGACTACTTACTAATGAAACATGACTTAGGCTGTGAAG
CTATAGAAGGTATGTGCTGTTTTAACCTATCTGACCATGGCACTATGATCCATAAACATATAAAAGATTTACATGACTTAGTACAGGAGATCAAACAGGATACTGGCTTCTT
TGGGGATTGGGATTGGACATTTGGGCTAGGAACATGGTTGACTTTACTAATTAAGAAGATTTTGTACTTTTTGCTGTTAGCTATATTTGTCATTATGGCTATTTTTCTAACCTT
TAGATTTTTTTCATGTCTACTAACCTCTCTATGTAAAAGCAAACCTACTATTGGTCGTACCTTAACTTACATTCCAGATCCGACTTCGAAAAGGTTTTATGATGTTACCTCGC
CTAACAAATGAAGGAACTCTCCTCTGCTGATTCTGGTGGTCTCTCAGTCCAAGGGAACCGTGGTGAAGCAATAAATGAAAGTCCCACAGGCTTTGGCTAGCAGGCCT
GGATATACCTTGGACATTTGGGTGACCCTTGGATCAAAAGAGGGGATTGTGATAGAATTTATCTTGATTCCAAGGCTCAAGCTTATAGGTGTCATATAATGATATCCTATAT
TAAAATTGGCTAACTTGGACTCAGGAAATGCTTGATGCTTAAACAACACTTAAGACCATATATGTCTAAGTTCCGGGAGCAGAGAGCCCCCACCTAAGATGTCTCTGCT
CTAATCATTTTGTTCTCTCATACCTTGAAACTTATTTGTTATGGGTACATGACGCTCATCCCTATGTCTTATCTGTATTCTCTTCCTATTACTGTTCTAAGAGTATTTGCACGTA
AGCACTTACTTTTTAATGTATAAAACCCAGCTGATGAATAAAACGTCAGAGGCTTATTTTGAATACCAACAGGTGTCTGGTGTCTAATTCTCGCTTCTCCAAGTGCACTTG
TAACAACAATTTGGGCTTCCTCTGAATATAACAAAGATCAACATTTCTGATCCACAACA

AnERV4

TGTTAAGGAAATTATCCAGCTTCTACTATGGATATAAAAGTGATTTTCATTTTATCTGTATACATGATATAATTTGCTGACATTTGAAAAGAAAACTATAGAAGAAGTGAACTGT
GTGTGTTAAGATAAGATAACATTTGTACTACTCCTTGTAATTTTGCCTCTTTTGCTTAAAATTCTAAACCCATTAGTTAACACATTTTTTGTAAAGTGAAACTTGAAAAATAACT
AACCAATCAATAGATGGATTCAAAAGCCATTATTGAAATGTCTGTAAACCAATCATGTTTTACTTTGAGGGGTTACACCCTGTGAACTGTGTATAAATAAAGGGTGCAGGA
CACTTTCACTTTAGGGTCCTGACAAGGTTTTGAACTATACTCAGACTTCTGTCATTTTCCTCCGTCGACGTCATTCACCTGAACACGAGGCAACGATCATCACGTGGCT
GACCTTTGGCCTGCCGCAACATTTTCTGGTGGAGACTGCGGGCACCATAACAACACTCCTAGCTGCAGATCGCGACACGGAGACCAGGACCCAGGAAGACCGC
GGTAAGTAACCCATTGTGTTACTACCTGTCTCCTGAGCTCTGCTCGCTGTTCTGCAGTGTGGTGTGTGAACGTGGTGCCTGTACCGTGTTCTGGTGAATGTAATATTTCC
CCTATATGTGCTAAACTGTCTTTAGGCGTATGAATGCACGAATGTGTGAGTGGCTGAGATAAGCTGAATCTTGGAGGGGTCTGTCCCTGCTCCTCCTGAGCCAAGCTG
AAAGTCTAGAGCTCGCTGACTGGCTTGTCCGGTCCCTGTCTCTGGAGGCCTCTGGTTCTGGTGGAGAAAGGGTTGACAGACCTATACTGGAGGAGGCTGTGAGAAG
TGGCAGTGTATAAAGCTGTCCAGTAAGTTTGGATAGGGGCCCTGCAGTTCCTTGTTGGTACCTGAAAAGGCGCCTGAGGTCAGGGGTTGTTCCAATGTTATGTCATAC
TGTGTGGTTTCACATGTACTGAGTGCCTGCTGATATTGTGTGAGGAAGAATTACTGCATTGCTGCCAATTCCTGTGCTGTACAGTGTATACATGCGTGGGTATTATGATTC
CAGTTTGGATTGGGTGGTTGCAGTTATAATCATTTGGATTGCAGTATTTGTTTGCTTCCTGTGCCATAGAAAGAACGAAGAGGGGTTTAAGGGGTTTCCAGTGAGTAACC
CCAGAGAGGACGTTGAGTTACTCTGTGTGAGGGCAAGACTATATTGTAGTATTGCCAGACCAGTGGATAGTGCTGCCTCGGAAGTGTGCCCACAGCACCACTGGTAG
GAAAATTGGTGCCGCCTCGGAAGTGTGTCCACGGCACCAATAGTAGTGCCGCCTCGGAAGTGTGTCCACGGCACTACTGATAGTGCTGCCTCGGAAGTGTGTCCA
CAGCACTGTTGTAATTCACCAGGTGTGTGAGTACCTGGATAGTGAATTACAAGGAGCCACCCACGGGATGGGGGCTACACAGAGTAAGCGTTCCTCTGAGAAGGGA
GCAGAGAAATGGGTCTCTGCAATTGATATTATGCAATGCGCAAAGGGCACTGAAGCAACCAAAGACTGTAGACAAATGCTAAAGAAATTAGGAGCAGGTGTGAATGG
AACATTGGATGAACCAGTATGGCAGAAATTTCTTACAAATTTTAAGGGGTGGTTGGAAGACCATAACAAATATGACCAGGCCACAATGTGGTATGATGTAGCCCAGGGA
ATGACTTTGACTGCCCATAAGGCAAAAGACACTGACAAAGGGACTATGTATGAAATATGGGATGAAATTAAATTGCCTACAAAAGAATTAAACATCCCTTCTGAAGTACG
GTGTATTATCCCCCATAACCCACAAAAATCTCTCCCTCCCACAGCACCACCTCCATATAAAAGCCCGGAGGGGTGGACTTGTCCTAATTGTGGACAACAGAACCCA
GACACAAGAGTGAGATGCTCAGCCTGTGGGGGGTCCAAGCCACACTCTCGTGTGGGCTTATACCCATTAGTCAATAAGACCCTCCCCACAGTGAAAGCTGGGCAA
CTGGAAACACATGATTATGAGGTCTATAGGCCATGGACTCCTAGTGAGTTACTGGCTATGTGTAATACTTTGCCAGACCCCCGTACTGTCCCCAGTACTTTCCTTCGTAA
ATTGGAAAGTATCCAGACCGCATATAAAGCCACTTGGGCGGATCTGGAGAGTCTTCTAGAGACTAAAGGAGGGGAAGTACTTAAGAATCAAATGATGGCAAAAGCCC
CTGGTACTGTCCCAGTGGATAAGGCCACAGACCAGTCAGGCCAGGAATTTATCATGGCCCTTAAACCCTGGGTAAAAGAGCAACAGGACAATCAGACTGACACTTT
TATGATGCAAACACAAGACGCTAAGCAGACTGTAGCTGACTATGCCACAGCTTTAGAACAGCTCTGGCTTGACTATGGGTATGAGACAGGAGACAGTAAGGACATGA
GACTACTAAAACATACATTCATGTCTGGATTAAAAGCACCCATACAGGATGCGGTGAAGAAGGCTCGCCCTGATTGGCGGACCTGCCAGTTCCCTGACCTAGTGGC
CATAGCAAAGGGAGCTGAGCTAGACTTACAAAATAAAAAGAAGCCAATAATGGCTAACCAACCTTTTAAACCTAAGGGCAGGTTTCACGAAAGAATACCAGGTGGGG
ATCAGTCTCCACGTAACTGGGACCAAGTAACATGCTGGAATTGTGGGAAAAAAGGGCATTTTTCAGTCAAATGTCCCCGGAGAGGTCAGGCTACTGCCCGGACCTC
TCAATGAAGTGACGGCAATCCTGTGAGTTCCCTCAACGTTAACTACCCCCTTGACGCAATCCCAGGTCCACAACTAATCATCACATCCACATGTGAGGACACAGGGA
ATTCAATTGACTTTTTGGTAGACACAGGAGCAGCCCATACCCTAATACAAGAAAAATATGTGCCTGAGGAGCTTATCTCCTCAGATTACATCTACTGTCAAGGGGTTGAG
GGGACAACTGTTAAACTCCCACTAACAAAGCCTGTWAAACTTAATATTCATGATGCTGCATCTATTGTCACGCGTGTGTTGGTAAGCTCAGATAGTCCATACAATTTGCT
GGGGACTGACGTACTCAGTGCCATGAGGGCATCTGTGGATTTCTCTTCAGGAAAACCCCAAATTACAGCCCCTGTGTCTGAAGAGACTTTGTGTGCACTTATGTACTT
AATGACATTTAGTGCCCCCTCTGCCTCTCAGAATACACCCCCAGATTGGCTGTCTGAAGTGCCATCACATTTGTGGGCAACTTCCAAGACAGATGTGGGCAGACTAC
CAGTAACTCCGGTGGTTGTGCAGCTGAAACCTGGCATTAAGCCGCCATGTGTGCGTCAGTACCCATTAAATTCTTCACAGGAAGATGCCATTGCAAATGATCTGGAA
GAGCTGTTGGGGAAAGGGTTAATAATCCCTATAAAAAGCCCCTTTAATACACCCCTATTTCCTGTAAAGAAGAAACAGACTGTGAAGGGTCAGCCTCCAGTCTATCGTA
TGGTACATGATTTAAGGGCAGTGAATGCTGTAACTGTCATAAATACACCACAAGTGCCCAATCCACACACACTGTTAAACCAAGTGCCCATAGATGCCACATCTTTTTC
TGTCATTGATCTTGCAAATGCTTTCTTCTCTGTCCCTTTAGATGAAGAATGCTGGCCCTACTTTGCTTTCACTTTTCGTCACAAGCAGTATGCATGGGTCGTTTTACCACA
AGGGGCTGCCAATTCCCCTGATATTTTCTCCACTGCTTTACACTCTGTGTTGCAGTCCTGGCAGCCGGCCTATCCGGACACTGTTCTTTTACAATATATTGATGATCTGT
TGCTTTGTTGCCACAGCTCTGAACAGTGTAAAGAAGAAWCTTTATCTTTGCTTATTTTCCTTGCTGAAAATGGTTGTAAAGCCTCCAAGGAGAAGCTGCAACTTTGTAAA
TCTACTGTGATTTTCCTAGGTCACTGTATTTCTCCAGGTACCAAACACCTCACGACTGAGAGGGTTACTGCTATCCGGCAGCACCCCCTGCCAAGTACTCCAAGGGC
TCTGCGTGCCTTTATTGGACTTGTCAGTTACTGTCGGCCCCTGGATTAAGGATTGCTCTCAGATGATGCAGCCACTATATGATTGCCTCAAAGGAGAGACCTTTGAACTC
ACACCAGGAGCTGTGAGTGCATTTCATACATTAAAACTCACTCTCACCACGACACCAGCATTGGGCATCCCCAATTACAAACAAACTTTCACGTTGTATTGTCATGAG
CAAGGGGGATTTGCCTCAGGTGTACTGACGCAGAAACATGGGGATCGTATGCGGCCTATAGCATACTATTCTACCAGTTTGGATTGTATTGCCAGAGCACTACCCAC
ATGTTTCAGAGCTGTCGCTGCTGCCTCTGAATTGGTGCTAAAGACGGAAGACTTGGTACTRCAACACCCTCTTACCTTACAAGCACCACATTCAGTTGCTGAATTACTA
CACTCCACCAAACTGAAACATTTGTCTCAAGCCAGGATCACRAAATATGAGATGACACTCCTCAGTCCCCACATTACCTTACAGCGATGTACTGTTCTAAATCCAGCTA
CTTTGATGCCAGTGGATTTAGAAGAGGAGGGGGGAGTGGAGTAGTAGACATGTCTGATCCAGTTGCAAATCATAACTGTCATGAATTGATGATTGCAGAAACAAAAG
GGTTACCAAATGTTACTGAAACACCATTAACTAATCCTGATGCTGAGTATTTTGTAGATGGCTCCAGATGGTGGAATGACGAACAAGGACACTTTGTCACCGGGTATGC
GGTAATCCAGGATGGTAAGGTAATCATACAAAAACCACTCCCATCTTGTGCGTCTGCACAGGAGGCGGAGTTGAAGGCACTCACTGACGCATGTACAATGGGTAAG
GGACTGAGAATAAATTGTTACACTGATTCCAGGTATGCTTTCGGGGTAGCACATGATTTTGGTGTTATCTGGAAAACTAGGGGATTTCTGACTTCCTCGGGCAAACCAAT
TAAAAATTCAGAAGCAGTACAGAGCCTCATAGAGGCCCTAGAGCTGCCCACAGAGGTAGCCATTGTTAAAGTGCCAGCCCATGTGAAGCAAGATGATCCGCTGGCT
GTGGGAAATGCTCAAGCAGACCAAGCAGCCAAACAAGCAGCCGGTCTGCCGCTCCTGACATTGTTTTTCCAACTATGCCAGATTTGTCGCTCTTGCAGGAATTACA
GGCCCAGATAACGCCTCAAGACAAAAAGGTATGGATGGAGGCGGGGGCTACGAAGACAGAAGAGCTCTGGCAGGTAAATGGCAGAACATGTTTGCCTCGGAGTT
TGTATCCGATGATGGCCAATTTAGCACATGGCCCTACTCATTTGTCGAAAGGAGCTATGAGCGCACTAGTGTTAGAACATTGGTTTGCACCTGGAATTCAGACCCTCTT
TGAAAAGTTTGTTCAGGGTTGTCTTACATGTGCCAGACATAACCCTGGGAGGTTTGTAAAGACCCCCAAGAAACACACTCCCAAACCCTTGTACCCGTTTCAAAGAGT
GCAGGTGGATTACATCCAGCTACCCAAGGTGAGCACATATGAGTATGTCCTTGTAGCCATTGATGTATTCTCAGGCTGGCCAGAAGCTTGGCCAGTAACTACAGCTAC
AGCCAAAAATACTGCTAAGAAAATATTGTCAGAGGTAGTCTGCAGATATGGGATTCCAGAGGTAATAGAGTCAGACAGAGGTACCCACTTCACTGGGGAGATAATGGG
ACACATACTGAAGACCCTAGGGATCACACAAGCCCTGCACACTCCCTATCACCCGGAAAGCAGCGGAAAGGTGGAAAGACTTAATGGGACAATTAAATTAAAATTG
CAAAAGTATTTAGACTCACACAGTTCTATGAACTGGGTAGAAGCTTTGCCAGTAGTATTATATGCCATCCGCACTACTCCTACTGGACAAGAAAAGCTTAGTCCGTTTGA
AATTTTGTTTGGTTCTGCACCAAGGACAGGGTTATATTTTCCACAGCAGCTGCAAGCAGACTACTCTACCCTGACTTCATATTTGGTGGAATTGCATGAACATCTGTCAA
AGGTGCATTCACGTGTTTTCTCTTCTATACCAGATCCTAATTCTCTTGAAGGTACACATTCTCTAGTTCCTGGAGATTGGGTAGTCCTGAAGAGGCATGTAAGAAAGACC
CTGGAGCCAAGGTTTGACGGTCCATATCAGGTGCAACTGACCACACCCACCGCCGTCAAGCTGAACGGGAGACCAGTCTGGGTCCATGCAAGTCACTGCAAAAA
GGTTCTGAAGACAGACAGTCACGAACAGGAAAGCAGCCCTACAGTTGTAAATCCAACCAGTTGTGAGAATAACGTGGATCAGTAGACCATCACAATGCATGCAATGC
CTTGCCTGTTTCTTTATATTCTATGTTTTTGTGCGCAGGTGAAAGGAGGAAAACATACAAATTCCATTTGGCAACTACATCAACAGACTGCCAAGCAGCTGAATACTACT
GACTGTTGGATATGCTCTCATATCCCTGCCAACTACAAGGGAATCCCTCTGATAGGTGTTCCAATCTCTATGAATGATTTGAATCTCACAGACTATAGCTATGATGTTCAA
ATAGATGTAGATCACACCTCACATAATGCTGTTTCTGATCAAGGGATATACTTAGAGATTACTGGTATAATCTCTCCGCCCACTATATGTTTGGGGCCTATGTTTGTGAACC
ATATAGCTAAGATAAAGGGCCGTTTTGTCAGTTTACCACAGATATATGTGGGGGAAACAGATTGTAGAAATGCTACTCTGATATTCAATATGGAATTTAGGAAGACTTCATG
TGACATAGTTAATACTAACTGTAATAGTTTGTGTTTCTGTCCAACTATGAAAGTTTCTTGTGCCCCAAGGTGTGCCTGTCCAGAAAAGAAGGACAAGGACAAATGGACTT
GTAAGACGAAATTGTATGATAATATGATATGGTTTCAGGGATGGCTGGGACATCACGGCAGGATGATTCCCAAAGTCCTGAAACAAGGACTATATTACATTTGTGGTAAC
AGAGCCTACTCATGGCTCCCTATGGGGGCATGGGGCAACTGTACCATAGGAAGAGTTGTCCCAGCCATCAGACGACGCACAAACATTTCACTTACTGATGCCCCTG
AAACTCTTCTTGGGATGACTAGAAACAAACGAGAGTTATTCACATCAGCAGATAAAGCCTGGATGTGGTTCCCTTCTTGGACAGGTTGGGGAATTGAGTTAGCAAATAA
ATTAAACAAATATGCAAATATCATGGATGGAATTATTAATGAGACCACAAGTTCAATCCATGCCCTTAATGAGGAACTGGCACAAGTCAGGAAAGTGGCTCTCCAAAATA
GGATGGCCTTAGACTACCTCCTAGCTATAGAGGGTGGGACTTGTGCTATCATAGGAGAAGAATGCTGCACCTGGGTTGAAGATTCACATGATCCCATAGAGGCCCAC
ATGAATAAGGTAAAGGAATTGCAACAAAAAGCTAGAGACATAGCTAAAGAAGGATGGAACCCCTTCTCTTGGATGGGGTCTATAGGTGTCTGGTTCAACAATATGATCT
CTGATTTACTGAAGCCCATAGTGGTAGTGATAGGCCTGGTTTTGATACTGCTAATTGCTTGGAAATTACTAATGTTGTGTATTTCTCATTGCAGTAGGGGAAATTCCCTCCC
TTGAAAATAGATGACACAACAGCCCTGGAGATGACAGTCTGAAGCTCCTGTGCTGCGATGTATCCCTAGTTGCTGTTGTCATCCACTTAGATAGTGTGTTTGTGTGTAGT
ATGTGGTTTTAATAAAATGTATGCTTAATCAAATGACAACTACTAGACAGGCTGGAATCGATGCTTGCATCGCATGGTCTATGCAGCTTATCTGTATACTCTGTGAAGTACG
GCTGTCTAAGGGGTCATTTGTTTGTTTAGGCAGGTTAGTAGTTAGCAAGACACACATGATACAATGAAGCCTTCAGAAGGAAGAAAACGTCTACCACGTGAACAGAAA
AATGAAAAAAAGGAGGGAAATGTTAAGGAAATTATCCAGCTTCTACTATGGATATAAAAGTGATTTTCATTTTATCTGTATACATGATATAATTTGCTGACATTTGAAAAGAAA
ACTATAGAAGAAGTGAACTGTGTGTGTTAAGATAAGATAACATTTGTACTACTCCTTGTAATTTTGCCTCTTTTGCTTAAAATTCTAAACCCATTAGTTAACACATTTTTTGTAA
AGTGAAACTTGAAAAATAACTAACCAATCAATAGATGGATTCAAAAGCCATTATTGAAATGTCTGTAAACCAATCATGTTTTACTTTGAGGGGTTACACCCTGTGAACTGTG
TATAAATAAAGGGTGCAGGACACTTTCACTTTAGGGTCCTGACAAGGTTTTGAACTATACTCAGACTTCTGTCATTTTCCTCCGTCGACGTCATTCACCTGAACACGAG
GCAACGATCATCACGTGGCTGACCTTTGGCCTGCCGCAACA

AnERV5

TGACATGGACATTCGTTGTACTGTTCATTTAAGCACAAGAGACTCCATTTTGTTCTCACTGTTGAAATGTGTTCAGTAACCTTCACCTAACACACAGACACATCTCCTCTA
GAAACTCTCTACTCCCCCCCTCCCCTATCCATGTTTTACTTTTGCAATTGTTCTATTCGCTAGCTTTTAATAATATATTTCTATTTGTTTAGCTTTTAATAACGTATTTCTATTTGT
TAGTTTTTAATAACATCTCACACCACCCCTTTCTTATCTGTGTATAAAATAACATGTAATAATAAATTTCGTCAGAACGGACATTTTAAACACAGTGATTGATTCCTGTGAATC
TGTTCCTGCAGCTGCAGTATTAACTTTTATTTTAGAGTCCCAATCAGAAATTAGTCATAACA
GTTTTGGCGAGTCAGATATCAGGAGTTTGGCAGAGTTTCCAGAAGGAACCGCGATTCGATCGGGATCATCGGGAACGCAGACCTCTAAATGCCGGCTAGGTAAGCT
GTCCTTAAGCTTGCTATATTTCTGCCTTTCCTATGTATTCTATCTGTCGGTTCTCAAGGAAACCAGACCACTCTCGGACCTTTCTGGTTGGGTAACGTGTGTGTGTGAATG
TGTTTGTGTCCCCCTTCACGGGTTGGACTGTGTGTGGATAGGGAGAATCCTCTGCAGTTCCCTGCGTTGACTCCTTGTGGGCGACCTGGGTCAGAGGCGCATGGTA
GGGTCAGATAAAACTCACGGAGAAGAGAAGAGACGAGGAGGGTCTCTTAATGAGTGTTTTGTCTATTGTCATAAACCCCTTCTATCCACGTAGATAAGTGGAGACTATC
CTTGTAAATCTGCAGATTTGCTGTACTATTTGTTTTAGAGGCAAGAGGGTATAGGCACTAGTAGAGAA—GACTGGCCAGTCATT
ATGGGCATTAAAATAAGTAAGGGAATGAAGGGTAAGAGGCCTTCAGATAATGCCCAGCGCAAGATGAGCGCTAGAGAATATATGAACCGAAGGTACGGTTCCGCCTA
TACTGAGCCCCTAGATAAATGGGTAGAATGGACAAAGGGAACAGCTAAACCTTTAAAATCTGAAGGGACTTTTGATTTACAAGTATGGCAAACTTTTCTGCACGATTATG
GCCCACTACTGTGCAGTGAAGGAACGTGGAGAATAGCTTGCACATGGGAGACAGAAGCTAAAGTAGCAAATCAAACTGGCTTGACGGAGATGTTTAATAAAAGTACT
GCCCAAATTTACTACGTTTGGCCAGAAGACTCTGTAGAAGAGGACCCCCCACCTTATGTGGCTGCCAGTTTGGCTTTAAGGGGTCCAAGCACAAAAGCAAAAGAAT
TAGATTACATGGAACACACAAAAGATGAGCTGGGCAATATAGTAGAGAGCAGAGGTTTAACAGCCCCGTATCAACCTACAAAGAGGGATTTGTCTCATATCCTAGAAA
CGGATGATTGGCATAAATCAAAAGGCACATATGCAAAACTAGATCTCTCTGAACTTAAGGCTTTGGCCAGAGAAAGAGAGATAGATTCTGAACTTCCAGGGCAAACCA
TAATTTCCAAATTGTGGCAGCAAGATGAAGAGAGAAGGAAGGGAGAAACTGCCCCTTCTGCCCCGGTCACAGAGCAATCTGCTTCAGTTTATCCTGTCAGAACCCA
ATTAGTTTTTGAAGGTAATGCTGTTGACAGTAAGTCTCAAATTAAATGGCATGTACCTTTCAGTCCCCAAGATATGAGAGCCATATTAGAGGGATTGGGAGACCCTAGAA
AGAATCCAATTGGGTTTGCAAACAAACTTAGTGCTGCACAGGAAGCATATGAGGCTTCTGCTACTGACATTCATCAGCTTCTTATTAAAGCAGTGGGAAGCAACATGTC
TGGCTATATACTTAAGGAATGTGGTGTAGACATCAAAACTCTAGGTAAGATGGTTAGTGGACGGGAGGTTTGTGGGAAATTGAAAGAAAAATTGCCCCAAATATATGGAC
AAACCAGACATGCTGAATTTATGAATGCTAAGCAAGGGAAAGAGGAAGCAGCAGGGGCTTATTGGAGGCGGTTAAGCGATATGGCTGAGGAAGCAGGCATTAATATA
GAGGAAGAGAAGGTGGGAGAAACTGCAATCCATCTACGCAACTGGTGAAACAAAGGTTTCTTGATGGTTTAATACCCCAGTTGAAAGAGAAGTTGTTCACAGCAAGG
CCTGAGGCAGGCAATATGTCTATTCGAGACGTCCTGCCTATCCTGTTATCCATAGAAAACAGGATTAACCAAAAAGACAGCAAACCCAAGGTTATGTACATGGGGAAG
CAGAGAGGAGGAAGAGGAAGAGGCAAAGGCAATTTCAGAAATCATGGCGATCGGGGACCAATCATATGCTACAATTGTGATGGGGAAGGACACATAGCTAAATTCT
GCCATCTTCCTGACCGTAGACTAGAAAAGAGCTTAGAAGCACCTCATGCTGATCCCAGTACAAAAGCAGTGGATCAGGCATAGGAAAATGGCATACCAGTATCAATC
ATGTTAAATAGAGGAGACAATGGTCCCGAGACCAGAGTAAAAGTATTTTTACAAGAACTCCCCTATCCTGGAGGAGAAACAGAATTAACCTGTTTAATAGATACTGGTG
CAAGCCGAAGTGTTTTTAATATAGAGGACTTACCTCCAGGATTATTATCTACAAATGTTATCTCTGGTGTTGGCCTAACCGGGACCTCTACTGATTTGGTCGAGTCCAAG
CCACTTAAACTCAGGCTAGGACAGCAAACAGAAATTGTCACTAAAGTGTTAGTTTCAGACACAGTTCCTACCAATCTTCTGGGAGCTGACATTTTGAACCAAATTTTAG
CAAACATAGACTACACTCCTACAGGAGTCCAAGTGGTCAGTAGTCTGTCAGGAGAAGTTTTAAATTCTGCCTGTGGAGTATTTTTAATCCAAGACTCCACTCCGCCATTT
CCACCAGAATTGAAAGTGATTCCCACTGAATTATGGGCCACCAGCAAATATGATGTAGGTCTGCTGGATTGCACCCCAGTCATGATAAAAATAAAACCAGGATCCCAT
CTCCCACAGATTAAACAGTACCCACTGAGTATTGCTCAGACTGAGGGGATAAAGGATCAAATCACTCAACTGAAAGCAGCACGGGTTGTTGTACCAATTAAGTCCCAA
GTCAACACACCCCTGTTCCCAATTGCAAAGAAGCCAAACAAGAAGGGTGGGCAGGTGACATACCGCATGGTGCATGATTTGAGAGCCATTAACAATATAATTGAGG
CAGATACCCCGATCGTGCCCAACCCCCACACTTTACTAAGTGGAATACCTGCCTCCAGCACCTACTTTACAGTGATTGATCTTGCAAATGCTTTCTTTTCGGTGCCCC
TGCACCCAGATTGCTGGCACCTTTTTTCATTTACCCATGATGCCCAAAAGTTAGCATGGACAAGATTGCCCCAAGGCTGTGTGAGTAGCCCCTCTGAATTTAATACTG
CGTTAAAGCAAGTATTGGACCAATGGTCCCCCTCACACCCAAACACAGTTTTGCTTCAGTACATAGACGATTTGCTTATTTGTTCTGATACAAGGGACATTTGTAGCAAA
GAAACAGTAAGCCTTCTCATGTTTCTCTATAACAGTAACAACAAAGTTAGCAAAGATAAGCTACAATACTGCCAAGAGAAAGTGATCTTCCTGGGTCACTGCATTTCACA
TGGAATCCGTCACCTAACCCCAGATCGAGTTAAAACTCTACAAAAATTTGAAAAGCCAAGAAATGTACGGCAACTACGTGCCTTCCTGGGGTTAGTGGGATATTGTAG
GGACTGGATTCCGAATGCATCAGAACTTTTGGCTCCTCTATATGAGGCTACCACTAGAAATCCTTTCAACCTCACATGGGAAAATGAATTGCAGATGAAGGAGCTTATT
AAAATTTGTCTGAAGCTCCCGCCATTGGACTTCCTGACTATACTAAAACTTTCTCATTGTTCTGTCATGAAGTCAATGGATTTGCTGCAGGTGTACTCACCCAGCTACAT
GGAGACAAACAGAGGCCTGTGATGTATGTATCGGCCTCATTAGACCCCGTCATCAAAGGCTCCCCCAGTTGTATGCGAGCCATAGCAGCCTGCATTTTGCTGCTAG
ACAAAGTAACTGAGCTTGTACTAGATTCTGCTCTGATCCTGTATGTACCACATGCAGTACAGGAAATCATGAATCAAGTAAATACCAGACATGTGTCTGCCAGTAGATTTG
ACAAGTACCAGTCTGCTCTTTTTGCTCCATCGAATTTAACAATCAAACGATGTGTTACTTTAAATCCGGCAACTCTTCTTCCTATAATTGATGAGGAAGATGCAACAAATG
GTACTGATGTAGAACCCGAGAGTTTCTCTCATGATTGTATGGCTCTGATGGAGGTAGAAAGGGCCTCTGTCACACCAGCAAAAGATGAGCCTTTATCAGATTGTGACAC
ACACTTATTTGTAGATGGTTCCAGATTTTACACTGATGATGGTCAGCCACACACAGGGTTTGCTGTAACCACTACAGACACCATTTTACATCAGGAAGCATTGATACCCC
AGATGTCAGCACAGGAAGCGGAGTTGTGTGCTGTAGTCAAAGCATGTCATCTCCTAGAGAATCAAAGGGGAAACATTTACACTGACAGCAGATATGCATTTGGAATAG
CTCACGATTTTGGGCCTATCTGGAAAGCACGGAATTTTCTGACTAGTGCAGGGAAGCCAGTGAGACATGCAAAACTCATAGAGCGGCTATTTGAGGCCTTTCAGAAA
CCAGCAGAAGTTGCCATCTTGAAGGTAAAAGGGCATGCAAGGGGAAATGACATGACAGCCAGAGGAAATGCTTTGGCGGACCAAGCGGCAAAGGGTGCTGCCCT
TATGGCATTTGGAGTCACACCCACTGTAGGGGTCGCCCAGGTAGCGGGGTCAGATTCCCCTCATCAAGAGCAGTGGGATGTAGAAACTCTAATTCAGTTGCAGAAC
CAGGCCCCTGATGAAGAGAAGGATAAGTGGATAAAATGTAATGCTACCCAAAATCAGCAGGGCCTGTGGAGCGCTGACAACAGGTGGTGTCTACCCAGGAGTTTGT
ACCCAGTTTTAGCACAACTGGCACATGGCGTCAGTCATCTTGCTAAGGGCGCGATGGTGGCACTTACTGACTGCTATTGGATAGCACCTGGATTTAGTGCATTTGCAG
CCAGGTTTTGTGCTTCATGTCTCACTTGTGCTCTGCACAATGCAGGAAGGCCTGTCCATGTTCCCCAAAGGCATTCTCCTAAGCCAGATTTCCCTTTCCAAAGAATTC
AAATAGACTACATTCAGATGCCCAGAATAGGCACCTATGAGTTCGCTTTAGTTTGCATTGACATGTTTTCGAGCTGGCCTGAATGCTGGCCAGTAGCCAAAGCAAATAC
AAAAACAACAGCAAAGAAAATTTTGAATGAAATAGTTTGTAGATATGGAGTTCCTGAAGCTATTGAGAGTGACAGAGGAACCCATTTTACTGGGGAAGTAATGAAAGAG
GTAATGGAAGCCCTTCACATACAGCAGGCTTTCCATACACCTTATCATCCACAAAGTAGTGGTAAAGTTGAGAGACTTAATGGAATTTTGAAAAACAGATGTGCAAAAAT
TAAAGCAGATACGGGAAAAGGATGGGTGGAAGCTTTGCCATTAGCCCTATATTCTGTACGAACTACCCCTAAGCAACCACATGGGCTGTCCCCCTATGAGATTTTATTT
GGAGGGCCACCCAAAACAGGGCTATATTTTCCCCAACAGTTGCAGGCTTCTCACTCTAGTCTGTTAGAATATGTTCAGATTTTGCAAATAACATTGAATAAAATACATGA
AGGTGTTTATAAATCTATCCCCGACCCAGATTCTGTTTCAGGTATTCACAGAATCCAGCCAGGTGACTGGGTAGTGGTCAAGCGACACGTCAGAAAGCCACTAGAAC
CAAAATTTGACGGACCACATTTGGTGCTTTTGACGACCAGTTCCGCTGTCAAGTTGGAAGGAAAGCCCACCTGGATTCACGCCAGTCACTGCAAGAAAATCCTGAA
GTGAATGATATCCTTTGAAAAAATTATGATAATGTTTCTTTTCTTTTTGTTGTTTGTGAGAACTGAGACATGGGAAAATAACCCTTTCATCCAAATGTTAAAATTGTATGCAAC
CCATAGCAACGCTAGTAACTGCTGGGTGTGTTCTCACCTAAATCCCCAAACAGGAACAATACCCATGATAGCAGTGCCTCTCAAATTCACTGATCTAAACAGGACCA
GAATAAATGGCTCATACATGGTCTCCACAGCTAATGAGCACAGATCAGTGACTTTGATGGTTTCAAAACAGATAGGCACTCCACAGATTTGTTTAAACATTACATGCCCA
TTTAAAACTAAATGTACACGGTTGGGCAATTCAGATTGCACCGGAGCAAAAATTACAGTTTATGGAGGAAACTATGGTCAATATTATCATATCCCAGATCCAGAACTAAAT
AGAACCCTTTATAGTACAAATTGCTCACAGTTCCCAATATGTCAAAATATCGGTTACCTGGGGTTCCAGGTTATTGCTTCCAGTTATTTAGAAGCAGGCATACCATCCATA
CGTAGAGGGTTATATTATATATGTGGAAATAAAGCTTACGCCTGGCTTCCTGCTAATGTTACAGGTACATGTTATATTGGGAAGTTAGCACCTGCTTTGGCTGTTCGCAAA
GATACAGTGCGTCCAGATCAAAGTCCAGAATACCCTTTAGTGTTCAAAAGAGAGCTGTTCACTGAAGGTGACATGGCATGGTCATGGTTCCCTTCCTGGACAGGATGG
GGAATTGAAGCAATGAAAAGAATAAATAATTACACTAGAATTGTAGATGAAATTATTAACCTTACAACCACTGATATAGGTTTATTAAGTGAAGAAATGGCTCAGATTAAAAA
AGTAGTATTAGAGCACAGATTAACTCTAAATTATCTCACTGCAGCTCAAGGTGGGATGTGTAAGTTGTTTCGCACTGACTGTTGTATATACATTTCTGATAATGAGGACATTA
TTAAAGGGCATCTTGCTAAGATAAGAGAACTACAGAATAAGGCCAGAGATATTGCTAAAGATGGATGGAACCCCTTTAAGGGTTTAGGAGGTGTTGGTGATTTTTTATATA
GCATAGCAACATGGTTACAGAAACTAGGCGCTTACATTGTGATGTTCCTATTTCTCATATTTGTCATTTACTTCCTCATCAGGCTGTGCCTCTGTGTGACTACCCGCTGTA
TCAACAAATTGTCCACCAGCCCTGATGAACCCGTTGTTATGAGAAAGCTAGGAGAGAACTGAACAGGACGCCGCCTCTCTGCGCCAATGTAACAGGAGTTACCAGC
AATCGGCTGAATGTAACGCGGAGAATAATGGTGTCAAATAAACAAAAGGGAGGAAATGACATGGACATTCGTTGTACTGTTCATTTAAGCACAAGAGACTCCATTTTGTT
CTCACTGTTGAAATGTGTTCAGTAACCTTCACCTAACACACAGACACATCTCCTCTAGAAACTCTCTACTCCCCCCCTCCCCTATCCATGTTTTACTTTTGCAATTGTTCT
ATTCGCTAGCTTTTAATAATATATTTCTATTTGTTTAGCTTTTAATAACGTATTTCTATTTGTTAGTTTTTAATAACATCTCACACCACCCCTTTCTTATCTGTGTATAAAATAACAT
GTAATAATAAATTTCGTCAGAACGGACATTTTAAACACAGTGATTGATTCCTGTGAATCTGTTCCTGCAGCTGCAGTATTAACTTTTATTTTAGAGTCCCAATCAGAAATTAG
TCATAACA

AnERV6

TGTATTACAAATAAAAATCTGCCAGTTTTTATATAACACTAATGGTTTATTGAAAGTAAAATTGTATCATAGGAATCTATCTGTCTTTTAATTGCTGAYTTGGATGTCACTGTGCT
GACTCAGACATTGGGAAATATAGCTSTCTCTCTCATGAAGCCATTGTTTATCCCTTAGACAATGTGTACGTTTTCCTGTCACCCTCTACATCCTCGTTTATGTCTTCCCGTG
AACTTTTATTGGGGAAATGAATGTTTATGCAYATTTTTCCCTCCACTGTCCATATCCTGAGTATTGATTGTTGACTTCACACAATMCATTGTTAGAAGGCTGACATCTTAATTT
AAGACTTACGAAACTAGGGCGGGAATTTGGTCTTCGGAAGTCTGAGYTTGCAAGCTAGGTTGACCCTAAGGTCAAGTAAGGAAATGGCCATTCCCTGCTTGATTTTGG
GTAATTTATGTCTAGTCAGCAAATAACTGCTGACTTGYCTCCATAAAWAGGGGTTTCGGAKTGCGGTGTAAGACAGGAATTATTAACCTGTGCTTTGAGACGTCTTAGCA
CAGTGTGGCTGCYACGAATAATATCCTGTCTTTTCGCCGTTATTCTGAAGATTCCTCCGGCTGATGCTCAAAGACAGATGATGACAGAAGCCCCCTCTGTTTCGTGGTG
AAGTGGTGTCAATTTCCCATTTGAGTCTCTCTCTTGAAAGATGATGATACAAAGCTCATGGTGATGTAAGTTTTATGTTTATGTATCTATATTGCTGTACCTTTGTTTGCTTATTT
AGGAGTGTATTTGCYGTGTTCTTTGTATTCTATATATATATATATATACATATTATTTGTAATAAAATCATAACTTTGTTGCAATTTGATCTCTTTAAAACAATATAAAGAACCTGCA
TTTATATTAGTAGCCCTGCCCTGGGCATTAATTAGTAAAGTATTAGATATTTGGGTGTGCTCTTGGCCACCAGCAAAACATTTGGCGTAGTCGGCAGGATATTGTGGAGA
TCAAAGTATATGTATATATAGAATATATATAGAGGAATATACTCATAATGTCTTTATTTAAGCGACAAAGTGTTGTGAAGGGACCAGTGGTAATACCAGGRTGGGAGCAGGC
TCCTTATCAGGTGTTAGCACATGAATGGTCTCAATGTGTTGGTTTAGGTGAGTCTTGGGACTCTTGTTTAAAGAACACAGAGCCCTCTGATGTAGTAGAATGTTTACAAAG
AGTTAAGATTGAAAAAGGTAGAGAATTTGCTTGTGTTAGYAGGAGGGGGTGGTTGCTGTTAACAGCATATCGCAGGGAAAATGAGGAAAAGCACAAGCTACAGGAAGA
GGTAACTAGATTAGAAAAAGAACTAAGTAAAATTAATGGCACTTTGATAACWACTCAGTGTCAAAATAAGTTGTTRATGGACAAACTTGATAAATATTCCACMATTGCAGA
AAAAGCAGCTGTGAGAGTTGCACAGTACAAATACAAAAAGAGGCGAGGTAAAGTAGATAGCCGAAAGGTTAGAACTTTGATAGCCAARGCTGATGCTGATTGGGATYT
TGATCACTGGGATGGGGATATTTGGAATTCCACTAGTGAAGATGGGGACACTTCTGAGGAGGGAGGWATAGAAATGCGCCCTGCTATGAGGAGGAAAGAGGAGAAA
ACTATTGAACCAAGGGGACCTGATAATCAGGGTAATCCTCGCCCCCCCAGAGAAAAAATAATAGGTAAGGAAGTAATTGAAGATTTTACGCAGCAAGAAATTATGGATT
TGTTATCACGTTTTAAACAAAATCAAGGAGAAAGTTTAATTAAGTGGATGGTTAGGATTTATGATACTGGAGCTGCAGGTATCATGTTAGATGAAAAAGATGCCCCTAAATT
TTTGACAGTGAGTACTGATCCCCATATACAGACAGCTTTTAGAGAWTACCAAGGGGGAAATTCTACTTTATTGGCAATTGCTGGTGCTGGGTGTGACAGGAAGTATATTT
CAGAAACAGAATGGCCCCCTAATGATAAACCTTGGTTCACCCTGAGGGATTGTATACAGCGTATTAAAGAGGAAAGCATGAAAATATCTATATACACTGGACATAGTGAT
SAGATGTTATTAGGCCCCTTTGCGTTAGCAACACGCAACAAAATCATAAGGACTGCCCCACCTGCTTATAAATCTGTAATAATGACYTTGTTGGTGAATCATGTAGGTAAT
CCAGTAGGGGAAGTAATTGARGCAGTAAGACAGTTAGGAGACCTGGGTGAATGGGGTCCTAGTCAGGGGGATCGTGGAAGGGATATGACACGGAATGTTCCCCGA
GAACGTCAACATTCGTCTTCAGTACAGGGATCCCGGGTAAGTAGAAGAGATATGTTTATGGCTTTGTTAAAGGATGGTGTTGCCCGTGAGGAGATTGATGGAATAGGGA
CGGATCAGTTGTGGAAATTGTACAAAAAAAGAGGTCTGAATAACCCAAAGAAAAGTACATCACAGAATAAACCTATTCATGTAGAAAAAGCAGTGCGAGGTGATAAGG
GATCACACTCTGAAGGAGGGCCCACCTTTGGGGAGTTRTCTCATACYCGCCCCACTGCTCCGGGTTATGAAGATCTSTATGCAGATGCAAAAGCTTGTTGTATGGAT
CCAAACAAAAAAGGCCGTCCCATTCCACTAATCCTTTTTATGGGGAAGCCAATACATCAGATTGACTAGCWATAAATAGTGTACAAAAAGATATTCGCCCCCATGTAG
ATGTAGTTATTAACTGGAAACACAGAAATTCACAACAAGTCAGGGCACTGGTAGATACAGGGGCTGAAGTAAGTTTAATTTATGGAAATCCCCGTAAGTTTAGTGGAAAA
AGAGTAGTTATYACTGGACTAGGGGGAAAACAAATAGAGGCAGTACARACTAAAATTAATATGAAGATAGGAAAYTTACCAACTCAAAATTACTCAGTAATGATAGTCCCA
ATACCAGAATTTATAATTGGAATTGATTTATTAAAAGGACTAACTCTTCAATTACCGGATGGTAAGTATGAATTTGGGACCCCCAGAGTAGAATCTTTTAATGTTCATAATCT
GAATGTGAATACAGCCACAGTACAGGTAGACCCTGTACTAGTAGGACATGCCAAAATACCTCCAGTATCTATACCACCTGCTACTAAARTTGTCAGTCTGAAACAGTATA
GGATTCCTGGGGGTGACAAAGAAATAGAAGAAACTATACAAGAACTTTTACAGGCAGGAGTTATAGTGCCAACTACTACACAGTGGAACAGTCCTGTGTGGCCAGTW
AAAAAATCAGATGGCACATGGAGAATGACTGTAGATTATAGAGAATTGAACAAGAATACTCCTCCCTTGACAGCTGCAGTGCCTGATACAATTACTTTAATTGAACAGATT
CAAAAACATAGAGGCAATTGGTATGGGGTCATTGATTTAGCTAATGCTTTTTTTACAATACCCATAGCTCCCGAATTTTGGGATCAATATGCTTTYACGTGGAAAGGAAGA
CAATTCACATTCACCCGTTTACCTCAGGGGTGGGTACACGCACCTACCATCTGYCATAGGGTAGTGGCTGAATGTCTAGACACCTTTGACTTGCCCAAAAACATGCA
AATATCACATTACATTGATGACATTATGATACAGGGAGAATCACAGGCAGAGGTAGCAAAATATCTACCAAAGTTGATTGAACATATGAGGAGTAAAGGGTGGGAAATCA
ATCCTGCCAAAGTCCAGGGACCTTCTCAAGCAGTAAAATTCTTAGGTATTCAGTGGAATAGGGGACATCGAGAAATAACTGATAAGGCCAGACAAAAGATCTTAGAGT
TTGCTGTCCCTAAGACAAAGGTACAAGCTCAGCAATTTATAGGACTATTTGGGTTCTGGAGACAGCACATTCCCCATTTAGGACAGATTCTAGCTCCTCTATACAGGGT
GACAAGGAAGAAATTTCATTTTGAATGGCTCTCAGAACAACAACAGGCATTTGAAAAGGCCAAAAGTGTAGTACAACAAGCACTTGATTTATGGCCTGTTCTGTCCAAT
GTCCCAATTGAAATTAATGTGTCAGTGCAGGAACAGTATGCTAATTGGAGTTTATGGCAGAAACAAGGGGGGAAAAAGGTGCCACTAGGTTTCTGGAGCCGTAARTTA
CCTGATGCTGCGCAAAGGTATACTCCTTTTGAAAGACAGCTACTGGCYTGTTATTGGGCTTTAATAGATTCTGAACAATTAACTTTAGGACATGAGGTGACTATTAGGCCT
GAAATTCCCATTATGCACTGGGTACAGAGTAATCCAAAATCACATCGCATAGGTCATGCCCAAGAGGCCTCAATCATAAAATGGAAGTGGTATATTCAGGACAGGGCA
AAGCCTGGTATAGCAGGTATAAGTTCCTTACATGAAAAAGTTGCCAATGTCCCTCATGAGAATCAGACTGAACATGTCCCTGAAGAAAGAATAATTCAGGATTCCCCTG
TAAAGTGGGGAGTTCCTTATGAACAGTTAACAGACAAACAAAAGAAACATGCTTGGTTCACTGATGGTTCAGCTAAATATCAGGCTGGAGAAAGAGTTTGGAAGGCGG
CTGCCTTCCAGCCAAGCACTGGAAGACTTATGACAACTACAGGTATAGGRAAAAGTAGTCAATATGCTGAGTTATACGCTGTATTTAGTGTACTGAGACAGGAAAAATTA
GAAGAGTGCCACATTTACACTGACTCCTGGTCGGTTGCAAATGGYTTAGCTACCTGGCTTCCAACTTGGGTAAGTAACAGCTGGAAAATCTATAGCAAAGAGATATGG
GGAAAGGAGTTGTGGCAAGATATCTGGGAAATTGTATCACATATACCTGTAACYGTTTTTCACGTTGATGCTCATGTCACCACAGATTCCCTGGAGCGATTGTACAATTC
AATGGCTGACGAGCAAGCCAAAATTGCACTCACCAATATCGTGCCTGAGGTGGAATCTTCATCTTTACAGGGKTTTACTAAATGGGCTCATCAAAAATGYGGACATCTC
GGAGAAAAAGCTACCTATCAATGGGCCATATCTCGAGGAGTGCCCATCAGGTTAGACATGGTGAAAGATGTTATAGCTCATTGTCCTGTCTGCCAACATAGTCAGCAG
CGACCAATTCCCCATGTGGTGAGAGGACAATTAGCTYGTGGTAAACTCCCTGGTCAGATTTGGCAAATGGATTTTGTTGGGCCCCTACCAATGAGTGGTGGGTGTCAA
TATTTATGTACAGCTGTAGACACATACTCTGGGTACTTGATTGCCCATCCGTGTAAGTATGCTAATGCTGCAGCCACCATACGAACATTAGAATTGATTGTCCTGTATTATG
GAATTCCTCTACAAGTTCAAACYGACAATGGCTCTCATTTTAAGAACAAAACAGTGTCACAATTTTGTGAGGATCACAATATTGAGTGGATATTCCACATTCCTTACTACC
CCCAAGCAGCAGGACTGATAGAGAGAATGAATGGTCTGTTAAAAGAACAATTACATAAACTTACTAATGAACAAATGGGAGTTTGGCGTAAATATTTAATTGATGCCTTG
CAAATTCTGAATAACCGCCCTATCGGGAATCAAGAAACTCCTTTAATGAGGATGATTACGCCCAATTTAACTGTAATGCCCCTTAGAGCTGCAATGTCAGGAGAAACTG
TGGTTTATTGGCCTTTGAATGATCARGTAAAAGCACCATTTCGAGCAACCCCTGGCTCAGCTGGTTTAGATGTCCATGCCACAGAGGAAATTACCCTAGACCCTGGGA
AAATAGCTACTTTTAATACAGGGCTRGGAATGCAGTTACCTGCCCATCATTTTGGATGGTTAGCTCCTCGCTCAAGTCTAGCCTTAAGAGGTATTCAAGTCTTAGGGGG
AATAATAGATGAAGATTATCGAGGTGAACTTAAAGTTCCCTTACTGAATAATGGAACTGAATCAATCGATTTTCGTATTGGAGACAAAATAGCTCAACTTCTGGTCATTCAA
CAGAAATATTTTGAGTTGGTAAGGGGGAATAGGCCTCTTATAACAACTGCTCGAGAGACAAAACAGTTTGGGTCAGTTGATGACAAGTTTGTTAATGGGACCAAAATTTG
GGTTAAATCACCAGGCCTAAATTCAGGCCCACCTGATGCAGCAGAAGTAATTGCACAAGGTCATGATAATACAGTAATAATAGTTAAATCTGGAGAAAATAAGTGGTAC
ACTGTACCCTCACACTGGTGTTACTTACGAGAATAATTTTATTTTATTTTTAGGTTGTGTGGCAACTTGCAATAATTTCCATTTTGATCATCCATTTGGAATGTGTATCTATAAA
TCAATGGCAATCAACTCGCAGTGTGACAACCAATCAAAACTTTTGGGAAACCAAGGAAAATATATGGTCATATCTAGCCCGAGTCATGAAYATCAGCCACTTTTGTTTAT
CCAAGAGTTCATCGATCTCTAATACCTTGGAAACGTGCCTTGTTTCTATCCCCACACCTATACAAATATGGACTTCATTGTTGAGAATTGAAYTAAAAGATGCTGATCTAC
AAGAAGGAGATATAACTCAGCACGAATTATATAATCCTTATTACACATGTTTCAATTGCACAGACACTACAATACCTACAGATGATATGGTTATGATACATACATATAATATTG
CTGCTGCAGATATTTGTTATATGATGACCTGTAATGCTTTGCAGATCCCTAGGTGTAAAATGAAAATAATGCAAGGGCAARGTCTTAGCAATGATACACTTTGCAAAAGAC
AGGAAAAAGACTGTAAATACATAAATGTTACTAACATGAACTGCAACTCCACTTTGATTGCTCCCTCTCACACCAGATATGTAAAATTACCCCCTGGGTGGATGTTTTCAT
GTGGAAATGACACATATAATTATATTCCAGCTAATATTATGGGGGGGCCTTGTGTTATTAGTAGACTRGGATTAATGATGTATCCATTATACCCCTCCGARTCACACAAATTT
AGGAGAGATACTTCTCATCTATTACCTGTAGATTGTAACTATGATRTTACTTTATTAACTGAAGGGGAAATCATAGCATTAGGTGTTAGTTTAGTAGGTGTGCCAGGATTGTC
CATATATACAGCAAGACAGTTAAATAGATTAACTTGTTTGGTAGCAAAAACAATAAATAGTACAAGTATAGCCCTAGAGCGTATGTTGCTTGATCTACAAGGGTATAAAAAG
GCAATAATGCAGAATAGAGCTACTATTGATTATTTGCTATTAGTACACAATCATGGCTGTAAGGAGTTTGAAGGAATGTGTTGCTTTAATCTTAGTGACCATTCCAAAGATAT
TCAAAAACAGATAGATGATTTAAATGGTTATGTAAAACATATAAAGCAAGATAATCAAGGGTGGTGGGATTACTTTTTCTCATGGTTACCTGATTTAGGATATCTGAGACAAA
TAATAGGAATTATTCTTGGTGTAGTAATTGGACTTTTAGTTTTAGGATGTTGTGTTCAATGTGTACCTTCTTTTTGTGTTATGTTTCGAGTTTCACCCGAACCAATCCCTACAG
GATACAATGTCCATACTTTCAGATAAACCAAGGGCCGAATGTATTACAAATAAAAATCTGCCAGTTTTTATATAACACTAATGGTTTATTGAAAGTAAAATTGTATCATAGGA
ATCTATCTGTCTTTTAATTGCTGATTTGGATGTCACTGTGCTGACTCAGACATTGGGAAATATAGCTGTCTCTCTCATGAAGCCATTGTTTATCCCTTAGACAATGTGTACGT
TTTCCTGTCACCCTCTACATCCTCGTTTATGTCTTCCCGTGAACTTTTATTGGGGAAATGAATGTTTATGCACATTTTTCCCTCCACTGTCCATATCCTGAGTATTGATTGTT
GACTTCACACAATACATTGTTAGAAGGCTGACATCTTAATTTAAGACTTACGAAACTAGGGCGGGAATTTGGTCTTCGGAAGTCTGAGCTTGCAAGCTAGGTTGACCCT
AAGGTCAAGTAAGGAAATGGCCATTCCCTGCTTGATTTTGGGTAATTTATGTCTAGTCAGCAAATAACTGCTGACTTGTCTCCATAAATAGGGGTTCGGATTGCGGTGTA
AGACAGGAATTATTAACCTGTGCTTTGAGACGTCTTAGCACAGTGTGGCTGCTACGAATAATATCCTGTCTTTTCGCCGTTATTCTGAAGATTCCTCCGGCTGATGCTCA
AGACAGATGATGACAGAAGCCCCCTCTGTTTCGTGGTGAAGTGGTGTCAATTTCCCATTTGAGTCTCTCTCTTGAAAGATGATGATACAAAGCTCATGGTGATGTAAGTT
TTATGTTTATGTATCTATATTGCTGTACCTTTGTTTGCTTATTTAGGAGTGTATTTGCCGTGTTCTTTGTATTCTATATATATATATACATATTATTTGTAATAAAATCATAACTTGCAATT
TGATCTCTTTAAAACAATATAAAGAACCTGCATTTATATTAGTAGCCCTGCCCTGGGCATTAATTAGTAAAGTATTAGATATTTGGGTGTGCTCTTGGCCACCAGCAAAACA

AnERV7

TGTTAGGGGTAATATGAACCTGATGGCAGCCATCTTGTTCCTCGCAGCCATTTTAGAATGAATAGACTTTATTATACTGGGTTATTATGCAGTGAGAATATTATGCATGTTATGAGTTTCT
TTAGCTATTCGGCCTTGCCAATGTACCCTGGCCTAACGCCTAGAAGTAAGATAGCATAAGATAATGTAAACAAGAGGCTTTAGACACTTGGTTGTCTCTGCAGATGGTAGTTTGTTAT
GACTCTGTAAATATTGTTATGAGAAACTCATGTTCTAGCTTTTCCATGTATTTCTGCTCTGTATAACTGTGTAAAATGACGCGGTCCAAACGTGTCATCCCCTTAACCCTATGTGCTGTC
TCTGTACGCTATAAGACATGATTACAATGTTCAATAAACATGAATAGCTTTGGATCATAATGCTGCGTGGTCTGAATCTGTTCAGGGAGCTCCTTATCAGATAAATCTGCATCTGCCTC
TCGTGGTACCGTAGGCCCCAGGCTTTGGTCTGCGCTGACACCCCCCCGTTGGATTATACACCTTACAATTCTTGGTGTCAGGAGTGGGATTCGCAGAGGGTAAGTTATCGTGGT
TTTATTACGCTGCTTTCCTTCTGTGAATTTGAACCTAAATTTAATTGTTTGTATTTTTAAAAGGTGTAATATATGATATAAGGTTAAAGGGGTTCCTGTGGTTGAAGCACAGGTTTGCGGA
GGTTAAAGTCCTGCCGGCAGTTGTGATACCCCTCCAGACAGGTAATCAGTCCTGTGCTGGGACACGACAGGTATTGTGAGTCCTGTCGGTGATAAGCGCGCAAGTAATATATGA
ATGCCAGTTGACGGGAGGTTCTTTTAAGATAAGGTCCCCCGGGCAACTTGGCAGTACGTGAGTAAGTCTCATGTGTGTTGAAGTTTCCCTGATTTATTCCGTTATAGAGTTTTGGTAT
CCACTTGAGGCTGGTGTTGCTATCTGAAGCTTTGCTTTTGCTTATTTTGCTTTGTTTATTGTGCTGTGCATAGAACATATTAAGTGATTGTGCATTTTATTATTGCTGCTGCTAATATCGCT
GTAATTATTTTTGTGTAACAATGGGGTCTGCACACAGCAAAAAATGTGATAAAAATGCTGATCAGAAATGTAATATTAAGTACAAGACTGTGCAGCGTGTACCATCCAGTGCTGGTAA
AAATACTGTGCGTGTTACAAACCCTAAGCCAGTTAAAGGGGAAACTCCAAAAGACTTTGTAGCTAGAATTGCAGGACAATATTATGTAATTCCATTTGTGGATAACATGAGTGGGTG
GAGCGAATCCTTGTTCCCTCAGGAGCAAAATCCGTTCCCCCGTATAGGCTCCCTCTGTGAACCGCATGTAACTATGATAAGAGGGATGTGTTTAAAGGATCCTGCTTATAAGGGA
CCAGAGGGCGATCCTAAATGGGATAAGAAATACATGAAGGAATGTGGAGAGGTTTGGATAAAGCTGTCTGTGTTTTGGACTGAATGCAATATCCTGCCTCCACCATATCTGGCTGT
GCCCACGGCATCAGCGGCTGTGCCCATACCATCAGCCGCCGTGCCCACGGCACCGGCATTGCCCCCACCTGCCCCGACACTTCCCCCGCCTCATCCATCTGCACTGTA
CCCTTCGTTGTCTGTGCTTAAGAAAGCACACAGCGTCAAGGAAATAGAGCAGCTGGAAGATGTTGCTGTTGCCACTTTAAGTGCTAGAAAAGGGTTAACTGGTTCTCTTCAGAGC
AGTACAGATGCAGGAATGTCTGAAGGGGAGGGGGAGAGAGTGGAATGTCGGGCTCCTGGTAGTGTAAAGAAATTAGCTCTCACATTCCCTAACAGTGTGGGCAGAGAAGGGG
GGTTGCAGAGCGCTGCAAAGACATCTAAAGCTGTAAAAGTGAAAGCTGTGAAAACTGCAGCTGTCACAGGTCCCAGTCCTATAGGGAAGCATACGAGATCACACACGCAGGG
AGCAGACACTGATTTAGAGGATGATGTGAACATAGAAGCACCGTTTTTTATAATCGGGGACCAGACGATGATTCGTCCTCCCGACACAGACCGCTTGATGGATTGGGCATTAAAA
TGTCCTGATCCTGTAAAGCATCCGATGGGAGCAATAACTTATCTTAGACGGGTGACTAGAGATATGTGTCTGGGTGGTCCAGACTACAGGTTTTTGTTGAATGTGATCACTAAATACA
CAGATGTGAATTTTACTGAAGTGTGCAAGTTCCTGGATACTAAGTATGATGTTCCAGTGAATGATGATTATGTGTGGAGAGATGCGGCTGTTGTGAGTGTAATGTGGGAGGATGTTGA
GAAATTCTTTAAAAGTCAGCATGGTGATCGTAAAGATATTACGAAAGCTGGTGCATGTAAGCAAAAGAAGGGTGAACAGGTGTCTGATTTTCTGAAGAGGTTTGGTGATTGTTGGCG
ACAGGAATGTTGTTTGGGTACGCAAGGAGAGTTGGCACAACTGTATGTGCAACACTTGTTGAATAATATGCATTCGCAAACTGCGCAATTGGTGAGACAGACAATTTCAGGTATTC
ATGCAATGAAAGTGGAAGAATTTGAAAAGGCTATAAGAGAGAAAGATGCAGCAGATGTTTTTCGTGTGTATAATGAGAAAGGTCAGGAGAGTATGTTTAATGCGTCAGGTAACAGGA
GAGGAAGAAAGCAGAATAGAGGGCAGAATAGAAGCTTTCGAGGAAGGGGAAGAGACAGATTTCAGTGGAACGGTGATAGGAGACAAGAGGCACATAACTCCACAGACAGATC
TTGCTGGAATTGTGGGTCTTATGATCATTGGCAGCAGGACTGCCCACAGCCTAGAAGGGATAGAAATGATCAGTATGGTGGGAATAATAATTCCTCCCAGAATCCCAGCAACACT
CCTAGATATCAGATACCCTGGACTCCTTCCCAGAATCAGAATAGTACCCGCTAGGATTGCCAGCAGAAGGGACTGCTCACATGTTATACTGCTCAGTTGTCGTATCATTGGCAAG
ATTTACCCTTAATGTCTTTAGTTGTGCAAGGTGTCCCACATGATTTTTTGGTAGATACCGGAGCAACTAAATCTAGCATTTCTGCTAGAGAATACAATGGGCCAGTGACTAATACATGT
GAAAAATCTGTTGGCATTACAGGAGTCCCAATTTCGTGTCCCAGGACGCCTCCTTTGGAGGTGTATCCAAATGACAGCCCTTCAAACAGATTTGTTTATGCTTTTCGCATAATACC
CAATAGTCCTGTTAATTTGCTTGGAAGGGATTTAATGGCTAAGATGCAAATGTCTGTTAACATTGATTGCAAGGGAGGATTGCATGTTGACACTCCCTGGGATTCAGTTCCGTTGCTG
TATGCAGCTGATCCTGAACCTGCCCTTACTAAACAGTTGTACCTTGATAAACTCAGTTTACAGACTCTTGAGAATGTGTTTGAGTATGCCTCACACCCTGACAAAGTTGTGACACAA
GCTTTTTACAGACCTTGGATTACTGACCATGTTGAGAAAGAGATGTGGAAAGCACCAGTTAATGCAGTTGTACTTACACCAGATACAGTTTATGTGCAAACACCAGAACATTTGTGG
ACATGGAAATATGGCAGGAATATTGTAAAAGATCCTAAGGTTATACCTGTACAATTAGAGCCAGATGCATTCACACATGCTTCACATGTCACTAATGATGTGCCAGAGGCACTTGAT
GAGTTAAAACAGAGTTTATGGGCCACTGGATTTAATGATCCAGGTTTCATTTCATGCACACCATATAAAGCCACTCTCAAACCAGGTGCTAAACCTGTATACATTAAGCAGTACCCT
TTGTCTAAGGAAAAGGAACAGGGAATAGCTCCTGTAATTAAGGACCTATTGTCTAAGAAAGTTATTAGACGCACACTGTCACCATACAATACACCTATCAATCCTGTTCCTAAACCT
GGAGGTACAGAATACAGATTTACACAAGACTTGCGAGCAGTAAATGCTCTGGTGCAGCCACTTGCACCGATAGTGCCAGATGTTAATGCTATTTTAACAGCCATCCCAAGTACTG
CAACTTGCTTTACAGTCATTGATCTCTGCAGTGCATTCTTTAGCATCCCTGTGCATCCCGACACTCAACCGTTGTTTGGTTTTACGTATTTAGGGAATCAATATACGTTTAATCGTTTAC
CAATGGGCTTTATTGATTCCCCTGCCGCGTACTCAGCTGTAGTTCGTAAGACCCTAGAATCCTGGACACCTCCTGAAGGTTCCACTTTGATACAGTATGTGGATGATTTGTTGTTGT
GCAGTGTTGATGAACAGACTTGTAAAACTGACTCTCTCCATTTGCTTCAGCACTTAGGTGATGCAGGGCACAAAGTGACTTTGAGAAAAAATGCAATTTTGTCTTCCACAAGTAACAT
ACCTGGGTTTTCTTCTTTCAGCAGGCAAGAGACTGTTATCCCCTGAAAGAGTTAAGGCTCTCATGGTTATTCCTCGACCACAAACAAAGACTGCTATGTTATCTTTTCTGGGTATGGT
TACTTACTGCAGGCAATGGATACCAGACTGTTCATATTATGATCATATTTTGCGTTCAGTTACAGGTCAGCATGAACCAGAACAAATCCACTGGACACAGGCTCTTGATAAGGCCTA
CAACACCCTCAAGCAGGCTCTCTGTTCCGCTCCAGCCCTAGGTTTACCCAACTATAACCTACCATTTCACCTGTATTGCACAGAAGGGGGAGGTACAGCAGCAGGGGTCCTTG
CTCAGGAACACGGTAATGGGTACAGAGCGGTAGCTTTTTATTCAAAATTATTGCCAACGGTGGTACAGGGTATGCCAGCATGCTTAAGAGCTACAGCAGCATGCGCTATGATGGT
GGAAGCGGCACAAACATTGGTACTGTCACATTCTCTGATATTGCATACATCACATCAAGTCTTGCAAGTGTTAAATAATCTCACTACACAGCACATGACTGCACAGAGAAGGTCAG
GCTATGAAACAATTCTCACATCCACTGCAAACCTCACTATCAAATACATTGCACCCACAGGAGGCCCAGCACCTTTATTACATGCATTGATTTCTAAATGTCCTTTAGAACACCCTG
AGGACCACAATTGCATAGACACTGTACACACAGAGACGTCACCACGTCTTGACTTGCAAACAACTCCACTAGAGGAGGGAGAAGTTGTTTTTGTTGATGGGTCATGTTCTAAACC
TGCTGATGGTGTTTATCTTACAGGCTTTGCTGTGGTACAGTTACCCGACACAGTGGTTACTGCTGGTTCTCTGCCTTTTGTGTCAGCCCAAGCAGCTGAATTGGTAGCCCTAGCTC
AGGCATGTAGAGCGTTTGCGATGAAAGATGTGACCATTTACACTGACTCAAGGTATGCGTTTGGAGTTGTGCATGATTTTGGAGTGATTTGGCAGAACAGAGGTTTTGTTGCAGCTG
ATGGTAAGAGTATCTCACATTCTACACTGGTACAGGAACTCTTAGATGCAATTAAATTACCACATAAGCTAGCAATTGTTAAATGTAAAGGCCACAGTACTGACTCCACTGACGTTAG
ACTTGGCAATGATTTTGCAGACACCATTGCTAAGGAAGCTGCGTTTCGGGGTCCACCGCACCCTGACTATCTCAGCACTACTACACAGGAACAGTTACACATGGTAATGATTCCT
GCCTTAGCATCTGATGTGGATGTGGCGCACCTACAGTTTGGCGCAACTGAAACTGATTTACAGACTTGGATCTCTGCTGGTCTGACCAAAGGCCAAGATGGCCTGTGGTCAGAC
GAGGAGGGGAGAGTAGGGATACCACAAATTGCCGCACCATTGTTTATTAGTCATTTTCATGGTTTTGGCCACATAAGTAAACAACAGACCAAGAATCTGATGACATCCAAGTATGT
CATTAAAGATCTGTTGCGCATGATTGATAGACAACTGGACAGATGTCTAACTTGTGCCAGAAATAATCCTGGAGGCCTTGTACATGGTAAACTTGAGCATTTACCACCCCCTGATGG
TCCTATGCAGGTATTGCAAATTGATTTCACACACATGCCCACTGCGCGAGGTGGATACAAGTATCTTTTAGTCATTGTTGATCAATTCAGTAAGTGGGTTGAGGCCTTTCCAACCAC
TCAAGAGAATGCACGTACTGTAGCAAAGATATTGTGCAAGGAAATTATTCCTCGTTTTGGTTGTCCATTACAGATAAACAGTGACAGAGGTACTCCTTTCACTTCACAGATTACACAA
ATGATTTGTACTATGTTGTCCATTGACTGGAAGTTTCATATTCCTTATCACCCACAGTCATCAGGACAGGTAGAGCGCATGAATAGAACCATTAAAGATAAGTTCAGGAAGGGAACTG
GTGGTACATTTACTAATTGGCTTGAACATTTACCTGCAGTATTGGCAGAAATCAGAATGACCCCTAGCACCACTACGGGTTATTCACCTTTTGAAATTCTGATGGGCAGACCTTTTC
CAACCCCATGGTGTAAACAAAAAGGTTCTCCTGTGGTAAGGGGAGATATCAATGTTGTAAGGGAAGAATATGTAACTAATCTGATTGAAACATTGAATGGCATCTATGGTGATGTTTC
TTCTACTCTTCCTCAGCCTACAGGTATCCCGACACACCCGTTCAAGCCTGGAGATACTGTCTTGGTGAAGCAGCTCCACAAGAATAAGTCGCTGGATCCTCCTTTTGGCCTTCCA
ACTACAGTGATCGCTGTGACGAGAACCGCTGTCCTTACAGAGGAAGCTCCAGTATGGATCCACGCGAACCGGATCAAGAGGGCCCCAGACGACGTCAACCCTGGCCGCTC
CATGATTCCATCGAGTATCGAATAGTGTGGTACCTCCTGCCAGCAGTGTTCTTCGCATCCGTGATTGTCATCCTGATCCTAATCTTCAATCCTGGTGCGCCCCTGCACAAACTCAG
GCCATTCCCTAGTCCTGGTGCGCCCCGGCACAATGATAATAAAAGTGTAATTGACAATGTCACTAATCCCTTATCCATCCCTCTCACCCGTGATGTCAGAGAGTTGCATATGTCGG
CATGGGAAACTGACAATGCTTTTGTTAAAGCTGCCACGAACTTATACAAAGTGACTAATCAGACTGATTCTTGTTGGGTTTGTGGCTATGTCCCTCATGGGGGGAAATGGGGTTATC
CATTTGTGGGTTTTCCTATTACCGCACACAATTTGACACAAATGATAAATGATGATACTATGTGGAATTTTACACCAATAGTTAACACTGATAAAGGTTTTGAAAAGGATAGGGTACACT
TAGTCAATTGTGTAAACCATGGTGTTATTATGTCCCATAATCTTAATGGTACTAGTTTTCACCCCAGTACACTTAGTCAAATGCCTATTACAGTTGCCTTAATCTACAATGCAGCCACAC
CAGTTAATGCTACACTGTTAAATATATGGGATGGATCTTATCTTTCTTTTGAGTATATTCGTAATTTTACTGCTAAGTATACTGAACCACACAACACAATTCGTAATTGGTTCTGGCGTAA
CAACAGAAATATAGGTAAACAGACAGGTGCTTCTCCAGAAGCTAAGAGGAAAAGATTTCCCTCATGTCCTTCAGTTATTAAGTTACCTTATGGTTATTATTGGCTTTGTGAAAGATGG
GCAGTTAAAATTATACCATGTAGCCATCTCTCACCTTGTTATTTGGGTTACATATTCCCAGCATTAAGTATTCATGACACACTCCCTAATGCTAAACTGCGAACACGTAGAGCTTCTGA
TTTTACAGTTAAGACTAGTACTGCCGAGATAGCTACTGCATCTTGGTTTCCTACTGCAGCTGTCATGCGATTATATTATAAGCTTAATTTGTTTGCTACACTGGTAGATGCAGCTTTTAAT
TCCACTGCAGACGCCATAGAGTCCTTAACTATGCGACAAGAGCAGATCGCTCAGACTGCTCTCCAGAATCGTATGGCTTTAGATTATTTGCTAGCAGCAGAAGGAGGAGTATGTA
AGCGCATAGGTTCTTCTTGCTGTGTTTTTATCTGGAACAACACTGGTAGAATACACCATGATCTTGACACTTTGCATGACATTCAATCACACATACGTAAGCAGCCAACAGGTCTCTT
TGATGGTATAGATTGGACCTTTGGTCTGAGTTCCTGGCTAGGAGCTCTTGGTATGAAAATTTTAATGGGTTTTGCATTTCTTCTTTTTCTGTTTCTTGCTTTCTTTTTCATATATAAGCTTAC
TGTGTGTTTGATTAATAAGCTTACCAGTACTCATGCTAACCCTCACCATTCCTTTTTCCATAATGTCACCGCTCAGCCATCTCGCTATAAGGTTCTTTATATTCGTTAAGTTGTTTTAGTT
TTTATGTTATTCTGTTAGAATAAAAGGGTGGTATGTTAGGGGTAATATGAACCTGATGGCAGCCATCTTGTTCCTCGCAGCCATTTTAGAATGAATAGACTTTATTATACTGGGTTATTAT
GCAGTGAGAATATTATGCATGTTATGAGTTTCTTTAGCTATTCGGCCTTGCCAATGTACCCTGGCCTAACGCCTAGAAGTAAGATAGCATAAGATAATGTAAACAAGAGGCTTTAGAC
ACTTGGTTGTCTCTGCAGATGGTAGTTTGTTATGACTCTGTAAATATTGTTATGAGAAACTCATGTTCTAGCTTTTCCATGTATTTCTGCTCTGTATAACTGTGTAAAATGACGCGGTCCA
AACGTGTCATCCCCTTAACCCTATGTGCTGTCTCTGTACGCTATAAGACATGATTACAATGTTCAATAAACATGAATAGCTTTGGATCATAATGCTGCGTGGTCTGAATCTGTTCAGGG
AGCTCCTTATCAGATAAATCTGCATCTGCCTCTCGTGGTACCGTAGGCCCCAGGCTTTGGTCTGCGCTGACACCCCCCCGTTGGATTATACACCTTACA

AnERV8

TGTAAGGAAAGTGGTTGCAGTAATTGCTCATATGACACTTGCTTAAACTTGTGTTGCCTTTAAGAGAGTGAAAGCATAAGACACTTGCACTAACTTGTGTGACTTTGCCTTTAAGAGA
GTGTGCCTTTAACAGAGTAAAAGACATTAATTAACTTCAAGTAGATGACACTGAATTTTGCTAAGTCATGCTACAAGTGGTAAACAACCGATGGATGCTGATGATAAGAAGTCAAAAC
CCAGACAGAACAAGGGTGTAACCTTGGCAGGGGTCACTTACTGACATTAAGGAACACATTGGTATAGGGGCTAATGCCATGGTAATATTGTTAATTATTATTAAGTGCAATAGGCTA
AGTAATATTAATTGCAATAGGCTAAGTAATAAAGCCATTATGATGTTTAGCTATTAATTATATCTATGGAAAGATTGTTAAAACTATTATTGATGTATTTGACAAAAGTAACTGCATAAGACC
ACATAATTAATGTTAGGCAGAAAAACATGAAAGATGACGGCATTGCAGTTTAGGAATACGTCAGAGATAGTACCTGGGCATGCCCTGATAAGCTTGGTTGAGGCTAAGACAGACTG
GCCAGATGTCTTGGGCCACGGCCAACCTCAGAGGGAGCGGTCTCAGGATCTGTGGAGTTGCGCCAAAACTGTTACACAAGTGAACAGTATAAAGAGGGGTACATCTAGGCAA
TCTGATCAKTTTGATKTTGAGTGTCTGAGTGTCTGAGWGTCTGTGTATTTTTGTGTGTGTATTTGAGTGTATTTTTGGGACCTGTGGATGGACGCATCTTCAGTGTTGTTTCCCGCTGAT
CGCCTGTTTTTCTCCTGGTCCAAGATTGCCCAGATGTCCCCTCCCATTGCGCCCCAGAAGCGGCAGGATAGGATGATGCAACCAGAGATCACGGTGATGTATTATTTGTTTGTTT
AAAACTAGCTAGACTTGTTTTAGTGGTAATACAACATAAGATTGTTATCAGTCGTTGCGCAGGCAACTAAATGTAACTGAAAACTGTATAGATATTTATATAAGTATTGCAATTGCATAGA
GTGTTATAAGTATGAATAGTGGTTTGTGGTATTTTGTTTCCATGATTATGTTGCAGAATAAACCATCATTGTTATTATAAGATTGTGAAAACCTGTGTAATGTGTCAGTCATTTCCTTGCTGA
AAAGTGATATATGATAACATCATATCCTGAAAAAAGAATCACATAACATTATTGGCGTAGTCGGCAGGATACATAATTGTTTATCACTTGAGCAAAGAAACTGACAGTTTTCTTACTGAA
TGTGCAGAGCTGATAGACGCAGCTCCGAGCGAAAGTGAAACCCCTTTGTAGATAAAATAATAAGCCAGAAAAATATATAGTTGTATCTTGCACGGTCAGGAGTATAAATAATATGTTT
CTGGATTAGTTAGAATTGAATTGTTAATTCCTCAAATGAGCCATGAAAGCTGTGTAAATAAACAGATATTTCATGGACATAGTCACAATTGTCACCTTGGAATGAAATTAATTGATATTGC
ACAGGTATATACATATTAACGTATATTGTTAYTGTTACTGTGAATTTATTGATTGATGTCAGCAATAATTGAAGGTATTATGAAAATGTTTAGGAAGAAAGCACAAACGGTTCCCCCTTCA
CGGCCTCAAAATATTGCGGCTTGGGAGGGGACCCCATATCAGGAATTAGCAGGGGAAGTCTTGTCTGGGACTGACCTCCCAGATAACGCTGAATTTTTTGACAAAATAGAACCT
TGTGAAGTGTTGAATAAATTACAGAATGTAAAAGTTAAGACAAATTCAGACACAGAAAGGTTACATAAGAAATTATGGGTGTTATTACAGGCATACAGAAAGACACACGAGCAGTTAG
AGGAAGCTGTAGAGCAGACGGTGTGCCTGCAGAGTCAGAATAAATTGCTATTAAAAAAGAGCCAGTGTTATGAAGATGTAGCTCAACAGGCTGTGGTTAAGGTAGCCAGGGCCA
AAGTAAAGCGGAAGGGTGGTAGGTTGAGCAAARGTAAGTTACAATCAGTATTACAAAGCGGGGCTAATTGGGACCCTGAAAGGTGGGACGGTAATATTTGGGATAGTAGTGATCC
TGGAGAGGAGTGGGATGATAGGGACCCTCACAGGGTGACAGCTCACCCTGTTACACGGCGTAAGGTTTATCATGGGCCTGAGGGTGCAGAAAGAGTAGACACAGTAGAGGAT
TACACGCAGCAAGAGGTGTTAGACATAATACGGCAGTATCAACAGAAACCGGGAGAGGACCTCATAACTTGGCTCGTGCGGATGTACGATCAAGGAGGTCATAACGTGACATTA
GATAGTCAGGATGCCATGAAATTCATACAGATCACTAGCGATCATGCCATACAACAGGTCCTGCGAATTGGACCTGGCCTGGCCGCTGATGGTATGTTGACGCAGGCATCATTGT
TGGTTCAGTGTGCATACGGTGCCAGGGACAAATATCCCACAGAAGGGGATTGGCCCACTGAGGATAGGACATGGCATTCCATTAAGGAGGCCATCCAATTATTAAAGGAATTGG
CTATGAAGCATGCGATATATAATGGACACCTGTATGCCCCATATCGCGAGCTACTAACCAATGCCATGAGAAACTATTTGATACGTGGGGCTCCCCGAGCATACAAGTCAGTAATC
ATGACGTTACTGATAAATCAGACAGGGAATCCCATTTATGAGGTAATGGATGCAATAAGACAAATAGGGGAGCTAGGGGAATGGGATGCCCCAGTAAAGCCTGATAAAGGCAGC
GGGCCGCCCCCTCCTAGAGCCAATGCTCCTCCCCGACTGTCAAGGAAAGAGATGTTCACAGAACTCCTCAAAGCAGGGGTAAAGAAGGAACAAATAGATGGGATAGAAACTA
AGGAAATGTGGAAGTTATATCAACAGCATGTGGGCAAAAGGATAAATGTGCAAAAGGGAAAGTCCGGTGACCTCATAACAGGTACCGCAGTCGCCCCTTCTGCCCCCACGCTA
GATGATATACAAAGGSCTACCACAGGGGAAAGCAAAGAGACCAAGGAGATACTGCAGCTGATATGTGATCAGTTGGAAAGTATAAAAAAGGATAGTAAAGAAGCAGCAACTAAGA
CCCGGTTATATCCTAGATTGCCTTTGCCTCTGGGATATGGTTACCAGGGAGGGATGAGAGAGAATCAGGTCCCCATGGCCCCGGCTAACACCCCAGAGGCCCAAGGACCTGA
GTTGGATGTGCTATCTGTGGGTGATCTGGAAGAATTCTTCCAGAACCCATAGGAATCAGGTCAAACTCCCAGTATCAGGGCCGCCTATTTTGATCATCGCCCCCATGTGGAACTA
GAGATCCACTGGAGCAAAAATAATGTACAATGTGAGCGGGCTTTGATTGATACGGGAGCGGAGGCCACACTGATACAGGGAAATCCGCGGGCATTCAAGGGGGAAAAGACAA
TAATTACCGGGCTGGGAGGGAAAACTATAGAGGCAACAAAAGCATGGGTAGATTTAAAGATAGGGCATCTTCCTAAACGGCGATGTTCTGTTTTAATCGTACCCATCCCAGAGTA
CATAATTGGCATTGATGTGCTAAAGGGAGAAACATTGCGGTTAGAGGATGGAATGTACAGCTTTGGGGTAAGGCCTTATGTGTCCATCAAAACCGTTACTGTAGGAAAATTGTTGAT
GCCCCCCATACAGGTCCCAGTGGCCTTGAAGAGGGTATACATTCGACAATACCGAATACCGGGAGGGCATAAAGAGATAGGTGACACCATCAAAGATCTAATAGCTGCAGGG
GTGGTGAGACCTATCACCACATCTTGGAATAATCCCGTTTGGCCGGTCAAGAAAAGTGATGGGACCTGGCGTATGACTGTGGATTATAGGGAGTTAAATAGTAATACTCCCCCATT
AACGGCAGCAGTACCAGATATGGTTACGCTGATAGAAAATATACAGCGTCACCCTGGAAATATGTATGCGGTCATTGACCTGGCAAATGCCTTCTATACAATACCTATAGAGGAGC
AGTGCCAAGAGCAATTCGCATACACCTGGCAGGGACGGCAATTCACGTTCACGCGCCTCCCAATGGGCTATGTCCATAGCCCCACGATCTGCCACAGGATTGTGGCAGAAC
ACATGGAGGAAGTTACTCTGCCCCCCAACGTGCAGGTCACACATTATATAGATGATATAATGATACAGGGACCTGATGAGCAAGTAGTGCAACAAGTGTTAGAACAGGTGATAAC
CCACTTAAAGAAGAAGGGGTGGGAAATCAACCCTGCCAAAATACAAGGGCCAGGAACCTCAGTACGATTCCTGGGTATACAGTGGACAAATGGCCGTAGGGAAATAACCCCT
AAAGCCAAGCAGAAAATAGTGGAATTTACATCCCCTAAAACCAAAAAGGAAGCCCAGCGCTTCATTGGGTTATTTGGCTTTTGGCGGCAGCATATACCCCACCTGAGCCAAATA
CTAGCCCCCATCTACAAGGTTACCAGGAAAAGTATAGCTTTGAGTGGGGGTCTAAGGAGGAGTTGGCTTTTGAAACTGCAAAACAGGCCATCCAAATGGCGATGGATCTGTGG
CCAGTAAAGGAAGGGCCAATAGATTTAAATGTGTCTGTCCACAAGGAACATGCTAATTGGAGCCTTTGGCAGAAACAAGGGAGAGGGAAAGTCCCCCTTGGTTTCTGGACTAGG
AAATTGCCTGAGGCAGGTGACCGATACACGCCCTTTGAGCGCCAGTTATGTGCTTGCTACTGGGCTTTAGTAGAAACAGAGGAGCTAACGTTAGGGCATGATGTATTCTTGAGG
CCTGAGATTCCCATAATGAACTGGGTAATGGGCAGTCCTAAGACCCATAAGATTGGACATGCTCAGGAATCAAGCATCATAAAATGGAAATGGTATATACAGAACAGGGCAAAGA
CAGGTCCCAAAGGGGTTGCCGCACTGCATGAAAAGGTTGCTGAGCAACCCCTCCAGGGGACAGCCCCCATTCAGGAGACACYAACAGAGGAGTCTCTAGTAAAATGGGGTG
TGCCATATGATCAACTGTCAGAAGAACAGCAAGCCCATGCTTGGTTCACTGACGGATCAGCAAAATATGTGGGTTCACAGAGGAGTTGGAAAGCAGTAGCCTACAATCCTGTGA
CCAAACAGATGCTGGTGTCTACAGGGAAAAATGAAAGTAGCCAATACGCTGAGTTAGTTGCTGTCCATCAGGCCATAGCCTGTGAACAGGGCGAATGCCACGTATACACCGACT
CATGGTCAGTGGCAAATGGGCTAGCCACCTGGCTCCCCACCTGGGAACGAAAGACTACCAGATATACTCAAAGGAGGTGTGGGGGAAGCAGATATGGCAGGAACTCAGCA
GATTGGTTAGGGAAAAGACAGTGACTATTTACCATGTAGATGCCCACACAAATCAGGATTCACTAGAACGGCTGTTCAATTCTATTGCAGATGAGGCAGCCAAAGTCTCCACTGTG
GAAAGTGAGGATGTGGATGAGGAGGCTGACAATAAGGGTGTGGCCCAGTGGGCCCACCAAAAAAGTGGACACCTTGGAGAGAGAGCGACTCATAGGTGGGCATTGCAAAGG
GGTATCAAGCTATCTATGGAAGTAATAAAACAGAGATAGCAAATTGTCCTGTATGTCAGCATGTCAGGAAGCGACAGATTCCTGGCATGGTGCAAGGACATATCAAAAGAGGCAA
GCTGCCGGGCCAGATCTGGCAGATAGATTTCATAGGACCTCTTCCCTCCAGCAAAGGCTGCCAGTACGTGTGCACGGCTGTGGACACGTACTCAGGCTACCTCATAGCCCAC
CCATGTAAGCAAGCAACTCAGATAAACACCATTAAGACCCTAGAATTAATCACAAAATTCTATGGCACTCCATTGCAGATACAGAGTGATAATGGTACTCATTTTACAGGACACATG
GTACAGGGATATAGTGCAGAAAGGTACATACAATGGATATACCATATACCCTATTATCCCCAAGCAGCAGGATTAATTGAGCGAATGAATGGTTTGCTAAAGCAGCAGTTAAAGAAG
CTAGGGGATGGTACCTTGCAGAAGTGGAGGGACCATCTTGAGGAAGCAATTAATGTGTTAAATAACCGTCCTATTACAGATAGTGAGACCCCACTGTCTAGGATGATAAGCAGAC
AGGAGTACACTGAGTGTCGAGATCAACATGCTGTGGTAACGTGTTGGGAAACTAGCCCTGGGGGTTTAGCGCCAGATAAATTGCTCCCTCCTAGTGGCGGGATATACTTGACCA
TCCAGGACCAGGTAACAATATCCCCTGAGAGCTCCCAAACCATATCAACAGGGATAGGGGTCCAAATACCTGACAAACATGTGGGTTTCATAGTTCCAATGCCCTCTCTCATAGT
CAGAGGTATTAGCATAGTGGAGAAAACGCTAGACCCTGGAAGAACTGAGGAAGTGAAGCTCACCCTCCTCAATAGAGGAAAGGAACAGGGAGACATGTCAGCATTAGAAGTAG
TAGCAAAGCTTATAGTTACTCCCACCTGCAACAGTGAGATAGCGGAACAGGAAACTGCAGAAACTAGGGGAAGGGTGGGCATAGGGGCTAAGGTCTGGTTGAAACAAGAAAA
GGGCCCCCCCGTGCCTGCAGAGGTGATAGCTAAGGGAGATGACAATGTGGTAATTGTGGTAAAGCAAGGGAGCAATGATTGGCTGCATGTGCCTGCGGGCAAGTGCTATTTAC
GGAATTAATTTGAAGTATGTTTCCTTCCTAGATTCTTGTTGGCATGGCTGTTATTGTAACCGGTGCTCTGTTGGGACTCCTTTTGGCCGCATGCGTAAGCCAAGCACTGATGAGCCA
GAGGATGAAACGTGAGGTGTCCGGTGCCAGCTATCACACCTTCTATGAGGGAAGCATGACCGTTCCGTATTGCACAACATGGGCATCCTGCTCCATCCCATTTACCTTTGATGA
CGACGTAACCTGTTTGACCAAAAATGAATATGGAAGTTGGATGTTCCTTACCAAGCGGGAAAGGGTGGTATTTGAACTATCAGAGACAATTACCTGTGACATCCCTTCCTTCACTGT
ACATTGTGTAACTAAGATTTTTGTTCTGCTGCAGTGACCAGGGAAGACTGTAAGGAGTCTAAGGAGGATGAGAGCTGGACAATTGTTTTACCGGATGAGAACTTAATATTTGAATTG
CAAGAAGGGGTTGGGTTGATTTGTGAAAATCCCAGTATGCCTTCCACTTCCCCAAAACTAACAGTGGGGATTCAGGAACGCATTTTAGGTTGTAAAACATCCACTGATACACAGG
TTCGTGCTGTAATGTTAAACATATCATACACTGATGGGGAACCAGAGGGACTGACATATGTGTGTAAATATACAGGAAAAGAGTCATCTGAACCTCTTTTAATGTATATCAGGAGACCT
GTAATGGATGGGAAATTATTAAACAACCGACACAAGGGAAAGAGGGACCATGCACGGCGCCATTTAGGCTCCCTGCATGCCTCCTCATGGGAACTTGATAATACCTTTGTAAAA
ACTGCCAAAGATGTGTTTAATGTGACAAATGTTCAGGGACCATGTTGGGTATGTGGGTTTGTGCCTCACGGGCAGACTAGGGGTTACCCTTATTTAGGTGTCCCTTTAACCATGAGT
ATGCTTGCCGATATGATTGGGAATAAGACTGTGTGGAATTTTACAGGTCTTAACACACCAGGTAGGGATGACTTGCTGGATAAGTTGCATCTAGCTAATAATGTGACTACAGGTGCTA
TTATGTTTCAAAATCTAGATGGGGATATATATAGTACTAATAACCTAACCCAGCTACCTGACATGATAATGTTAATGTATAACATAGGACGGCTAGTTAAAAATAGGACAGGGGTATTCA
ACATTAAACTGAAATTCTATGACTTCTCATATTTGACATATGCCCATGAGAAATGGAAGACTAAGTTTGGTGAGCCCAGAGACCTTATTAAAGACTGGTACCAGCTTAATAGGGAAAC
ACTAGCAGGTAACACACGGACTAAGGATGGGAGGAAGCGATTGAATAGGATATGTCCTCCTTTTGTTAAACTGCCAGTGGGATATTATTGGTTGTGTGGGAGGTGGGCAGTAAAG
GTAATACCTTGCACCCATTATGGCCCATGCTTTTTAGGATATATATTCCCTGCTTTTAACATAACCACCACGCTTCCACCACCTATATTGCATAGAACACGAAGATACCTTGACCTGG
ACAAAATACACAATGGAGTTAAAATTAGTGAAGGTATGCGCCTGCTTGCAACATTTGTCCCTCATTATGGGGCTGCTACAGCCCTCACTAGATTAAAYACGCTAGCTGATTTAGTAG
ATGAAGCTTTTAATGTAACTGGAGATGCAATAAAGCTTTTAGCATTAGAACAGGAACAAATTAGGATGGTGGCATTTRCAGAATAGAATGGCTCTAGACTAYCTCCTAGCTGCAGAAG
GAGGAGTTTGCCARAGAATCCACCAGCAATGCTGTGTGTACATAGAAGACAATACAGGTAAAATCAAACATGACCTACACAGACTAGAAGAAGTGCAAAAGCACATACGTGAGG
TACATGATGATTCATTTAGCAAATGGCTTAAAGACTTTGACTGGACTTTTGGACTGGGGGGATGGCTTGGAAGCTTAGGCAAAACTATAGTTAAATGGCTATTAATAAGTCTGGCTTG
CTTGTGTGGGATATTCATTATAGTAAAAATGTTTGGATGTTTCTGTAATACATTAGGTAGGATGTGTTTTAAACCAAAAGACAATATTACTTGAGTGTTTTAATTATTAAGGTATGTGTTTAGA
AACAAGCAAGGTGTCACATGACCAAGGGGTGGAATGTAAGGAAAGTGGTTGCAGTAATTGCTCATATGACACTTGCTTAAACTTGTGTTGCCTTTAAGAGAGTGAAAGCATAAGAC
ACTTGCACTAACTTGTGTGACTTTGCCTTTAAGAGAGTGTGCCTTTAACAGAGTAAAAGACATTAATTAACTTCAAGTAGATGACACTGAATTTTGCTAAGTCATGCTACAAGTGGTAA
ACAACCGATGGATGCTGATGATAAGAAGTCAAAACCCAGACAGAACAAGGGTGTAACCTTGGCAGGGGTCACTTACTGACATTAAGGAACACATTGGTATAGGGGCTAATGCCA
TGGTAATATTGTTAATTATTATTAAGTGCAATAGGCTAAGTAATATTAATTGCAATAGGCTAAGTAATAAAGCCATTATGATGTTTAGCTATTAATTATATCTATGGAAAGATTGTTAAAACTAT
TATTGATGTATTTGACAAAAGTAACTGCATAAGACCACATAATTAATGTTAGGCAGAAAAACATGAAAGATGACGGCATTGCAGTTTAGGAATACGTCAGAGATAGTACCTGGGCATG
CCCTGATAAGCTTGGTTGAGGCTAAGACAGACTGGCCAGATGTCTTGGGCCACGGCCAACCTCAGAGGGAGCGGTCTCAGGATCTGTGGAGTTGCGCCAAAACTGTTACACA
AGTGAACAGTATAAAGAGGGGTACATCTAGGCAATCTGATCA

AnERV9

TGTATTACAAATAAAAATCTGCCAGTTTTTATATAACACTAATGGTTTATTGAAAGTAAAATTGTATCATAGGAATCTATCTGTCTTTTAATTGCTGAYTTGGATGTCACTGTGCT
GACTCAGACATTGGGAAATATAGCTSTCTCTCTCATGAAGCCATTGTTTATCCCTTAGACAATGTGTACGTTTTCCTGTCACCCTCTACATCCTCGTTTATGTCTTCCCGTG
AACTTTTATTGGGGAAATGAATGTTTATGCAYATTTTTCCCTCCACTGTCCATATCCTGAGTATTGATTGTTGACTTCACACAATMCATTGTTAGAAGGCTGACATCTTAATTT
AAGACTTACGAAACTAGGGCGGGAATTTGGTCTTCGGAAGTCTGAGYTTGCAAGCTAGGTTGACCCTAAGGTCAAGTAAGGAAATGGCCATTCCCTGCTTGATTTTGG
GTAATTTATGTCTAGTCAGCAAATAACTGCTGACTTGYCTCCATAAAWAGGGGTTTCGGAKTGCGGTGTAAGACAGGAATTATTAACCTGTGCTTTGAGACGTCTTAGCA
CAGTGTGGCTGCYACGAATAATATCCTGTCTTTTCGCCGTTATTCTGAAGATTCCTCCGGCTGATGCTCAAAGACAGATGATGACAGAAGCCCCCTCTGTTTCGTGGTG
AAGTGGTGTCAATTTCCCATTTGAGTCTCTCTCTTGAAAGATGATGATACAAAGCTCATGGTGATGTAAGTTTTATGTTTATGTATCTATATTGCTGTACCTTTGTTTGCTTATTT
AGGAGTGTATTTGCYGTGTTCTTTGTATTCTATATATATATATATATACATATTATTTGTAATAAAATCATAACTTTGTTGCAATTTGATCTCTTTAAAACAATATAAAGAACCTGCA
TTTATATTAGTAGCCCTGCCCTGGGCATTAATTAGTAAAGTATTAGATATTTGGGTGTGCTCTTGGCCACCAGCAAAACATTTGGCGTAGTCGGCAGGATATTGTGGAGA
TCAAAGTATATGTATATATAGAATATATATAGAGGAATATACTCATAATGTCTTTATTTAAGCGACAAAGTGTTGTGAAGGGACCAGTGGTAATACCAGGRTGGGAGCAGGC
TCCTTATCAGGTGTTAGCACATGAATGGTCTCAATGTGTTGGTTTAGGTGAGTCTTGGGACTCTTGTTTAAAGAACACAGAGCCCTCTGATGTAGTAGAATGTTTACAAAG
AGTTAAGATTGAAAAAGGTAGAGAATTTGCTTGTGTTAGYAGGAGGGGGTGGTTGCTGTTAACAGCATATCGCAGGGAAAATGAGGAAAAGCACAAGCTACAGGAAGA
GGTAACTAGATTAGAAAAGAACTAAGTAAAATTAATGGCACTTTGATAACWACTCAGTGTCAAAATAAGTTGTTRATGGACAAACTTGATAAATATTCCACMATTGCAGA
AAAAGCAGCTGTGAGAGTTGCACAGTACAAATACAAAAAGAGGCGAGGTAAAGTAGATAGCCGAAAGGTTAGAACTTTGATAGCCAARGCTGATGCTGATTGGGATYT
TGATCACTGGGATGGGGATATTTGGAATTCCACTAGTGAAGATGGGGACACTTCTGAGGAGGGAGGWATAGAAATGCGCCCTGCTATGAGGAGGAAAGAGGAGAAA
ACTATTGAACCAAGGGGACCTGATAATCAGGGTAATCCTCGCCCCCCCAGAGAAAAAATAATAGGTAAGGAAGTAATTGAAGATTTTACGCAGCAAGAAATTATGGATT
TGTTATCACGTTTTAAACAAAATCAAGGAGAAAGTTTAATTAAGTGGATGGTTAGGATTTATGATACTGGAGCTGCAGGTATCATGTTAGATGAAAAAGATGCCCCTAAATT
TTTGACAGTGAGTACTGATCCCCATATACAGACAGCTTTTAGAGAWTACCAAGGGGGAAATTCTACTTTATTGGCAATTGCTGGTGCTGGGTGTGACAGGAAGTATATTT
CAGAAACAGAATGGCCCCCTAATGATAAACCTTGGTTCACCCTGAGGGATTGTATACAGCGTATTAAAGAGGAAAGCATGAAAATATCTATATACACTGGACATAGTGAT
SAGATGTTATTAGGCCCCTTTGCGTTAGCAACACGCAACAAAATCATAAGGACTGCCCCACCTGCTTATAAATCTGTAATAATGACYTTGTTGGTGAATCATGTAGGTAAT
CCAGTAGGGGAAGTAATTGARGCAGTAAGACAGTTAGGAGACCTGGGTGAATGGGGTCCTAGTCAGGGGGATCGTGGAAGGGATATGACACGGAATGTTCCCCGA
GAACGTCAACATTCGTCTTCAGTACAGGGATCCCGGGTAAGTAGAAGAGATATGTTTATGGCTTTGTTAAAGGATGGTGTTGCCCGTGAGGAGATTGATGGAATAGGGA
CGGATCAGTTGTGGAAATTGTACAAAAAAAGAGGTCTGAATAACCCAAAGAAAAGTACATCACAGAATAAACCTATTCATGTAGAAAAAGCAGTGCGAGGTGATAAGG
GATCACACTCTGAAGGAGGGCCCACCTTTGGGGAGTTRTCTCATACYCGCCCCACTGCTCCGGGTTATGAAGATCTSTATGCAGATGCAAAAGCTTTGTTGTATGGAT
CCAAACAAAAAAGGCCGTCCCATTCCACTAATCCTTTTTATGGGGAAGCCAATACATCAGATTGACTAGCWATAAATAGTGTACAAAAAGATATTCGCCCCCATGTAG
ATGTAGTTATTAACTGGAAACACAGAAATTCACAACAAGTCAGGGCACTGGTAGATACAGGGGCTGAAGTAAGTTTAATTTATGGAAATCCCCGTAAGTTTAGTGGAAAA
AGAGTAGTTATYACTGGACTAGGGGGAAAACAAATAGAGGCAGTACARACTAAAATTAATATGAAGATAGGAAAYTTACCAACTCAAAATTACTCAGTAATGATAGTCCCA
ATACCAGAATTTATAATTGGAATTGATTTATTAAAAGGACTAACTCTTCAATTACCGGATGGTAAGTATGAATTTGGGACCCCCAGAGTAGAATCTTTTAATGTTCATAATCT
GAATGTGAATACAGCCACAGTACAGGTAGACCCTGTACTAGTAGGACATGCCAAAATACCTCCAGTATCTATACCACCTGCTACTAAARTTGTCAGTCTGAAACAGTATA
GGATTCCTGGGGGTGACAAAGAAATAGAAGAAACTATACAAGAACTTTTACAGGCAGGAGTTATAGTGCCAACTACTACACAGTGGAACAGTCCTGTGTGGCCAGTW
AAAAAATCAGATGGCACATGGAGAATGACTGTAGATTATAGAGAATTGAACAAGAATACTCCTCCCTTGACAGCTGCAGTGCCTGATACAATTACTTTAATTGAACAGATT
CAAAAACATAGAGGCAATTGGTATGGGGTCATTGATTTAGCTAATGCTTTTTTTACAATACCCATAGCTCCCGAATTTTGGGATCAATATGCTTTYACGTGGAAAGGAAGA
CAATTCACATTCACCCGTTTACCTCAGGGGTGGGTACACGCACCTACCATCTGYCATAGGGTAGTGGCTGAATGTCTAGACACCTTTGACTTGCCCAAAAACATGCA
AATATCACATTACATTGATGACATTATGATACAGGGAGAATCACAGGCAGAGGTAGCAAAATATCTACCAAAGTTGATTGAACATATGAGGAGTAAAGGGTGGGAAATCA
ATCCTGCCAAAGTCCAGGGACCTTCTCAAGCAGTAAAATTCTTAGGTATTCAGTGGAATAGGGGACATCGAGAAATAACTGATAAGGCCAGACAAAAGATCTTAGAGT
TTGCTGTCCCTAAGACAAAGGTACAAGCTCAGCAATTTATAGGACTATTTGGGTTCTGGAGACAGCACATTCCCCATTTAGGACAGATTCTAGCTCCTCTATACAGGGT
GACAAGGAAGAAATTTCATTTTGAATGGCTCTCAGAACAACAACAGGCATTTGAAAAGGCCAAAAGTGTAGTACAACAAGCACTTGATTTATGGCCTGTTCTGTCCAAT
GTCCCAATTGAAATTAATGTGTCAGTGCAGGAACAGTATGCTAATTGGAGTTTATGGCAGAAACAAGGGGGGAAAAAGGTGCCACTAGGTTTCTGGAGCCGTAARTTA
CCTGATGCTGCGGCAAAGGTATACTCCTTTTGAAAGACAGCTACTGGCYTGTTATTGGGCTTTAATAGATTCTGAACAATTAACTTTAGGACATGAGGTGACTATTAGGCCT
GAAATTCCCATTATGCACTGGGTACAGAGTAATCCAAAATCACATCGCATAGGTCATGCCCAAGAGGCCTCAATCATAAAATGGAAGTGGTATATTCAGGACAGGGCA
AAGCCTGGTATAGCAGGTATAAGTTCCTTACATGAAAAAGTTGCCAATGTCCCTCATGAGAATCAGACTGAACATGTCCCTGAAGAAAGAATAATTCAGGATTCCCCTG
TAAAGTGGGGAGTTCCTTATGAACAGTTAACAGACAAACAAAAGAAACATGCTTGGTTCACTGATGGTTCAGCTAAATATCAGGCTGGAGAAAGAGTTTGGAAGGCGG
CTGCCTTCCAGCCAAGCACTGGAAGACTTATGACAACTACAGGTATAGGRAAAAGTAGTCAATATGCTGAGTTATACGCTGTATTTAGTGTACTGAGACAGGAAAAATTA
GAAGAGTGCCACATTTACACTGACTCCTGGTCGGTTGCAAATGGYTTAGCTACCTGGCTTCCAACTTGGGTAAGTAACAGCTGGAAAATCTATAGCAAAGAGATATGG
GGAAAGGAGTTGTGGCAAGATATCTGGGAAATTGTATCACATATACCTGTAACYGTTTTTCACGTTGATGCTCATGTCACCACAGATTCCCTGGAGCGATTGTACAATTC
AATGGCTGACGAGCAAGCCAAAATTGCACTCACCAATATCGTGCCTGAGGTGGAATCTTCATCTTTACAGGGKTTTACTAAATGGGCTCATCAAAAATGYGGACATCTC
GGAGAAAAAGCTACCTATCAATGGGCCATATCTCGAGGAGTGCCCATCAGGTTAGACATGGTGAAAGATGTTATAGCTCATTGTCCTGTCTGCCAACATAGTCAGCAG
CGACCAATTCCCCATGTGGTGAGAGGACAATTAGCTYGTGGTAAACTCCCTGGTCAGATTTGGCAAATGGATTTTGTTGGGCCCCTACCAATGAGTGGTGGGTGTCAA
TATTTATGTACAGCTGTAGACACATACTCTGGGTACTTGATTGCCCATCCGTGTAAGTATGCTAATGCTGCAGCCACCATACGAACATTAGAATTGATTGTCCTGTATTATG
GAATTCCTCTACAAGTTCAAACYGACAATGGCTCTCATTTTAAGAACAAAACAGTGTCACAATTTTGTGAGGATCACAATATTGAGTGGATATTCCACATTCCTTACTACC
CCCAAGCAGCAGGACTGATAGAGAGAATGAATGGTCTGTTAAAAGAACAATTACATAAACTTACTAATGAACAAATGGGAGTTTGGCGTAAATATTTAATTGATGCCTTG
CAAATTCTGAATAACCGCCCTATCGGGAATCAAGAAACTCCTTTAATGAGGATGATTACGCCCAATTTAACTGTAATGCCCCTTAGAGCTGCAATGTCAGGAGAAACTG
TGGTTTATTGGCCTTTGAATGATCARGTAAAAGCACCATTTCGAGCAACCCCTGGCTCAGCTGGTTTAGATGTCCATGCCACAGAGGAAATTACCCTAGACCCTGGGA
AAATAGCTACTTTTAATACAGGGCTRGGAATGCAGTTACCTGCCCATCATTTTGGATGGTTAGCTCCTCGCTCAAGTCTAGCCTTAAGAGGTATTCAAGTCTTAGGGGG
AATAATAGATGAAGATTATCGAGGTGAACTTAAAGTTCCCTTACTGAATAATGGAACTGAATCAATCGATTTTCGTATTGGAGACAAAATAGCTCAACTTCTGGTCATTCAA
CAGAAATATTTTGAGTTGGTAAGGGGGAATAGGCCTCTTATAACAACTGCTCGAGAGACAAAACAGTTTGGGTCAGTTGATGACAAGTTTGTTAATGGGACCAAAATTTG
GGTTAAATCACCAGGCCTAAATTCAGGCCCACCTGATGCAGCAGAAGTAATTGCACAAGGTCATGATAATACAGTAATAATAGTTAAATCTGGAGAAAATAAGTGGTAC
ACTGTACCCTCACACTGGTGTTACTTACGAGAATAATTTTATTTTATTTTTAGGTTGTGTGGCAACTTGCAATAATTTCCATTTTGATCATCCATTTGGAATGTGTATCTATAAA
TCAATGGCAATCAACTCGCAGTGTGACAACCAATCAAAACTTTTGGGAAACCAAGGAAAATATATGGTCATATCTAGCCCGAGTCATGAAYATCAGCCACTTTTGTTTAT
CCAAGAGTTCATCGATCTCTAATACCTTGGAAACGTGCCTTGTTTCTATCCCCACACCTATACAAATATGGACTTCATTGTTGAGAATTGAAYTAAAAGATGCTGATCTAC
AAGAAGGAGATATAACTCAGCACGAATTATATAATCCTTATTACACATGTTTCAATTGCACAGACACTACAATACCTACAGATGATATGGTTATGATACATACATATAATATTG
CTGCTGCAGATATTTGTTATATGATGACCTGTAATGCTTTGCAGATCCCTAGGTGTAAAATGAAAATAATGCAAGGGCAARGTCTTAGCAATGATACACTTTGCAAAAGAC
AGGAAAAAGACTGTAAATACATAAATGTTACTAACATGAACTGCAACTCCACTTTGATTGCTCCCTCTCACACCAGATATGTAAAATTACCCCCTGGGTGGATGTTTTCAT
GTGGAAATGACACATATAATTATATTCCAGCTAATATTATGGGGGGGCCTTGTGTTATTAGTAGACTRGGATTAATGATGTATCCATTATACCCCTCCGARTCACACAAATTT
AGGAGAGATACTTCTCATCTATTACCTGTAGATTGTAACTATGATRTTACTTTATTAACTGAAGGGGAAATCATAGCATTAGGTGTTAGTTTAGTAGGTGTGCCAGGATTGTC
CATATATACAGCAAGACAGTTAAATAGATTAACTTGTTTGGTAGCAAAAACAATAAATAGTACAAGTATAGCCCTAGAGCGTATGTTGCTTGATCTACAAGGGTATAAAAAG
GCAATAATGCAGAATAGAGCTACTATTGATTATTTGCTATTAGTACACAATCATGGCTGTAAGGAGTTTGAAGGAATGTGTTGCTTTAATCTTAGTGACCATTCCAAAGATAT
TCAAAAACAGATAGATGATTTAAATGGTTATGTAAAACATATAAAGCAAGATAATCAAGGGTGGTGGGATTACTTTTTCTCATGGTTACCTGATTTAGGATATCTGAGACAAA
TAATAGGAATTATTCTTGGTGTAGTAATTGGACTTTTAGTTTTAGGATGTTGTGTTCAATGTGTACCTTCTTTTTGTGTTATGTTTCGAGTTTCACCCGAACCAATCCCTACAG
GATACAATGTCCATACTTTCAGATAAACCAAGGGCCGAATGTATTACAAATAAAAATCTGCCAGTTTTTATATAACACTAATGGTTTATTGAAAGTAAAATTGTATCATAGGA
ATCTATCTGTCTTTTAATTGCTGATTTGGATGTCACTGTGCTGACTCAGACATTGGGAAATATAGCTGTCTCTCTCATGAAGCCATTGTTTATCCCTTAGACAATGTGTACGT
TTTCCTGTCACCCTCTACATCCTCGTTTATGTCTTCCCGTGAACTTTTATTGGGGAAATGAATGTTTATGCACATTTTTCCCTCCACTGTCCATATCCTGAGTATTGATTGTT
GACTTCACACAATACATTGTTAGAAGGCTGACATCTTAATTTAAGACTTACGAAACTAGGGCGGGAATTTGGTCTTCGGAAGTCTGAGCTTGCAAGCTAGGTTGACCCT
AAGGTCAAGTAAGGAAATGGCCATTCCCTGCTTGATTTTGGGTAATTTATGTCTAGTCAGCAAATAACTGCTGACTTGTCTCCATAAATAGGGGTTCGGATTGCGGTGTA
AGACAGGAATTATTAACCTGTGCTTTGAGACGTCTTAGCACAGTGTGGCTGCTACGAATAATATCCTGTCTTTTCGCCGTTATTCTGAAGATTCCTCCGGCTGATGCTCA
AGACAGATGATGACAGAAGCCCCCTCTGTTTCGTGGTGAAGTGGTGTCAATTTCCCATTTGAGTCTCTCTCTTGAAAGATGATGATACAAAGCTCATGGTGATGTAAGTT
TTATGTTTATGTATCTATATTGCTGTACCTTTGTTTGCTTATTTAGGAGTGTATTTGCCGTGTTCTTTGTATTCTATATATATATACATATTATTTGTAATAAAATCATAACTTGCAATT
TGATCTCTTTAAAACAATATAAAGAACCTGCATTTATATTAGTAGCCCTGCCCTGGGCATTAATTAGTAAAGTATTAGATATTTGGGTGTGCTCTTGGCCACCAGCAAAACA

AnERV10

TGTCAGAGAAAAAAGGAACCATCTTGTTTTCTGTACTCCATCTTGTTCATATCAATTAATTTAGGAGTGTTTTAAATATGTTTAAAAGTGTTGTTCTGTTAAAACTGTGAATTGC
TGATTTTGCATAATGGAGGGGCCTCCAACTAAAATAGCCCACTCCACACTTTGTACATGCGTCAGAGAGGAGCCTTAGTTCTATTGGCTATAGGGCTCCTCCACCTTT
CGTCTGTGTTCAAGAGATAACAAAGTATTGTGTAACTTTAACYGTATAAATGATGGTAGTTAAATGCTTAAACTTTGAGTAGCATGTTGTAGGGACAACGTCAATGCGTCTG
TCTGTCTACTCCTTTCACGTGAAATAAACCACTTCAATCAAACTGACCTATCCAACAGTTTTTCTTTGACAGCTTTGGTGCCGAAAAAAGCCCGGGAGTGAGGAAATTG
CAGAACAGCCATACCGGAGAGTGAAAAGGATACTGAGACCTTTGGCGCCAGAATAAGGTGAGACCACATTTTTGTCTCTGATATAAAGCTCTCACTATCAATTTCATGT
CTCTTTTGGCTGTATCAGCAAGAGTAGCACACTTCTAGAACCTTGGATAGGTCAATTGTAAGTTGTTTGTATGTCTTTTGTTCTCTCACCCCACTTTGTGTTTATGTTTTTTGT
TAACCYGTGCTAARCTGCTGTGCATACCCATCCCCATCTTTCCTTCCCATAGCGATTTTAGTGTGAAAGCCGTTCACGACCCATTACACAGTTTGGCATTGCCCATTTG
TGGACTGGTAACTGCGGATTCACTAATATCACTATAATCCCTTGTCAGCTTTTTGCTCTCTGTGTATGTTAACCCTTGTATGTCCTTCCTTAAACTCTGTGTATGTTAACCCT
TGTATGTCYTYCCTTAAACTCTGTGTATGTTAAYCCTTGTATGTCCTTCCTTAAACTCTGTGTATGTTAATCCTTGTATGTCCTTCTTAAATTCTGATAACCTTTGTATGTTTTTCC
TAACTTTAATCCTTATCTCCTTGATCACTAAACCCACTGTAATTGTGCATGTCCCTTAACAAAGTTTTTCCTCAAATACTCTATGACTAACCTTTCACTGTTAAATGATAGTGA
TAAATATTGTCTGACATTGGATGTAAACTGTAGCCAGTGGCTTTATATACTGAAATTGGTGTTGGCACTTCTTATGGGATTTGGTTGCATCTATCTGTTCTGGAAAATTTGGA
GGGATCAAAGAAGGGGTTATCCTGTCTGTAAATTTTAGCCACTTAAAACCCACTTTTGTGAAAATCTGTATATTGGGAATATATAGATTTTTGTTTCGTGTAAAAGGGGGGA
GCGTGATCATCTCCAGCCCAGTTCCATATGGGGAATACTTTACCGAAACCGAGGGGTGGCCAGACCCCGACTGGTAATTTTTCTGTTCCGGGAACAGAGACAATAG
GAGCTCCTGAGTCATCACTGTCACCGGTAGCAGTGATGAAAGAGAAATATGGCAAGTGGGTATGTAGGTGGCTGAATAAATGGTTCAAATTAACCAAACAGACTGATTT
GGCAATTCCACTGAAAGGTACCTTTGATACCTTAAAGTGGGTCAATTTATATAAAAATTTTAAATCAAAATTGTCAGATCCAAAAGAATTGAAAGCCTGGGGCAGATGGAT
GCTAGAATCTCGTCAGAAGTGCAAACGATCAATAGAGGTTCATGTTACTAAGGTGGGACAGAGTCAGGAAGAGGAAGAAATAGAGGGGGAGGATTTAATGAGGGAAA
TAGAAGGGGTAGCTTTGTCAGTGATAGGGGATTGGTTGCCTAGAAATACATTAAYGGGTAAGATTAGACCAACTTCTACTCTTACACCTACCCCCCTAACACATGACAC
TGAAACTACACATCGGTCACACTCCACCCCCAAATTAGCACCTACATGGACACATACACCACCAMCTTATAACCCTTTAACTCCTAGTTCCCCAGACATGCCCAGTT
TATCCTTTGAAGGGGGGTTTGATCAGGTGTCCCCTGTAACTTTTCCTCAGGTTCAGCCCCCCCATGGGACAGGCTCCAATATACAGTGGATAGAGGGCTCCCTAGAT
GCACAAATTAAGGGGAAAATTAATGTGCAGCCTATTAATACCCCACCAATAAGTCATGCTGGGGGTCAATTAATACCTACCTTCCCAGTTAGGGAACAGCGGATAATG
GGAGCCCAGGGACCCACAGGGATACATACCCATGTCCCTTGGACCCCCGGGGAGATGAAGGGATTTTTAGCCACCATTCCAAAACCCCGTCACAACCCTCATGC
TTTTGCACAAGAGATTTTAAATATAGCCTTATGCTACTCTCCTACTTGGAGAGACTTGTTACAAATAATAAAGGGGGTGGCCGGAGTTTTGGATATGAAGGAAATATGTACT
AAGTTGGACTTACCAGAAAATCCAACTGCACCCTTTGATACCCCCGATAAAGCCAAAGAATATGCCCAGAAACTGTCTACTAACATCTCCACAGTTTTCCCACCCCAA
TCATGGAGTCGGGTTACTAACATAGTACAAAAACCCAAGGAACTAGTGGAGGATTACTTTGAGAGGTTTAGRCAGGCAGTAGAGGCAGCAGGGGGGGAATTAGAAG
ATAAAGGGTTAGGAAAGGTTTTGACCGCTTCTCTTGTCTCTGGCCTTAACAAAAACCTTCACGATAAACTTATCTCTGCCCATGCGGACTGGAGAGATAAGTCAGTGCC
GGCTCTAATGAGCATTGCAAGTTCCATCCAGCGGAACATTGAGGAACTGGAGGAAAAAAAGCAAAAAGTGATGGTAATATATACAGAGACRGGGGGGGCAGTTAAAA
GATTTCCTGACCTATCAAAGGTGAGATGTTATAATTGCAAGAATCTGGGGCACTATGCTAGGACCTGTAAGGCCCCTAAGAGGGAGGAGTCAAGGAATAAGAAACAA
GATTGACAGGTAGGGCTTCCAGTGTTGCCAGTAATTTCAGAAACTAGAGAGGGGGATGCCTTTGTTCAGCTGACATTAAGTGAAATGCCTTTCTCTACACTTGAATGTCT
GGTGGACACTGGAGCTGCCCGATCAGTTTTAAAGAAATCTGAAGCCCCAAATTTAAAAATAACAGGTGAGTTGAATGGTATTGGGTTAGAGGGGACCTCGGTACCCC
TAGAGGAAACAGTGGAGACAGAGGTTTTTGTGGGACCACTTAAGACCTCTCTCTCTTTTGTCCTCAGTGATTCTGTTCCATGTAATCTGCTTGGCAAAGATGCTTTAATG
AAATTAAGGGCTAACATCCTGTACACAAAGGATGGCCCAATTGTTACTGCAAACAGTAGTTCAGAGGATGTTAATTTATTTGAAAGTRTTATTCCTCTGATGGCCACCAG
GGTCCATGGAAACTTGGAAATACCCCCCGATTTGGAGGGGCTTCCCAGTTCCCTATGGGCAACAAAGGGTGGAACTACAGGGTTAATACATAGGGCAGACCCTGT
CACTGTGCAGATAAGGCCGGGGGCCAAATTGCCCCGAGTGCCCCAATATCCTTTAAGCCAGAAGCAGATTAATGGAATAAGACCACAAATTACTTCCTTGTTGCAG
CAAGGCATTCTAATACCAGTCAAGTCACCCGTAAATACTCCCTTATATCCTGTCGCTAAGCCTGGTAAACAAGGTGAGTACCGGCTAGTGCAAGATCTACGGGCRGTA
AATAAGATAATTTTAGAGAACACACCAATTGTGCCCAACCCACATACAATATTAGGGAATATTCCACCTGAGGCTTCGTGGTTTTCAGTTTTTGATTTAGTAGATGCATACT
TTTGTGTTGCAATTAATGCTGAMTGTCAATACCTGTTTGCATTTACCTATGAAGGTGAGCAATATACTTGGGGGCGTCTCCCTCAGGGGATGGTATCCTCACCCTCAGA
GTTTGGACAGGTGATGAAACAGGTGTTGGACAGGTGGCAGCCTATACAGGGAACTACTGTGGTACAATATGTAGATGACTTGCTGCTTACAGCAATCTCAAAGGACTT
GTGCAGAGAGGCTTCCATTTCTCTGTTACAACACCTACATGAGGAAGGATGTAAAATATCACCTTCCAAGTTGCAGTACTGTCAACAGAAAGTCATCTTTTTGGGACATT
GTATCTCACGAGGTACAAAACACATTACCACAGATAGGATATTAATTATACAGAAAGCATCTTATCCACAAAGTGCTAAACAAGTCAGAGCATTCCTTGGTTTAGTTGGGT
ATTGTAGGCAGTGGATTCCTAATATGTCACAATTAGCCACTCCCTTATATGCTTTAACCTCACAGGCAGGACATATCATCCTCACTGAGGAACAAAAATCCTCAGTAGAC
CAGTTGAAGTTGGCAGTTCTAAGGGCCCCCTCCCTTGCACTTCCAAATTACAAAAAACCATTCTTTTTGTTTTGTCATGAGCAGGGAGGGCAAGCCTCAGGAGTACTG
ACACAGAAGGCTGGAGATAAGCAGAAGCCTGTGGGATATTTCAGCTGCAAGTTAGATGCTGTTATCAGCGCTGCAGCAGGGTGTGTGAGAGCAGTGGCAGCAGCA
GCTATGCTTGTAGAGAAAACTGCAGACATAGTACTGGGCAGTAAACTTTTTGTGATGGTTCCACACGCAGTACATACTGTACTGCAGAGCCTCACAACAAAACACCTC
TCGGCAGCCAGGCTCACTAAATATGAGGTTGGACTATTAGTCCCAGCTAACATCACTCTTCTCAGGTGCAATGTATTGAACCCAGCAACTCTGCTTCCTACGGAGCAC
CAAGGGGAAAGTGAAAATGATCTTGAGCATGACTGCCAACAGGTGATGGAACAATTTTATTCTCCTCCTAATCCAGTACAGGATGAACCAATACCAAATGCTGATCTTG
TTCTCTTTGTAGATGGTTCAAGATTGCAACATCCAGAGGGAGGGTTTGGTGCAGGGTTTGCTGTGGTCAGTGGCTCAGAGGTATTATACTCAGAGTCTCTTGACCCCAG
TCAACACTCAGCCCAAAGTGCCGAGCTGCTTGCTCTCATAAAGGCCTGTGAGTTAGCAAAAGGTCAAACAGCAAATATTTACACTGATTCAAGGTATGCCTATGGAGT
TATTCATGATTTTGCCCAACTGTGGAAACACAGGAATTTCCTTACTTCTAGTGGAAAGATGATAAAACATGCCACTTTAATTAAGGAATTGCTAGAAAAGGTGCAACTCCC
TAGGGATGTAGCTGTAATCAAGTGTCAGGCACATCAAAAGATAAAAGATGATATAACTGCAGGTAATGATAGGGCAGATAARGCCGCCAAAAAGGCTGCTTTTATGGTA
CTAACAGTAATACCAGCAGATGAGTTACCTACATCAGTTACCTTAGACATCCTTAAGAACCTGCAAAAGCAGGCTGGTACACAAATCCATACTAAGTGGATTAATAAGG
GGTGTAAGGTAGAGAATGATATTTGGAAACATGAGGATGGTCGTTTGTGTGCACCTCCCGTTTTATTTCCTTTGTTAGCAAATATCACGCACTTCCCATCTCATGTTTCCA
AAGGGGGAATGATATCCATGGTAAACAAACTTTGGTTTGCTCCTGGATTTGCRCAACACGCTACAGATTTCTGTAAGAAATGTATGATCTGTGCTAAACACAACACGGG
TAAAACAGAAAAGACACCTTTAAAACATTTGGTTAAACCATTTTATCCATTTCAGAGGTTACAGATTGATTATATTCAGTTACCTAAATGTAACACCTATGAATATGTATTGGT
TTGCATAGATATGTTTTCAGGGTGGATCGAGGCGTGGCCAGTAACAAAAGCCACAGCACTAATAACAGCTAAGAAATTGATAACTGAGATTGTCTGTCGCTATGGGCTC
CCAGAGACTATTGAATCTGACAGAGGTACACATTTTACAGGGCAAGTATTTGCAGAAATGCTAAAGGGGTTACAAATACACCAACACCTACACACCCCATATAGACCT
GAGGCTTCAGGTCGGGTGGAGAGAGCTAACGGTACCCTAAAAACAAAATTAGGTAAACTTTGTGAACAGACAAAGTTAACTTGGGTACAAGCTTTGCCCCTTGCACT
GTCATCTATGAGACACACTCCTAGGGGTCCCCATCAGTTATCTCCTTTTGAGGTACTTTTTGGCAGATCACCAAATACCGGATTATTTTTCCCACAGGAATTGCAGGGA
GAACATGCTTCCCTAACTGACTATGTTAAGCAATTGCATAAGCAGCTAACTAACCTGCATGGAAAAGTATTCTCTTCTTTACCAGATCCAGAGACTGTATTTGGAACTCA
CTCACTTCAACCAGGAGATTGGGTGGTGATAAAGAGATTCGTGAGGCGACACCTTGAGCCTAGGTACGACGGACCATATCAGGTTCTACTCACTACTGCCACCTCAA
TTAAAGTAGAAGGAAAGCCAAATTGGATTCAYGCCTCACACTGTAAGAAGGTATTTCCCCCATCAAGTCCTGCTGAGAGCCAGAGTAGAGTGCTATCTCCAGTGACAG
AGTCYGAATCTGTCCCTGCACTCTCACCCCCTCTAACGAGATCAAAGACCAGAGCACATCAACAGAAAGACAACTAAGGATCCAGACCTTCCGGAATCAAGAGGAT
AAAAGGCATCTTTAAAGAAAGTCAGACGAGAGTGGTAACAATAATCCCAMTAGTCTGTTAACCAACGGTTCCAGTTTAAGGTTTTGACACCAAGTGGTGGGCCTACTC
TGGACGAAGAGGATTATATCATCATCAAGAAGAAGGGACTTTGACAAAACAATGACTCAGTATTTGATGCTCTGGGTAAAGGTTCTTGTCTTTGTTGCACTATACTTTACW
CCAGTAAAGGGGGGTTGGCAAACAAATTCTATATGGCTTTTACACCAGCAGGCTGCAAAGCAGTTAAATACTACTGACTGCTGGATATGTTCACACGTTCCAGTTAATC
ATAAGGGGATACCTTTAGTAGGGATACCTATCTCTATGAATGATCTTAATCTTACAGATTATAGCTATGAAGTCGCAATAGATACAAACCATACGTATCAAAATGCAATTAAT
GATCAGGGAACATATTTRGAGATAACAGGATTACTAGATACACCCACTATATGTGTAGGACCCATGTTTGTTAATCACACCATTAAAATGAAAGATCACGTAGTAAATATAC
CTAMGATTCATGTAGGAAACACAGATTGTGAAAATGCCACTTTGAAATTCTACATGGATTATATGGGAAAGAAATGTGATAAGATGCAGGGTAATTGTAATGATTTTTGTTG
GTGTGGGAGAATGAATRCTTCTTGTGCCCCCAGGTGTTCTTGCCCTAATCAAGATATAGATGGAATTGTAAATTGTGAGTCTAAGGTGACAGATAATATGAGATGGTTTCA
GACTTGGCTGGGGTATCACGGTACAATCCTTCCTAGGGTACTCAGACAAGGACTATACTATATTTGTGGAAACAGAGCTTATTCATGGCTCCCTATGGGAGCATGGGG
AAATTGYACCATAGGAAGAGTTGTACCTGCCATTAGACAACGTCTGAACRTCTCATATATAGACAATCAAGGTATCCATTCCAGGGTAAATAAACAAAAAAGGGAATTATT
CTCCAACTCAGATAAGGCATGGATGTGGTTCCCTGCCTGGACAGGATGGGGAATAGAATTAGCCAATAGATTAAATAAGTATGCTAGTATAATGGATGGTATCATAAATG
AAACTACAAGTTCCATACACATAATAAATGAGGAACTAGCACAAGTTAGAAAGGTGACTTTACAGAATAGAATGGCTTTAGATTACCTACTAGCAGTCGAGGGTGGGGTT
TGTGCAATCATAGGTAAAGAGTGTTGTACCTGGATTGAAGATTCGCATGACCMAATTGAGGCCCACATGCATAAGGTAAAGGAATTACAGCAGAAAGCTAGAGATATCT
CCAAAGAAGGATGGAATCCTTTCTCTTGGATGGGAAATATAGGAACTTGGCTATCTAGTTTTTTAAAACTTTTATGGCACCCGTAGTTTGTGTGATTATAATATGTTTGATGTT
GTATGTAGTAGTAAGGTTGATAATGTGTTGCAGCAAAAACAGTATCAGGGGATAATAGACGACTTACAAAGGGAGGAGGTCTGTCTATTGGGCCCCTGAATAAGAAATAT
ACATTCTTCTTCTAGTCAGCGGCCCTCTGTGCTTTGGCCAACTTTATCCACAGAAGGATCCCATTCCTGTTTGAGTAGGATAAGGAAGAGACTGCTGATTTGGATGGGA
ACCCAGATAGGCAACTCTGTACTGCCACTGAACCTGCCATTAAGAAGTAGGTGGTGGAGGAGAGGTCTGGGTAAGAAGAGTRTATATATATATATGTCCAGGTATGAAC
AACAGGGAGGAATGTCAGAGAAAAAAGGAACCATCTTGTTTTCTGTACTCCATCTTGTTCATATCAATTAATTTAGGAGTGTTTTAAATATGTTTAAAAGTGTTGTTCTGTTAA
AACTGTGAATTGCTGATTTTGCATAATGGAGGGGCCTCCAACTAAAATAGCCCACTCCACACTTTGTACATGCGTCAGAGAGGAGCCTTAGTTCTATTGGCTATAGGGC
TCCTCCACCTTTCGTCTGTGTTCAAGAGATAACAAAGTATTGTGTAACTTTAACYGTATAAATGATGGTAGTTAAATGCTTAAACTTTGAGTAGCATGTTGTAGGGACAACGT
CAATGCGTCTGTCTGTCTACTCCTTTCACGTGAAATAAACCACTTCAATCAAACTGACCTATCCAACAGTTTTTCTTTGACAT

Figure S1: Consensus nucleotide sequences of AnERV families 1-10. Highlighted in red are LTRs, green is *gag* gene, purple is *pol* gene and blue is *env* gene. Where sequence is underlined, it is shared between two genes. AnERV1 and AnERV5 are represented by a single sequence.

Figure S2

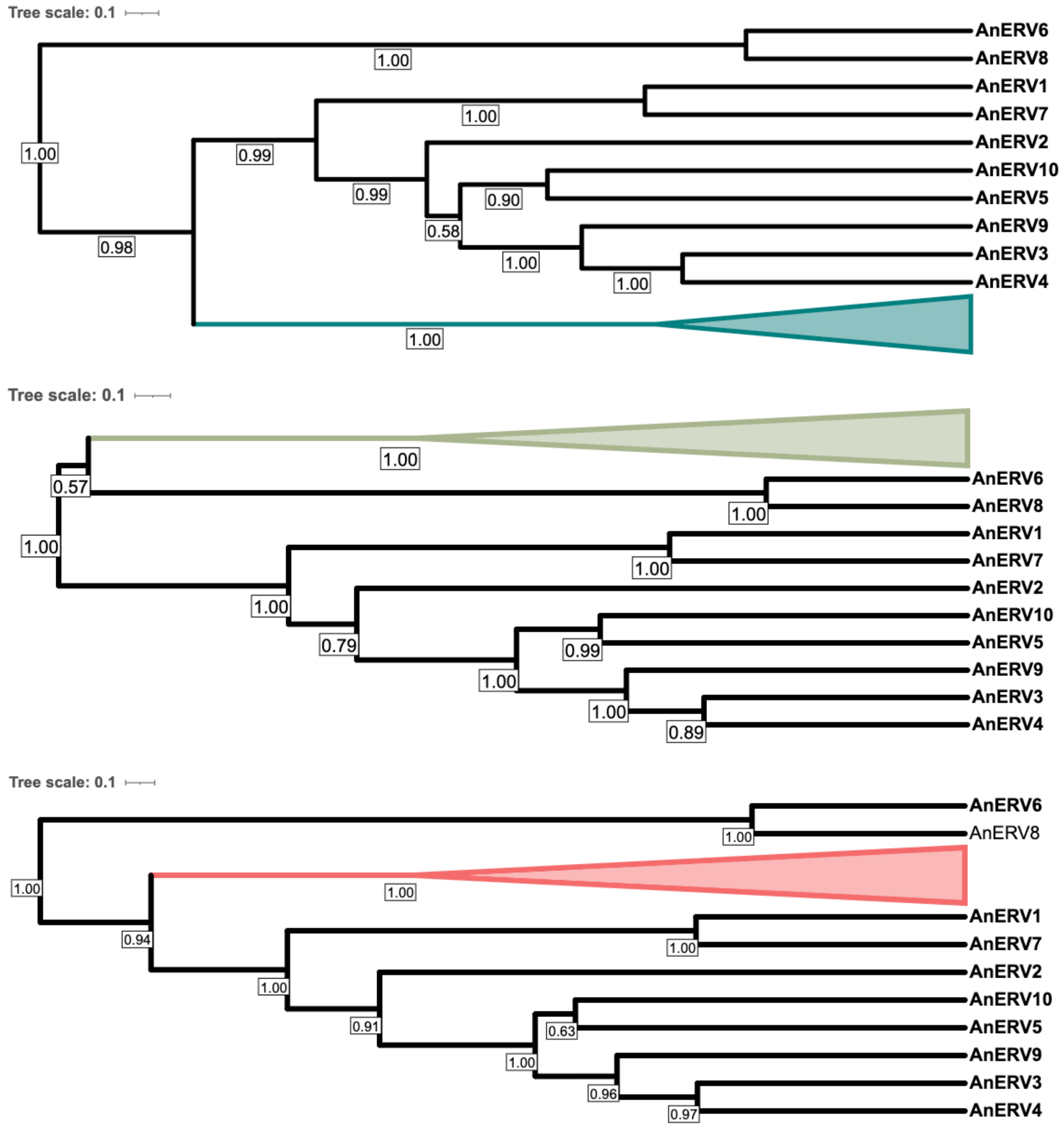


Figure S2: Bayesian phylogenetic trees of AnERVs identified in this study with representative retroviruses from alpharetrovirus (top, teal), deltaretrovirus (middle, green) and gammaretrovirus (bottom, pink). Posterior values >0.50 are displayed on nodes.

Table S1: List of Anuran genomes screened for alpharetrovirus ERVs.

| GenBank Accession | Family | Species | Genome size (Gb) | Assembly level |
|---|---|---|---|---|
| GCA_033576535.1 | Aromobatidae | *Allobates femoralis* | 5.3 | Contig |
| GCA_027579735.1 | Bombinatoridae | *Bombina bombina* | 10.000 | Chromosome |
| GCA_905336975.1 | Bombinatoridae | *Bombina variegata* | 4.679 | Scaffold |
| GCA_905171765.1 | Bufonidae | *Bufo bufo* | 5.045 | Chromosome |
| GCA_014858855.1 | Bufonidae | *Bufo gargarizans* | 4.545 | Chromosome |
| GCA_033119425.1 | Bufonidae | *Bufotes viridis* | 3.800 | Chromosome |
| GCA_027789725.1 | Hylidae | *Dendropsophus ebraccatus* | 2.400 | Scaffold |
| GCA_027410445.1 | Alytidae | *Discoglossus pictus* | 3.900 | Chromosome |
| GCA_019857665.1 | Eleutherodactylidae | *Eleutherodactylus coqui* | 2.789 | Chromosome |
| GCA_019512145.1 | Leptodactylidae | *Engystomops pustulosus* | 2.556 | Chromosome |
| GCA_027917425.1 | Microhylidae | *Gastrophryne carolinensis* | 4.300 | Chromosome |
| GCA_018402905.1 | Ranidae | *Glandirana rugosa* | 7.628 | Scaffold |
| GCA_029499605.1 | Hylidae | *Hyla sarda* | 4.100 | Chromosome |
| GCA_019447015.1 | Pipidae | *Hymenochirus boettgeri* | 3.211 | Chromosome |
| GCA_018994145.1 | Megophyridae | *Leptobrachium ailaonicum* | 3.536 | Chromosome |
| GCA_009667805.1 | Megophyridae | *Leptobrachium leishanense* | 3.549 | Chromosome |
| GCA_031893025.1 | Leptodactylidae | *Leptodactylus fuscus* | 2.300 | Chromosome |
| GCA_011038615.1 | Limnodynastidae | *Limnodynastes dumerilii* | 2.379 | Scaffold |
| GCA_002284835.2 | Ranidae | *Lithobates catesbeianus* | 6.250 | Scaffold |
| GCA_028564925.1 | Ranidae | *Lithobates sylvaticus* | 5.200 | Chromosome |
| GCA_000935625.1 | Dicroglossidae | *Nanorana parkeri* | 2.054 | Scaffold |
| GCA_009801035.1 | Dendrobatidae | *Oophaga pumilio* | 3.493 | Scaffold |
| GCA_033576555.1 | Dendrobatidae | *Oophaga sylvatica* | 5.200 | Scaffold |
| GCA_933207985.1 | Pelobates | *Pelobates cultripes* | 3.100 | Chromosome |
| GCA_022657655.1 | Dicroglossidae | *Phrynoglossus myanhessei* | 1.829 | Contig |
| GCA_025379985.1 | Hylidae | *Phyllomedusa bahiana* | 4.700 | Contig |
| GCA_021901965.1 | Pipidae | *Pipa carvalhoi* | 1.192 | Contig |
| GCA_019650415.1 | Pipidae | *Pipa parva* | 1.371 | Scaffold |
| GCA_016617825.1 | Limnodynastidae | *Platyplectrum ornatum* | 1.065 | Scaffold |
| GCA_023970735.1 | Hylidae | *Pseudis tocantins* | 0.024 | Scaffold |
| GCA_028390025.1 | Myobatrachidae | *Pseudophryne corroboree* | 8.900 | Chromosome |
| GCA_004786255.1 | Pyxicephalidae | *Pyxicephalus adspersus* | 1.563 | Chromosome |
| GCA_029574335.1 | Ranidae | *Rana kukunoris* | 4.800 | Chromosome |
| GCA_029206835.1 | Ranidae | *Rana muscosa* | 10.200 | Chromosome |

| | | | | |
|---|---|---|---|---|
| GCA_905171775.1 | Ranidae | *Rana temporaria* | 4.111 | Chromosome |
| GCA_032444005.1 | Dendrobatidae | *Ranitomeya imitator* | 6.000 | Chromosome |
| GCA_900303285.1 | Bufonidae | *Rhinella marina* | 2.552 | Contig |
| GCA_009364435.1 | Scaphiopodidae | *Scaphiopus couchii* | 0.484 | Scaffold |
| GCA_009364455.1 | Scaphiopodidae | *Scaphiopus holbrookii* | 0.711 | Scaffold |
| GCA_027358695.1 | Scaphiopodidae | *Spea bombifrons* | 0.989 | Chromosome |
| GCA_029215755.1 | Scaphiopodidae | *Spea hammondii* | 1.200 | Scaffold |
| GCA_009364415.1 | Scaphiopodidae | *Spea multiplicata* | 1.076 | Scaffold |
| GCA_951230385.1 | Ranidae | *Staurois parvus* | 4.000 | Scaffold |
| GCA_031769625.1 | Myobatrachidae | *Taudactylus pleione* | 5.500 | Contig |
| GCA_024363595.1 | Pipidae | *Xenopus borealis* | 2.747 | Chromosome |
| GCA_017654675.1 | Pipidae | *Xenopus laevis* | 2.742 | Chromosome |
| GCA_000004195.4 | Pipidae | *Xenopus tropicalis* | 1.451 | Chromosome |

Table S2: Table of the representative retroviruses used in the phylogenetic analysis.

| **Name** | **Genera** | **Host** | **GenBank Accession or Reference** |
|---|---|---|---|
| Avian leukemia virus | Alpha | Avian | NC_015116.1 |
| Avian leukosis virus | Alpha | Avian | KU375453.1 |
| Rous sarcoma virus | Alpha | Avian | J02342.1 |
| Avian endogenous retrovirus EAV-HP | Alpha | Avian | AJ238121.1 |
| ERV-AB.0-Mun | Beta | Amphibian | Chen, Guo and Zhang 2021 |
| Python molurus ERV | Beta | Reptile | AF500296.1 |
| Bovine retrovirus CH15 | Beta | Mammal | OM339153.1 |
| Ovine enzootic nasal tumour virus | Beta | Mammal | NC_007015.1 |
| Simian retrovirus 4 | Beta | Mammal | NC_014474.1 |
| Bovine leukemia virus | Delta | Mammal | LC760127.1 |
| Simian T-cell lymphotrophic virus 6 | Delta | Mammal | NC_011546.1 |
| Human T-lymphotropic virus 1 | Delta | Mammal | U19949.1 |
| Human T-lymphotropic virus 2 | Delta | Mammal | NC_001488.1 |
| Human T-lymphotropic virus 3 | Delta | Mammal | DQ093792.1 |
| Atlantic salmon swim bladder sarcoma virus | Epsilon | Fish | NC_007654.1 |
| Zebrafish ERV | Epsilon | Fish | AF503912 |
| Rhinella marina endogenous retrovirus | Epsilon | Amphibian | MG981046 |
| Xenopus laevis ERV 1 | Epsilon | Amphibian | AJ506107.1 |
| Walleye dermal sarcoma virus | Epsilon | Fish | NC_001867.1 |
| Walleye epidermal hyperplasia virus type 1 | Epsilon | Fish | AF133051.1 |
| Walleye epidermal hyperplasia virus type 2 | Epsilon | Fish | AF133052.1 |
| ERV-EA.a-Amex | Epsilon | Amphibian | Chen et al, 2022 |

| | | | |
|---|---|---|---|
| ERV-EA.a-Bga | Epsilon | Amphibian | Chen et al, 2022 |
| ERV-EA.c-Bga | Epsilon | Amphibian | Chen et al, 2022 |
| ERV-GA.b-Lle | Gamma | Amphibian | Chen et al, 2022 |
| ERV-GA.a-Mun | Gamma | Amphibian | Chen et al, 2022 |
| Galidia ERV | Gamma | Mammal | KF313135.1 |
| Duck infectious anemia virus | Gamma | Avian | KF313137.1 |
| Reticuloendotheliosis virus strain HA9901 | Gamma | Avian | NC_006934.1 |
| Reticuloendotheliosis virus strain 104865 | Gamma | Avian | OL857288.2 |
| Baboon endogenous virus | Gamma | Mammal | NC_022517.1 |
| RD114 retrovirus | Gamma | Mammal | NC_009889.1 |
| Porcine endogenous retrovirus | Gamma | Mammal | AJ293656.1 |
| Koala retrovirus | Gamma | Mammal | KC779547.1 |
| Gibbon ape leukemia virus | Gamma | Mammal | NC_001885.3 |
| Woolly monkey sarcoma virus | Gamma | Mammal | KT724051.1 |
| Feline leukemia virus | Gamma | Mammal | NC_001940.1 |
| Murine Leukaemia virus | Gamma | Mammal | AY818896.1 |
| Xenotrophic MuLV-related virus | Gamma | Mammal | FR872816.1 |
| Feline leukemia virus | Gamma | Mammal | NC_001940 |
| Rauscher murine leukemia virus | Gamma | Mammal | U94692.1 |
| Mus musculus ERV-L | Class III | Mammal | Y12713.1 |
| Rattus novergicus ERV-L | Class III | Mammal | AC127142: 48521-54855; Yedavalli et al, 2021 |
| Xenopus tropicalis ERV-S | Class III | Amphibian | MW779451.1 |
| White-tufted-ear marmoset simian foamy virus | Class III | Mammal | GU356395.1 |
| Squirrel monkey foamy virus | Class III | Mammal | GU356394.1 |
| Brown greater galago prosimian foamy virus | Class III | Mammal | NC_039023.1 |
| Puma feline foamy virus | Class III | Mammal | NC_039022.1 |
| Western chimpanzee simian foamy virus | Class III | Mammal | U04327.1 |
| Eastern chimpanzee simian foamy virus | Class III | Mammal | NC_039025.1 |
| Human_Spumaretrovirus | Class III | Mammal | U21247.1 |
| Guenon simian foamy virus | Class III | Mammal | NC_043445.1 |
| Japanese macaque simian foamy virus | Class III | Mammal | NC_039026.1 |
| Rhesus macaque simian foamy virus | Class III | Mammal | NC_039238.1 |
| Taiwanese macaque simian foamy virus | Class III | Mammal | MN585198.1 |

Table S3: Potential full-length ERV elements (LTR pairs <20kb distance and >95% similarity with *env* tBLASTn hit between) organised into families (A-J) in each species genome. Segmental duplication is indicated where alignment continued 1kb upstream and downstream of element. Family I in *Bufotes viridis* (*) has *env* ORF >1000 but untypical genomic organisation and so was not included in *env* comparison between species.

| **Species** | **Insertion ID** | **Number of loci** | **Alignment consensus** | | | **Segmental duplication** |
|---|---|---|---|---|---|---|
| | | | **gag ORF** | **pol ORF** | **env ORF** | |
| *Spea multiplicata* | | 2 | none | <1000 | >1000 | |
| | AnERV1 | 1 | >1000 | >1000 | >1000 | |
| | AnERV2 | 1 | >1000 | >1000 | >1000 | |
| *Spea bombifrons* | | 3 | none | >1000 | >1000 | |
| | AnERV2 | 5 | fragmented | <1000 | >1000 | |
| *Spea hammondii* | | 1 | >1000 | fragmented | fragmented | |
| | | 7 | >1000 | fragmented | >1000 | |
| | AnERV2 | 7 | >1000 | <1000 | <1000 | |
| | | 2 | fragmented | fragmented | fragmented | |
| *Leptobrachium leishanense* | | 62 | fragmented | >1000 | fragmented | |
| | | 13 | <1000 | fragmented | >1000 | |
| *Leptobrachium ailaonicum* | | 12 | fragmented | fragmented | >1000 | |
| | | 1 | >1000 | fragmented | >1000 | |
| | | 3 | fragmented | fragmented | <1000 | |
| | | 2 | >1000 | fragmented | >1000 | |
| | | 16 | fragmented | >1000 | >1000 | |
| *Pelobates cultripes* | | 8 | >1000 | fragmented | <1000 | |
| | | 2 | <1000 | fragmented | <1000 | |
| | AnERV3 | 18 | >1000 | >1000 | >1000 | |
| *Rhinella marina* | | 1 | >1000 | fragmented | >1000 | |
| *Bufo bufo* | AnERV4 | 2 | fragmented | fragmented | >1000 | |
| | | 4 | >1000 | fragmented | <1000 | |
| | | 21 | none | <1000 | <1000 | |
| | | 64 | fragmented | fragmented | fragmented | |
| | | 17 | none | fragmented | >1000 | |
| | | 15 | <1000 | fragmented | fragmented | |
| | | 45 | none | fragmented | fragmented | |
| | | 47 | none | fragmented | fragmented | |
| | | 8 | none | fragmented | >1000 | Yes |
| | | 121 | fragmented | fragmented | fragmented | |
| | | 118 | none | fragmented | <1000 | |
| | | 2 | none | fragmented | fragmented | |
| | | 2 | none | none | fragmented | Yes |
| *Bufo gargarizans* | AnERV4 | 26 | >1000 | >1000 | >1000 | |
| | | 3 | none | >1000 | >1000 | |
| | | 59 | none | fragmented | fragmented | |
| | | 4 | none | fragmented | <1000 | |
| | | 3 | <1000 | fragmented | fragmented | |
| | | 6 | fragmented | fragmented | fragmented | |
| *Bufotes viridis* | | 70 | fragmented | fragmented | fragmented | |
| | | 10 | fragmented | fragmented | >1000 | |

| | | | | | | |
|---|---|---|---|---|---|---|
| | | 40 | <1000 | fragmented | fragmented | |
| | | 10 | none | none | fragmented | |
| | | 3 | none | fragmented | fragmented | Yes |
| | | 3 | none | <1000 | <1000 | Yes |
| | | 2 | fragmented | fragmented | fragmented | |
| | | 1 | fragmented | fragmented | <1000 | |
| | | 1 | >1000 | fragmented | >1000 | |
| | | 2 | fragmented | fragmented | fragmented | |
| *Engystomops pustulosus* | | 2 | <1000 | fragmented | <1000 | |
| *Dendropsophus ebraccatus* | | 5 | <1000 | <1000 | >1000 | |
| | | 4 | none | none | <1000 | Yes |
| | | 2 | fragmented | >1000 | >1000 | |
| *Eleutherodactylus coqui* | | 3 | none | fragmented | <1000 | |
| | | 1 | none | fragmented | <1000 | |
| | | 2 | none | fragmented | fragmented | |
| *Taudactylus pleione* | | 1 | none | fragmented | <1000 | |
| | | 2 | none | fragmented | >1000 | |
| *Pyxicephalus adspersus* | | 8 | <1000 | fragmented | fragmented | |
| | | 5 | fragmented | fragmented | >1000 | |
| *Lithobates catesbeianus* | | 1 | none | fragmented | <1000 | |
| *Lithobates sylvaticus* | AnERV5 | 7 | >1000 | >1000 | >1000 | |
| *Glandirana rugosa* | | 2 | <1000 | fragmented | >1000 | |
| | AnERV5 | 1 | >1000 | >1000 | >1000 | |
| | | 1 | <1000 | >1000 | >1000 | |
| *Gastrophryne carolinensis* | AnERV9 | 18 | >1000 | >1000 | >1000 | |
| | | 38 | none | fragmented | >1000 | |
| | | 1 | fragmented | fragmented | fragmented | |
| | | 1 | <1000 | fragmented | >1000 | |
| | AnERV6 | 4 | >1000 | >1000 | >1000 | |
| *Hymenochirus boettgeri* | | 25 | none | fragmented | <1000 | |
| *Xenopus laevis* | | 4 | <1000 | fragmented | fragmented | |
| *Xenopus tropicalis* | | 12 | <1000 | fragmented | >1000 | |
| | | 2 | <1000 | fragmented | >1000 | |
| | AnERV10 | 2 | >1000 | >1000 | >1000 | |
| *Bombina bombina* | AnERV7 | 23 | >1000 | >1000 | >1000 | |
| | | 12 | >1000 | >1000 | >1000 | |
| | | 2 | none | fragmented | >1000 | Yes |
| | | 39 | none | <1000 | >1000 | |
| | AnERV8 | 12 | >1000 | >1000 | >1000 | |
| *Discoglossus pictus* | | 4 | <1000 | fragmented | >1000 | |
| | | 9 | none | fragmented | <1000 | |
| | | 7 | none | fragmented | >1000 | |
